# Understanding Supramolecular Assembly of Supercharged Proteins

**DOI:** 10.1101/2022.06.21.497010

**Authors:** Michael I. Jacobs, Prateek Bansal, Diwakar Shukla, Charles M. Schroeder

## Abstract

Ordered supramolecular assemblies of supercharged synthetic proteins have recently been created using electrostatic interactions between oppositely charged proteins. Despite recent progress, the fundamental mechanisms governing the assembly process between oppositely supercharged proteins are not fully understood. In this work, we use a combination of experiments and computational modeling to systematically study the supramolecular assembly process for a series of oppositely supercharged green fluorescent protein (GFP) variants. Our results show that the assembled structures of oppositely supercharged proteins critically depend on surface charge distributions. In addition, net charge is a sufficient molecular descriptor to predict the interaction fate of oppositely charged proteins under a given set of solution conditions (e.g., ionic strength). Interestingly, our results show that a large excess of charge is necessary to nucleate assembly and that charged residues that are not directly involved in interprotein interactions contribute to a substantial fraction (∼30%) of the interaction energy between oppositely charged proteins via long-range electrostatic interactions. Dynamic subunit exchange experiments enabled by Förster resonance energy transfer (FRET) further show that relatively small, 16-subunit assemblies of oppositely charged proteins have kinetic lifetimes on the order of ∼10-40 minutes, which is governed by protein composition and solution conditions. Overall, our work shows that a balance between kinetic stability and electrostatic charge ultimately determine the fate of supramolecular assemblies of supercharged proteins. Broadly, our results inform how protein supercharging can be used to generate different ordered supramolecular assemblies from a single parent protein building block.

## Introduction

The assembly of biologically encoded molecules into ordered supramolecular structures gives rise to unique properties in natural biological systems, such as increased stability and complex functionality of macromolecular assemblies in cells.^1–5^ Inspired by nature, researchers have used biomolecular assembly strategies to develop several new biotechnologies for a range of applications including drug delivery, sensors, and facilitated charge transport.^6–9^ In recent years, significant experimental and computational efforts have been directed at understanding the structure-function relationships of new materials constructed from assembled biological building blocks.^10^

Supramolecular assembly of proteins can be driven by several different strategies including receptor-ligand, metal-ligand, electrostatic, and hydrophobic interactions.^11^ Using *de novo* engineering of synthetic proteins, different assembled protein architectures have been created,^12^ including one-dimensional (1D) nanowires,^13–15^ nanorings and nanotubes,^16–19^ and 2D and 3D protein crystals.^14,20–24^ However, protein engineering strategies for supramolecular assembly generally require precise and targeted design of complex molecular interfaces due to the highly specific nature of the governing intermolecular interactions. Given the complexity of biomolecular interfaces and interactions, many supramolecular assembly strategies are not easily generalizable across different classes of proteins.

Electrostatic interactions between oppositely charged proteins are known to drive hierarchical assembly of both folded and unfolded proteins.^25^ Electrostatic interactions between folded proteins have been used to create binary protein crystals from highly-symmetric cage proteins^26–30^ as well as to localize charged proteins in Matryoshka-like cages^31^ and protein capsules.^32–35^ Supramolecular assembly of proteins via electrostatic interactions can be extended to initially uncharged proteins by supercharging protein surfaces to include charged residues.^36,37^ Recent work has shown that oppositely supercharged green fluorescent protein (GFP) variants readily assemble into an organized 16-subunit protomer, suggesting that highly symmetrical building blocks may not be necessary to drive hierarchical assembly into ordered structures.^38^ Although electrostatic interactions provide a promising method of protein supramolecular assembly, we lack a complete understanding of the underlying assembly process and the molecular design rules governing electrostatic-mediated protein-protein interactions.

Complexities in supercharged protein assembly arise due to the nature of multi-charge and multi-body interactions, which includes the role of surface charge distributions. In general, protein supercharging does not yield a uniform surface charge distribution, but rather incorporates localized charged regions dispersed over the entire protein surface. Prior computational and experimental studies on spherical and polyhedral colloidal particles have shown that both shape complementarity^39^ as well as patchy attractive interactions^40,41^ give rise to symmetric hierarchical assembly. By changing the distribution of attractive patches on the surface of a protein, it is thought that proteins can assemble into a variety of different quaternary structures. For example, ferritin cage proteins are known to assemble into different hierarchical structures by changing the location of the attractive patches or the assembly conditions.^20,22,23,30,42^ In addition, it was reported that changing the distribution of charges on protein surfaces affects their ability to form complex coacervates with strong polyelectrolytes.^43,44^ We therefore hypothesized that different hierarchical assembled structures can be obtained from a single parent protein building block by changing the net charge and surface charge distributions on oppositely supercharged protein partner pairs.

In this work, we use a combination of experiments and computational modeling to understand the assembly mechanisms of supercharged proteins. Using a series of oppositely supercharged proteins (**Figure 1**), we explore how net charge and surface charge distributions on a common parent protein building block (GFP) affect intermolecular interactions and assembled structures. We further demonstrate that supercharged proteins with similar net charges but different surface charge distributions assemble into a variety of different structures with distinct properties and kinetic stability. Overall, our work provides an improved understanding of how proteins assemble via electrostatically-driven assembly pathways for oppositely charged proteins.

**Figure 1.**
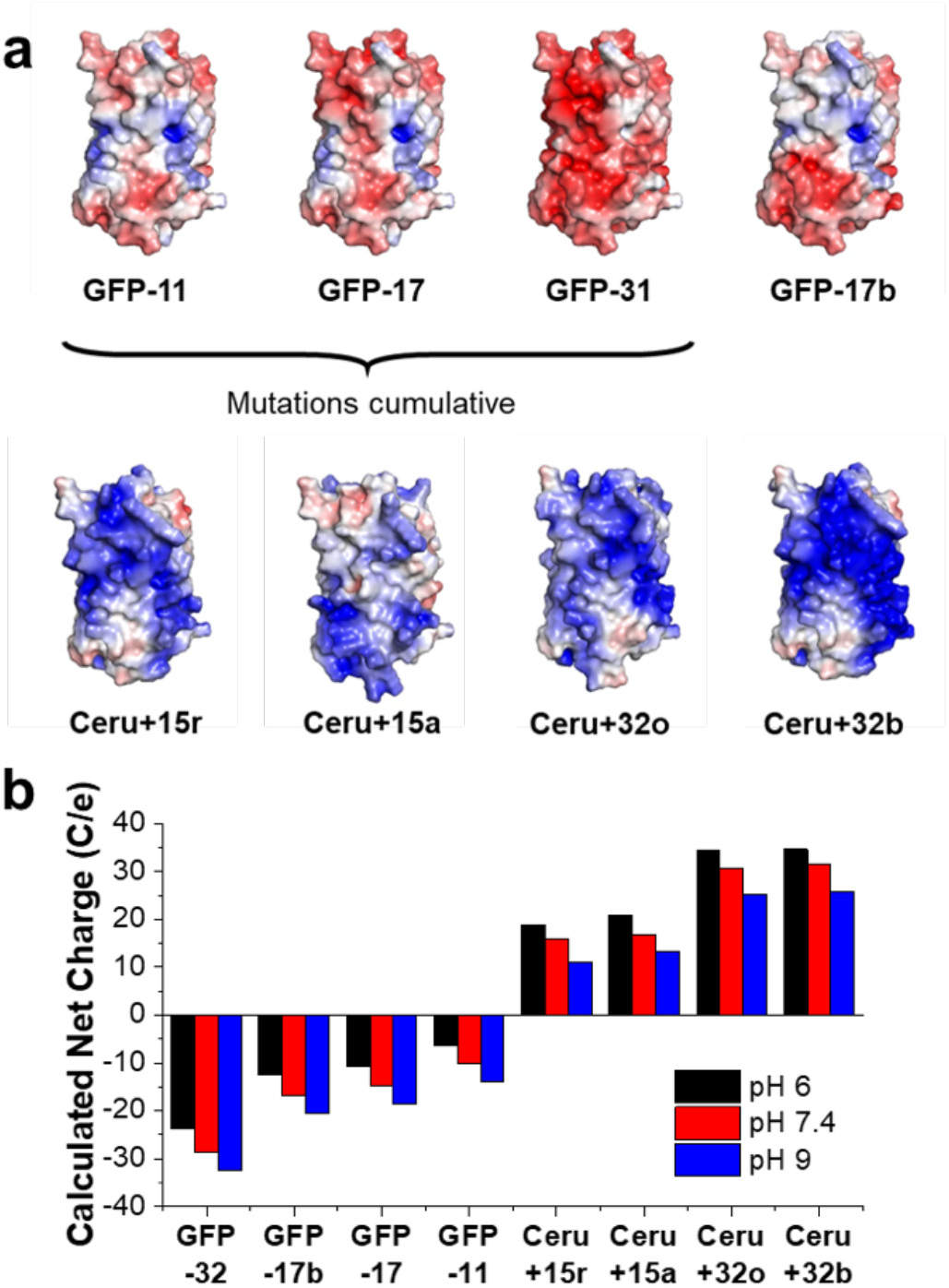
Supercharged GFP variants considered in this work. (a) Electrostatic surface potential representations of the solvent accessible surface area for GFP variants calculated using the linearized Poisson-Boltzmann equation with Adaptive Poisson-Boltzmann Solver (ABPS) (±10 k_b_T/e; blue for positive and red for negative). (b) Nominal net charge of the GFP and Ceru variants calculated using PROPKA at each of the pH values used here.^58^

## Results and Discussion

### Non-interacting charged residues affect supercharged protein assembly

We began by studying the role of non-interacting charged residues in supramolecular protein assemblies. Prior work reported the structure of an assembled 16-mer protein (two stacked octamers) composed of oppositely supercharged GFPs (denoted as GFP-17 and Ceru+32o in this work) solved at 3.47 Å resolution using cryo-electron microscopy.^38^ At this spatial resolution, 176 specific inter-protein interactions (e.g., hydrogen bonds and salt bridges) were identified as stabilizing the structure. Although 126 (72%) of the interactions involved a mutated charged amino acid, only 29% of the mutated charged amino acids were involved in stabilizing inter-protein interactions in the ordered assembly.^38^ A natural question then arises: what is the role of non-interacting, outward-facing mutated charged residues in promoting supercharged assembly?

To understand the role of non-interacting charged residues in supramolecular assembly of supercharged proteins, we designed and expressed two minimally mutated GFP and Cerulean variants that only contain the mutations participating in inter-protein interactions. The minimally mutated variant of GFP-17 (named GFP-min) has 4 of the original 11 mutations and a nominal net charge of -8 at pH 7.4. The minimally mutated variant of Ceru+32o (named Ceru-min) has 6 of the original 24 mutations and a nominal net charge of +3 at pH 7.4. Clearly, the minimally mutated variants have significantly smaller net charges than the fully supercharged variants, but they retain the requisite set of mutated amino acids involved in stabilizing the inter-protein interfaces in the 16-mer assembled structure.

We began by determining the Förster resonance energy transfer (FRET) efficiency between the fully supercharged-proteins (Ceru+32o / GFP-17) and minimally mutated protein pairs (Ceru-min / GFP-min) as a function of ionic strength (**Figure 2a**). As ionic strength increases, FRET efficiency between GFP-17 and Ceru+32o decreases as the electrostatic interactions between the proteins are more effectively screened. Based on the spectral overlap between unmodified Cerulean^45^ and superfolder GFP,^46^ the Förster distance *R*_*o*_ for the protein pair is calculated to be 5.5 nm.^47^ Thus, for FRET efficiency > 0.5, the average spacing between Ceru+32o and GFP-17 is expected to be less than approximately 5.5 nm. Our results show that the FRET efficiencies between Ceru-min and GFP-min are significantly lower than those between the fully supercharged variants at all NaCl concentrations, which suggests the minimally mutated variants do not associate as strongly as the supercharged protein pairs.

**Figure 2.**
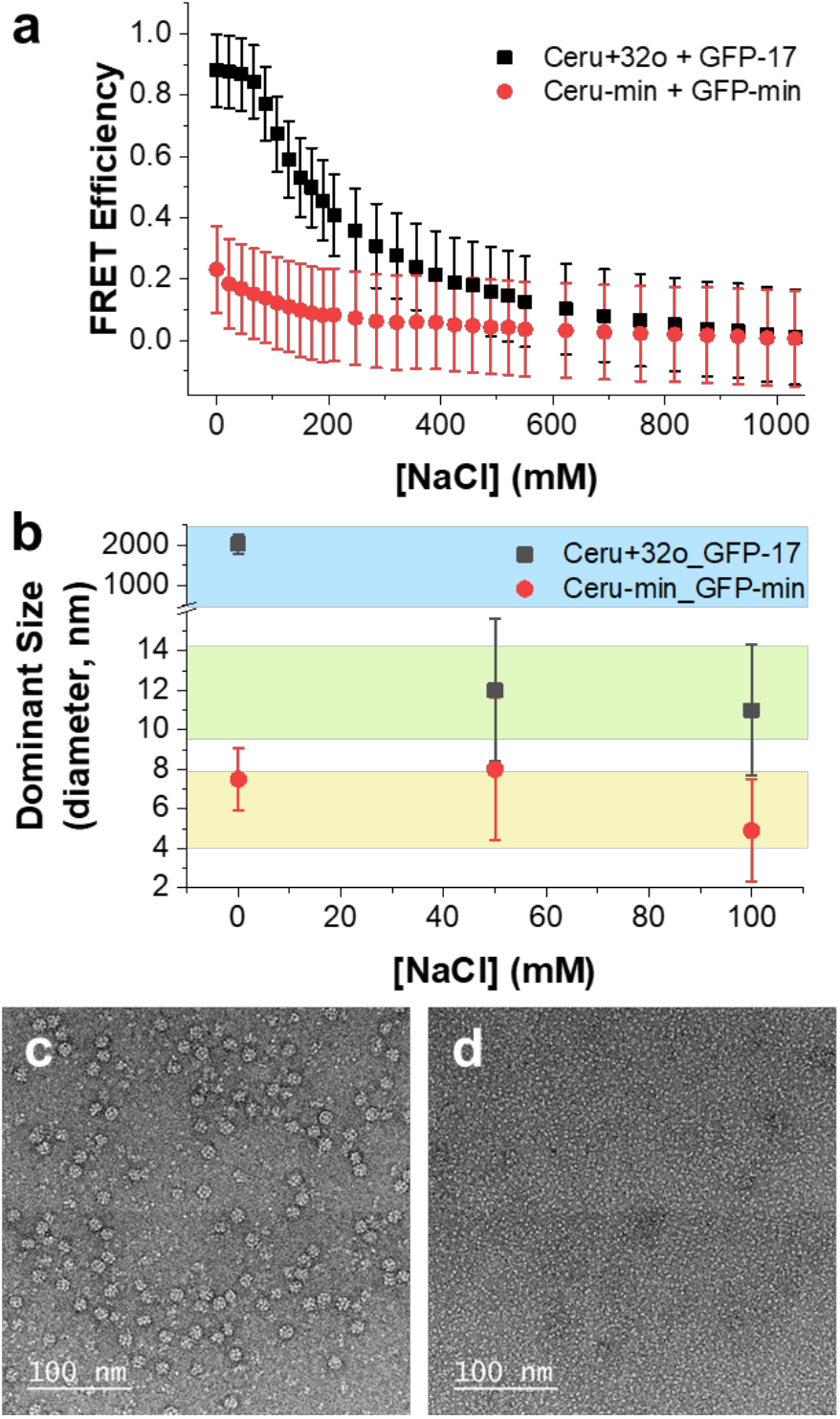
Characterization of fully charged and minimally charged GFP mutants. (a) FRET efficiencies between fully charged Ceru+32o / GFP-17 (black squares) and minimally mutated Ceru-min / GFP-min (red squares) at different salt concentrations. (b) Dominant assembly size as measured by DLS for fully charged and minimally mutated variants at different concentrations of NaCl. (c) Representative negative stain TEM image of assemblies from fully charged Ceru+32o / GFP-17 at 75 mM NaCl showing protomer formation. (d) Representative negative stain TEM image of minimally mutated Ceru-min / GFP-min at 0 mM NaCl showing no assembly or interactions between the proteins.

Dynamic light scattering (DLS) was used to determine the average size (hydrodynamic radius) of assembled protein structures as a function of ionic strength (**Figure 2b**). As previously reported for assembly between the fully supercharged proteins (Ceru+32o / GFP-17),^38^ large aggregates (>1,000 nm in diameter) form at low salt concentration. As salt concentration increases, intermediate particle sizes (∼12 nm) are observed, indicating that the proteins assemble into a protomer structure. Compared to the fully supercharged variants, assemblies formed from the minimally mutated variants (Ceru-min / GFP-min) have average sizes matching the monomeric species even at low salt concentrations. For example, the average diameter of particles at 0 M NaCl is 7.5 nm, which is not significantly different than the monomeric species (approximately 5-7 nm in diameter). These results are consistent with FRET measurements and suggest that Ceru-min and GFP-min do not strongly associate at the range of salt concentrations considered here.

Transmission electron microscopy (TEM) was further used to directly image protein assemblies formed from oppositely charged GFP variants. **Figure 2c** shows a TEM micrograph of a negatively stained protein assembly between Ceru+32o and GFP-17 prepared at 75 mM NaCl. An octomeric ring structure is clearly present, suggesting that the proteins assemble into an ordered 16-mer, as previously reported.^38^ However, only unassembled proteins are observed between Ceru-min and GFP-min even with no added salt (**Figure 2d**). Thus, our results show that the minimally mutated variants are incapable of forming the ordered 16-mer structure, and the externally facing, non-interacting mutated charged amino acids are necessary to promote assembly.

### MD simulations and binding energy of charged protein dimers

Based on the behavior of the minimally mutated variants, we hypothesized that the non-interacting mutated charged amino acids promote assembly by stabilizing long-range electrostatic interactions within the assembled structure and are therefore required to nucleate protomer formation. To test the validity of this hypothesis, atomistic MD simulations were performed to determine the standard free energy of binding (*ΔG*^*0*^_*bind*_) between supercharged protein dimers and minimally mutated protein dimers. The assembled 16-mer is composed of four distinct protein-protein interfaces:^38^ two intra-planar interfaces (named ‘Ceru+32 Clockwise’ and ‘GFP-17 Clockwise’, **Figure 3a**) and two inter-planar interfaces (named ‘Inter-GFP-17’ and ‘Inter-ring’, **Figure 3b**), where the plane is the toroidal plane of the 16-mer, normal to the axis of the torus. **Figures 3c-f** show the dimeric systems that were investigated: dimers with the fully supercharged and minimally mutated proteins oriented to obtain the four primary and distinct protein-protein interfaces in the 16-mer. *ΔG*^*0*^_*bind*_ quantifies the free energy released during the association process and depends on the protein-protein interface, number of contacts, and separation distance between the protomers.

**Figure 3.**
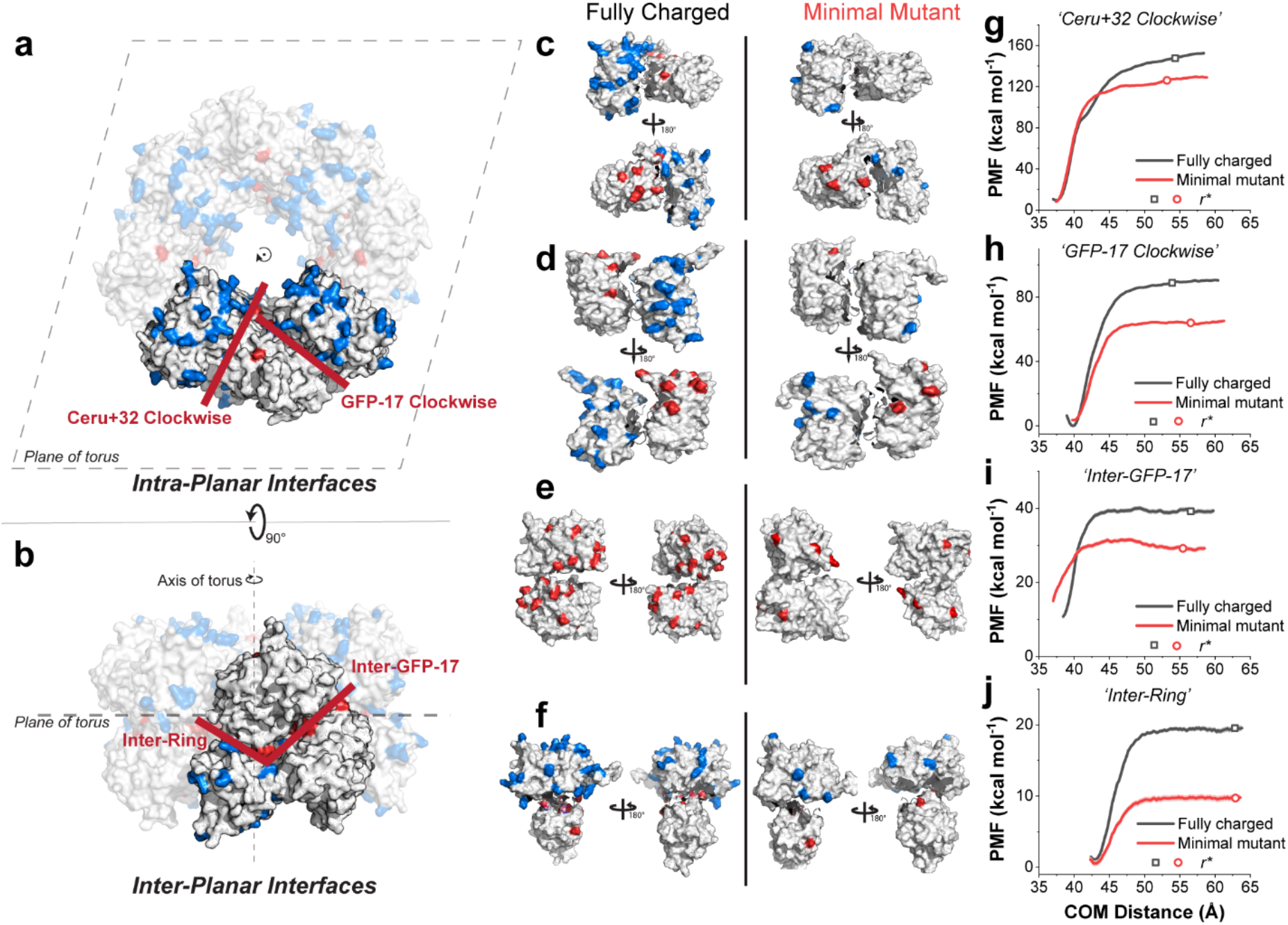
Binding free energy calculations for the protomeric interfaces for the fully charged and minimally mutated proteins. The four primary protein-protein interfaces comprising the assembled 16-mer identified by Simon et al.^38^ are shown. (a) ‘Ceru+32 Clockwise’ and ‘GFP-17 Clockwise’ are defined as intra-planar interfaces, and (b) ‘Inter-GFP-17’ and ‘Inter-Ring’ are defined as cross-planar interfaces, with the plane being the toroidal plane of the 16-mer, normal to the axis of the torus. The positively- and negatively-charged amino acid mutations are shown in blue and red, respectively. Dimer systems that represent each of the four interfaces are shown in (c) ‘Ceru+32 Clockwise’, (d) ‘GFP-17 Clockwise’, (e) ‘Inter-GFP-17’, and (f) ‘Inter-Ring’. The separation PMF values as a function of center-of-mass (COM) distance are shown for each of the interfaces (g-j).

MD simulations were performed using a previously reported method to determine the absolute binding free energies using biased simulations (**Figure 3** and **Table 1**).^48–53^ To calculate *ΔG*^*0*^_*bind*_, two proteins in each system are first separated from their initial bound pose until their respective center of masses (COM) had moved by at least 25 Å and the separation potential of mean force (PMF) has plateaued, indicating that the proteins are unbound and non-interacting (**Figures 3g-j**). The separation PMFs for each dimeric interaction follow a similar trajectory as the separation distance increases, such that a nearly linear increase from the equilibrium position to a plateau value is observed. The PMF is determined as a product of the pulling force and separation distance, which allows for a threshold separation *r\** to be defined where the contributions of inter-protein, non-bonded interactions taper out (i.e., only long-range electrostatic interactions remain), and the force required to pull the proteins apart decreases. At separation distances larger than *r\**, the PMF is observed to remain nearly constant. Separation simulations were performed under various conformational, orientational, and positional constraints using steered MD, and the contributions of the constraints to *ΔG*^*0*^_*bind*_ were explored using umbrella sampling and adaptive biasing force (ABF) methods (Supporting Information). In all cases, the separation PMF is found to be the largest contributor to *ΔG*^*0*^_*bind*_.

**Table 1.** Computed *ΔG*^*0*^_*bind*_ values for each interface.

| Interface | Fully charged<br>$\Delta G^0_{bind}$ (kcal mol<sup>-1</sup>) | Minimally mutated<br>$\Delta G^0_{bind}$ (kcal mol<sup>-1</sup>) |
| --- | --- | --- |
| Ceru+32 Clockwise | -84±4 | -56±3 |
| GFP-17 Clockwise | -52±1 | -34±2 |
| Inter-GFP-17 | -13.4±0.4 | -7.1±0.7 |
| Inter-Ring | -7.5±0.6 | -5.6±0.6 |

The values of *ΔG*^*0*^_*bind*_ and separation PMFs for eight different protein dimer systems show several interesting trends. First, interfaces with more contacts in the bound state have larger values of *ΔG*^*0*^_*bind*_ and separation PMF (**Figure S1**). For example, ‘Ceru+32 Clockwise’, an intra-planar interface with 8+ interprotein contacts, has the highest PMF at *r\** and the largest *ΔG*^*0*^_*bind*_ (−84 ± 4 kcal mol^-1^). This is followed by the intra-planar interface ‘GFP-17 Clockwise’ (6+ interprotein contacts, *ΔG*^*0*^_*bind*_ = -52 ± 1 kcal mol^-1^), inter-planar Inter-GFP-17 (3+ interprotein contacts, *ΔG*^*0*^_*bind*_ = -13.4 ± 0.4 kcal mol^-1^), and finally inter-planar Inter-Ring (2 interprotein contacts, *ΔG*^*0*^_*bind*_ = -7.5 ± 0.6 kcal mol^-1^). These results suggest that the intra-planar interfaces dominate the free-energy of binding in the assembled 16-mer structure, and mutations that disrupt these interactions are expected to inhibit supramolecular assembly.

Our results show that the values of *ΔG*^*0*^_*bind*_ for the intra-planar interfaces are significantly larger than previously reported values for protein-protein interfaces (which are typically 10-20 kcal mol^-1^).^54^ This is consistent with the notion that interactions between supercharged protein mutants arise due to strong electrostatic interactions that are absent in parent proteins. However, *ΔG*^*0*^_*bind*_ is also a function of the size of the interface,^55,56^ so equivalently large values of *ΔG*^*0*^_*bind*_ are expected for larger endogenous protein-protein interfaces. Nevertheless, the *ΔG*^*0*^_*bind*_ values for interactions between minimally charged proteins are consistently 20-30% smaller than those between the fully charged proteins. MD simulations were further used to understand the contribution of each constraint imposed on the proteins during separation to the calculated *ΔG*^*0*^_*bind*_ values. Interestingly, we find that the equilibrium values for the conformational, positional, and orientational constraints are similar (typically ±5%) for the fully charged and minimally mutated systems (**Figures S2-S5** and **Tables S1-S4**). Therefore, these results suggest that the fully charged and minimally charged protein variants preferentially adopt similar equilibrium bound configurations. Moreover, the difference in *ΔG*^*0*^_*bind*_ between the fully charged and minimally charged proteins is attributed to the absence of long-range electrostatic interactions in the minimally mutated variants. Due to the logarithmic nature of free energy calculations (Δ*G*_*bind*_ = −*k*_*b*_*T* ln *K*_*bind*_), small changes in *ΔG*^*0*^_*bind*_ can have dramatic effects on the equilibrium binding constant *K*_*bind*_. For example, the *K*_*bind*_ values for the fully charged ‘Ceru+32 Clockwise’ interface is a factor of ∼10^12^ larger than that of the minimally mutated protein interface, which suggests that long-range electrostatic interactions significantly increase the probability of forming an interface and promoting assembly.

Overall, the process of protein association is highly non-specific, and a continuum of metastable states exist in which the transiently formed interfaces do not lead to the final interface observed in the assembled structures. Such metastable states are accessible in the absence of a dominant free-energy minima that overpowers the preference for a single interface. We posit that these metastable states become more accessible in the minimally mutated variants, and hence result in non-specific binding in minimally charged protein dimers. Overall, these results suggest that identification of specific protein-protein contacts from high-resolution protein structures alone is insufficient to fully understand the interactions between supercharged proteins.

### Net charge on supercharged proteins predicts protein-protein interactions

We next studied the role of net charge and surface charge distribution on the assembly of supercharged proteins. Here, we designed and expressed a series of positively and negatively charged GFP variants (**Figure 1**). Differences in surface charge distributions are visualized from electrostatic surface potential representations (**Figure 1a**).^57^ At pH 7.4, the negatively charged GFP variants had nominal net charges of -11, -17 (two variants: GFP-17 and GFP-17b), and -32, and the positively charged Ceru variants had nominal net charges of +15 (two variants: Ceru+15r and Ceru+15a) and +32 (two variants: Ceru+32o and Ceru+32b). For variants with the same net charge, different sets of mutations were used to generate different surface charge distributions. By varying the solution pH, the net charges on the proteins (and surface charge distributions) were varied to produce more negatively charged (lower pH) or positively charged (higher pH) monomeric proteins (**Figure 1b**).^57,58^

**Figure 4a** shows FRET efficiency when Ceru+32o is mixed in equimolar amounts with each of the negatively charged GFP variants. For all mixtures, the FRET efficiency decreases with increasing NaCl concentration, indicating that the average distance between Ceru+32o and the negatively charged GFP variants increases with increasing salt concentration. Disassembly of charged protein structures has been previously observed with increasing ionic strength and arises from screened electrostatic interactions.^26–29,38,43,59,60^ At high salt concentrations, electrostatic interactions are more effectively screened (i.e., shorter Debye screening length), and proteins are expected to exist in monomeric, non-interacting states. For more highly negatively charged protein variants, larger ionic strengths are required to induce disassembly. For example, the assembly between Ceru+32o and GFP-11 had a FRET efficiency = 0.5 at 110 mM NaCl, whereas the assembly between Ceru+32o and GFP-32 had a FRET efficiency = 0.5 at 350 mM NaCl. The FRET efficiencies between Ceru+15r, Ceru+15a, and Ceru+32b and each of the negatively charged GFP variants at different NaCl concentrations and pH values are shown in **Figures S6-S8**, and the effect of pH on the assembled structures for Ceru+32o / GFP-17 is shown in **Figures S9 and S10**.

**Figure 4.**
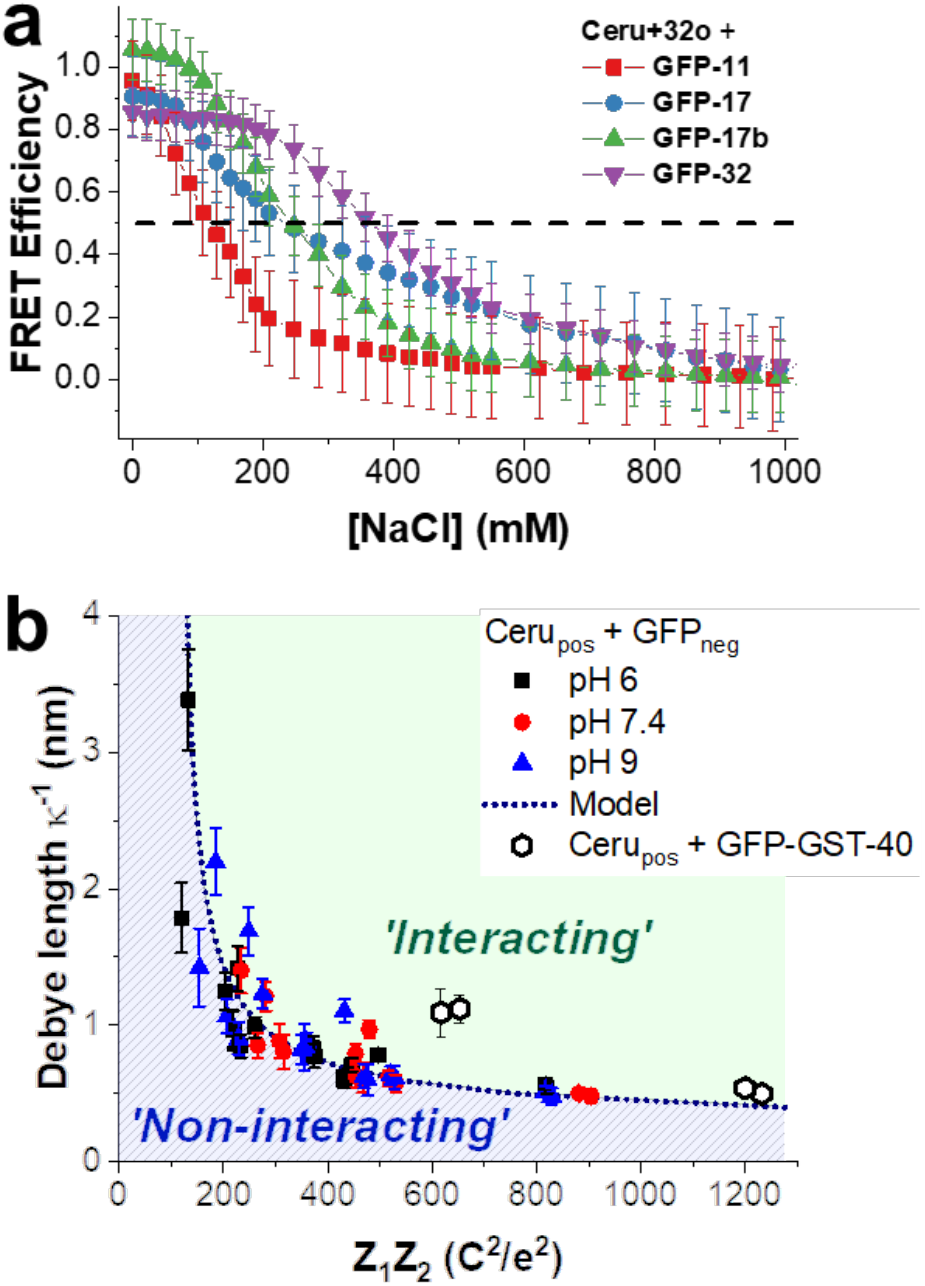
Assembly of supercharged proteins. (a) Measured FRET efficiency between Ceru+32o / GFP-11 (red square), GFP-17 (blue circle), GFP-17b (green triangle), and GFP-32 (purple triangle) at different salt concentrations. The dashed horizontal black line represents the critical FRET efficiency where proteins are either interacting (FRET > 0.5) or non-interacting (FRET < 0.5). (b) The critical Debye screening length *κ*^-1^ (calculated from the critical NaCl concentration using from Eq. 1) vs. Coulombic attraction (*Z*_*1*_*Z*_*2*_) for all supercharged GFP combinations at pH 6 (black squares), pH 7.4 (red circles), and pH 9 (blue triangles). The dashed line represents the fit to these data using Eq. 2. Critical Debye screening lengths for interactions between positive Ceru variants and GFP-GST-40 (open hexagons) are not included in this fit. The model represents a phase diagram: for Debye screening lengths *κ*^-1^ greater than or less than that predicted, oppositely supercharged proteins are expected to be macroscopically ‘interacting’ (green shaded area) or ‘non-interacting’ (blue shaded area), respectively.

Our results generally show that more highly charged proteins are robust to assembly into supramolecular structures at higher ionic strengths, which is attributed to the larger Coulombic attraction (*Z*_*1*_*Z*_*2*_) between oppositely charged proteins. To understand the role of net protein charge on assembly, we determined the critical salt concentration at which FRET efficiency = 0.5 (**Table S5**). The critical salt concentration was used to determine a corresponding Debye screening length *κ*^*-1*^:

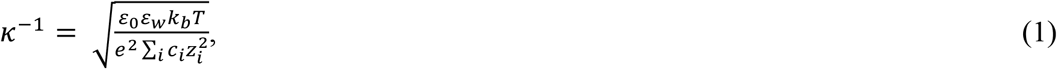

where *ε*_*0*_ is vacuum permittivity, *ε*_*w*_ is the dielectric constant of water, *k*_*b*_ is Boltzmann’s constant, *T* is absolute temperature, *e* is the elementary charge, and *c*_*i*_ and *z*_*i*_ are the number densities and valencies of the electrolyte ions (here NaCl). As shown in **Figure 4b**, the critical Debye screening length 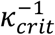 is inversely related to the product of net charge on each protein (*Z*_*1*_*Z*_*2*_). Upon increasing Coulombic attraction between oppositely charged proteins, a higher salt concentration (and smaller Debye screening length) is required to disrupt assembly. A similar trend is also observed at pH 6 and 9 for slightly different net charges (**Figure 1b**). Similar values of critical ionic strength were determined using different monovalent and divalent salts (KCl, KNO_3_, CaCl_2_ and MgSO_4_) (**Figure S11**), suggesting the observed trend is generalizable to different solution compositions.

We further used a simple electrostatic model to understand the observed inverse relationship between Coulombic attraction *Z*_*1*_*Z*_*2*_ and critical Debye screening length 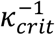. The electrical double-layer force *F*_*el*_ between two small spherical charged ions with diameter *σ* separated by distance *r* in solution is given by:^61^

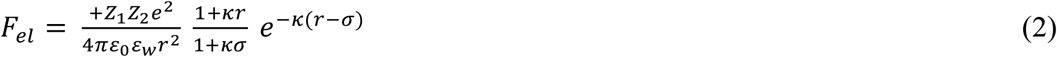

At a FRET efficiency = 0.5, the average positively charged Ceru variants are separated from negatively charged GFP variants by the Förster radius *R*_*0*_ = 5.5 nm. Here, we assume that a constant electrical double-layer force between oppositely charged proteins is required to maintain this separation distance. This model yields the dashed line in **Figure 4b** and shows an inverse relationship between Coulombic attraction *Z*_*1*_*Z*_*2*_ and critical Debye screening length 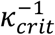. The data were best fit with an ion diameter *σ* = 4.4 ± 0.1 nm and an electric double-layer force *F*_*el*_ = 10.8 ± 0.8 pN (Eq. 2). Interestingly, the average center-of-mass separation between GFP-17 and Ceru+32o in the 16-mer was reported as 3.5-4.1 nm from the high-resolution cryo-EM structure,^38^ which is consistent with the electrostatic model. Moreover, the average electrical double-layer force between oppositely charged proteins with ≈1 nm separation is on the same order of magnitude as *F*_*el*_ measured between charged colloidal particles.^62^ We note that this simple model assumes an isotropic charge distribution around a spherical ion, though we expect that anisotropic charge distributions contribute to differences in 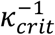 observed in the experimental data.

To assess the generality of the electrical double-layer relationship, FRET efficiencies were determined for mixtures of the positively charged Ceru variants and a fusion protein consisting of negatively supercharged glutathione S-transferase (GST-40) fused with neutrally charged GFP (named GFP-GST-40, **Figure S12**). GST-40 is a supercharged dimeric protein with a nominal net charge of -39 at pH 7.4.^36^ The critical Debye screening lengths for GFP-GST-40 and each positively charged Ceru variant are shown in **Figure 4b**, and the critical NaCl concentrations are provided in **Table S5**. Although an inverse relationship between Debye length and Coulombic attraction is observed, the critical Debye lengths for the fusion proteins are slightly larger than those predicted by the simple electrostatic model, which suggests that electrostatic attractions are readily screened, possibly arising from electrostatic shielding due to the neutrally charged GFP.

We further used the electrical double-layer analysis to develop an approximate phase diagram for supercharged assembly (**Figure 4b**). If the Debye screening length of the solution is larger than that predicted by 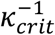, then oppositely supercharged proteins are expected to exist in an interacting state (defined when the average separation distance is less than the Förster distance *R*_*o*_ for a characteristic protein pair), and assembly is expected. Conversely, if the Debye length is smaller than 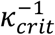, then supercharged proteins are expected to exist in non-interacting states. Overall, net charge provides a simple metric to predict whether supercharged GFP variants will interact and assemble in solution, but the precise phase diagram will change depending on protein concentrations or ratio of positively-to-negatively charged protein.^26^

### Assembled structures depend on precise inter-protein interactions

We further explored how changing surface charge distribution (at a constant net charge) affects the assembly of proteins. **Figure 5a** shows the FRET efficiency between negatively charged GFP variants (GFP-17 or GFP-17b) and positively charged Ceru variants (Ceru+32o or Ceru+32b) at different salt concentrations at pH 7.4. Because the net charge and Coulombic attraction are similar for these protein pairs, the FRET efficiency traces are similar, and nearly identical critical salt concentrations are determined. However, bulk average FRET measurements alone cannot provide additional detailed information on the assembly process for supercharged proteins.

**Figure 5.**
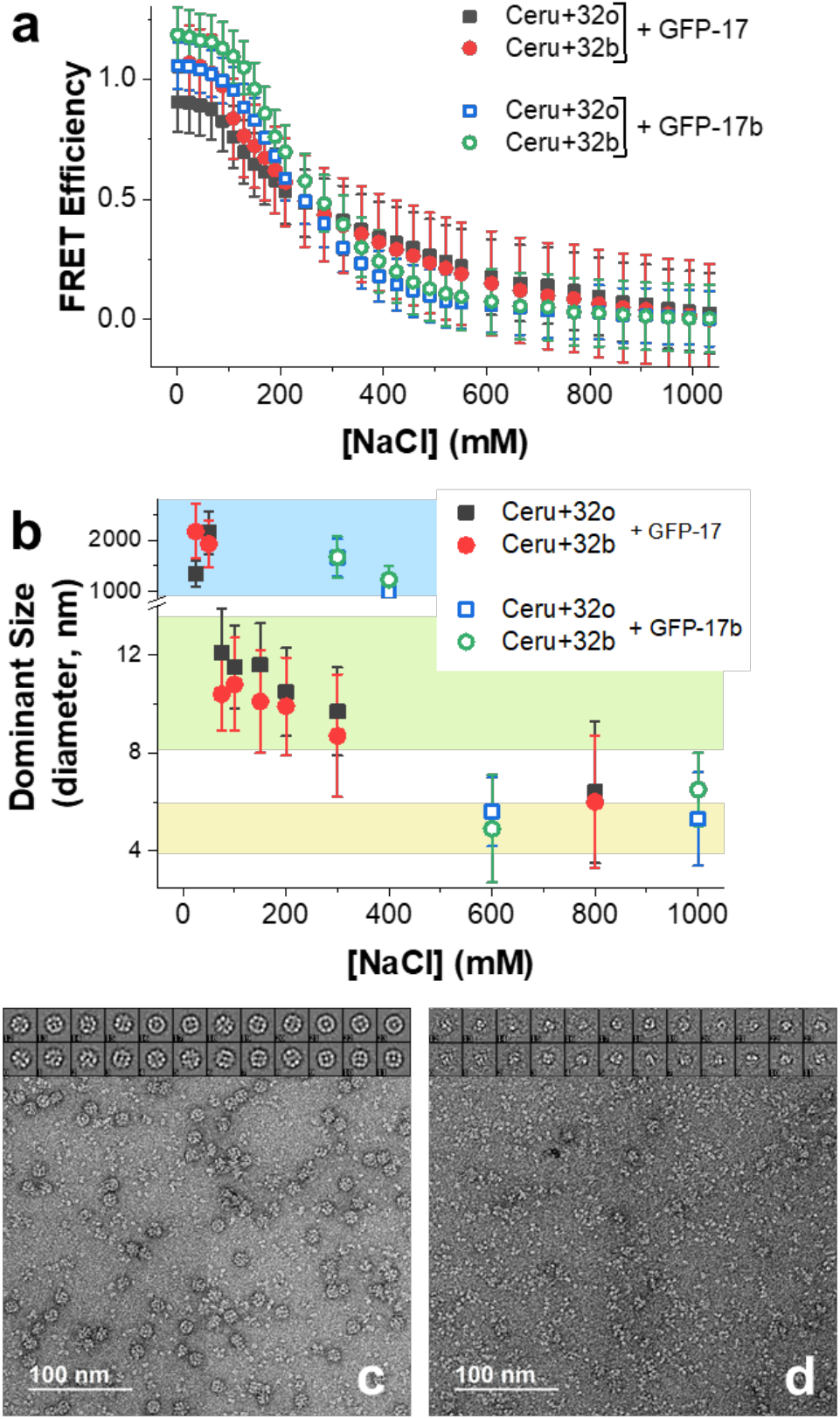
Effect of protein surface charge distribution on assembly. (a) FRET efficiencies between Ceru+32o / GFP-17 (black filled square), Ceru+32b / GFP-17 (red filled circle), Ceru+32o / GFP-17b (blue open square), Ceru+32b / GFP-17b (green open circle) at different concentrations of NaCl. (b) Dominant assembly size as measured by DLS for each protein combination at different concentrations of NaCl. (c) Representative TEM image of protomers from Ceru+32o / GFP-17 at 75 mM NaCl. The 24 class averages (*n* ≈ 2,000 particles) are shown at the top and show a highly homogenous structure. (d) Representative TEM image of protomers from Ceru+32b / GFP-17 at 75 mM NaCl. The 24 class averages (*n* ≈ 800 particles) are shown at the top and show a heterogeneous structure.

To understand the role of surface charge distributions, we studied supramolecular assembly of charged protein variants with different surface charge distributions using DLS, confocal microscopy, and transmission electron microscopy (TEM). **Figure 5b** shows the dominant assembly sizes determined from DLS. At low salt concentration, all of the oppositely charged protein pair combinations assemble into large (>1000 nm) aggregates with fractal-like structures (**Figure S13**).^38^ Upon increasing salt concentration, the average diameter of the dominant assembled structures between GFP-17 and either Ceru+32o or Ceru+32b was 10-12 nm, which is consistent with the aggregate size previously observed between Ceru+32o and GFP-17, suggesting a stable protomeric structure.^38^ On the other hand, the dominant assembled structures between GFP-17b and either Ceru+32o or Ceru+32b consisted of large aggregates (>1000 nm) for NaCl concentrations < 400 mM. Interestingly, no intermediate-sized protomeric structures were observed for protein pairs involving GFP-17b, which only differs from GFP-17 in the distribution of surface charges. At high NaCl concentrations (>600 mM), the dominant structure in solution from DLS matched the monomeric protein size for all protein combinations, indicating electrostatic-driven assembly was effectively screened.

Negative stain TEM was used to image protein combinations showing evidence of protomer structure formation. Ceru+32o and Ceru+32b formed a 10-12 nm assembled structures with GFP-17 at 50 mM NaCl. As expected, Ceru+32o and GFP-17 formed distinct, homogenous supramolecular ring-like assemblies, and class averages of selected particles (*n* ≈ 2,000) were consistent with prior reports (**Figure 5c**).^38^ However, assembled structures between Ceru+32b and GFP-17 were heterogeneous, and class averages of selected particles (*n* ≈ 800) did not yield evidence of an ordered structure (**Figure 5d**). Overall, these results highlight the role of surface charge distribution on facilitating and stabilizing specific intermolecular interactions promoting ordered assembly.

### Stability of assembled structures depends on size and specific inter-protein interactions

We next assessed the stability of supramolecular complexes by performing a series of dynamic subunit exchange measurements. First, the concentration dependence of supramolecular assemblies was assessed using dilution experiments, wherein the assembled protein structures were diluted with buffered solution, and equilibrium FRET ratios were measured (**Figure S14**). If proteins dynamically exchange with nearby protein partners in solution (which has been reported for electrostatically-driven complex coacervate formation),^63,64^ then the dominant assembled structure is expected to depend on solution composition and protein concentration. For assemblies between proteins with relatively small net charges (e.g., those involving Ceru+15 variants), our results show that the FRET ratio decreased with decreasing protein concentration, suggesting rapid dynamic subunit exchange in solution. However, for supramolecular assemblies consisting of proteins with larger net charges, the FRET ratio did not significantly change upon dilution, suggesting slow dynamic exchange in solution.

To understand the kinetic stability of assembled protein structures that are relatively insensitive to dilution, we performed a series of dynamic subunit exchange measurements using non-fluorescent analogs of the supercharged fluorescent proteins. Non-fluorescent analogs of the negatively charged GFP variants (containing a Gly-Gly-Gly construct in place of the chromophore)^65^ were designed and expressed. As expected, the negatively charged GFP variants and their corresponding non-fluorescent analogs (designated as ‘nf’) were observed to assemble with positively charged Ceru variants with identical average sizes and shapes as determined by DLS and TEM imaging (**Figure S15**). Dynamic exchange experiments were then performed by adding non-fluorescent proteins to a solution containing pre-assembled structures from oppositely supercharged fluorescent proteins, and changes in FRET were measured as the non-fluorescent analogs dynamically exchanged with GFP in the supramolecular assembled structures (**Figure 6a**). The time-dependent FRET ratio for GFP-17nf exchange with Ceru+32o / GFP-17 is shown in **Figure 6b** for a series of different NaCl concentrations. In all cases, the transient FRET ratio was found to decrease as a function of time when GFP-17nf is added to solution (at *t* = 0 min). The FRET ratio was not observed to change over the timescale of the measurement when buffer was added instead of GFP-17nf (**Figure S16**), suggesting that exchange of non-fluorescent GFP-17nf into the assembled structure (rather than photobleaching) is responsible for observed changes in fluorescence. In addition, the final FRET ratio at each salt concentration was measured by pre-mixing GFP-17 and GFP-17nf before triggering assembly with Ceru+32o (solid lines, **Figure 6b**). The transient FRET ratio approaches the final FRET ratio more rapidly at higher salt concentrations, indicating faster uptake of GFP-17nf and a more dynamic assembled structure. A series of dynamic exchange measurements was also performed with Ceru+32b / GFP-17, Ceru+32o / GFP-17b, and Ceru+32b / GFP-17b as a function of ionic strength (**Figure S17**).

**Figure 6.**
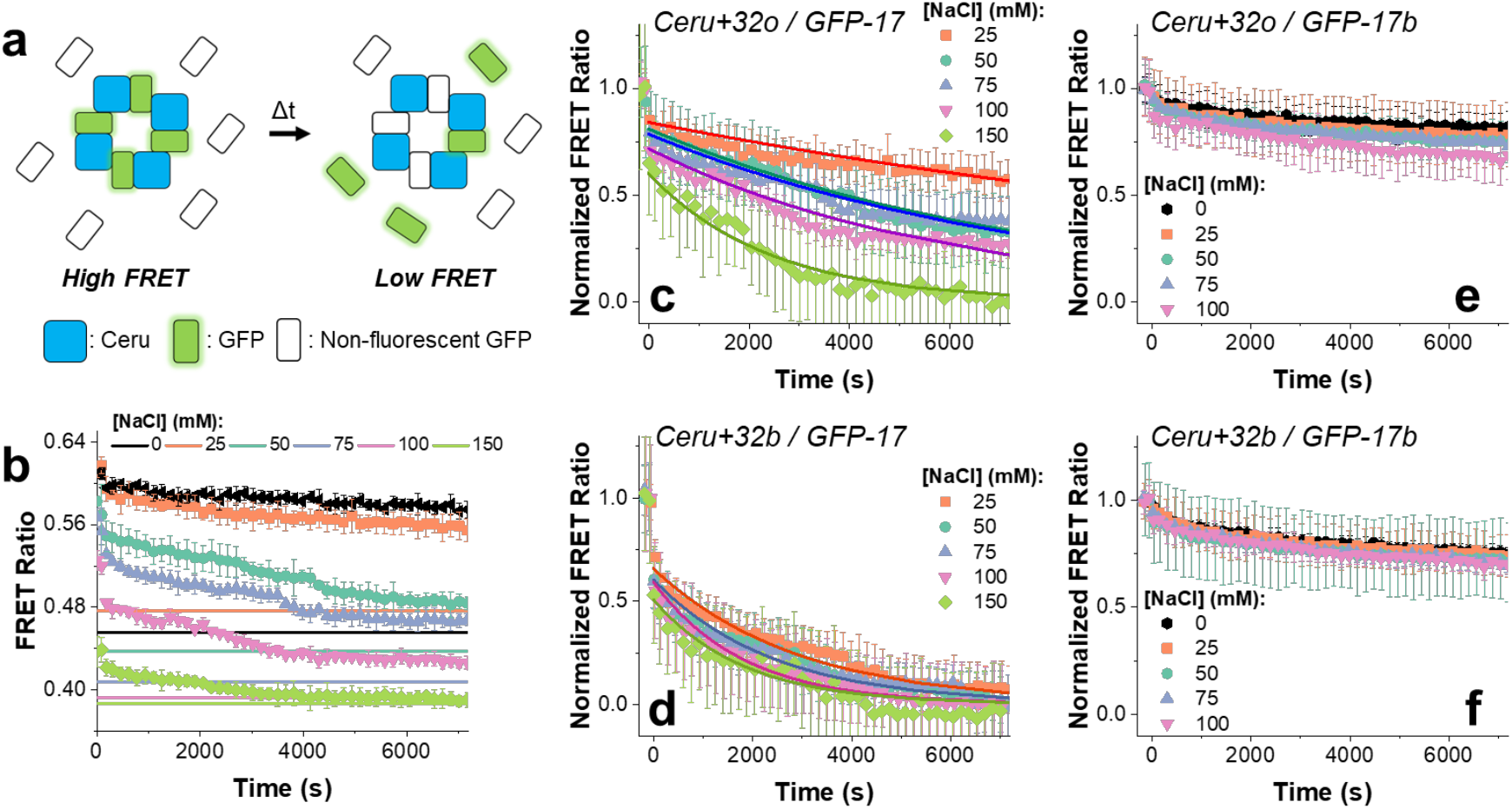
Dynamic exchange experiments. (a) Schematic of dynamic subunit exchange. As non-fluorescent GFP analog is incorporated into assembled structures, the FRET ratio decreases. (b) Transient FRET ratio between Ceru+32o / GFP-17 after the non-fluorescent negatively charged analog (GFP-17nf) is added to solution at different concentrations of NaCl. The solid lines represent the final FRET ratio with all GFP-17nf incorporated at each salt concentration. (c)-(f) Normalized transient FRET ratios using the initial (before non-fluorescent negatively charged analogs were added) and final FRET ratios at different salt concentrations for (c) Ceru+32o / GFP-17, (d) Ceru+32b / GFP-17, (e) Ceru+32o / GFP-17b, and (f) Ceru+32b / GFP-17b. Kinetic traces from modeled dynamic subunit exchange of protomers from Ceru+32o / GFP-17 and Ceru+32b / GFP-17 are shown as solid lines.

Transient normalized FRET ratios after addition of non-fluorescent negatively charged GFP analogs are shown in **Figures 6c-f**. The normalized FRET ratio for protomer-forming assemblies (i.e., assembly between GFP-17 and either Ceru+32o or Ceru+32b) decreases significantly after addition of GFP-17nf, and the rate of change varies with salt concentration (**Figures 6c and 6d**). In some cases, there was an initial rapid decrease in the transient FRET ratio, which was observed even for samples where large aggregates were removed by centrifugation (**Figure S18**). Based on these results, we hypothesize that the initial rapid decrease in FRET ratio arises from displacement of weakly associating proteins from the protomers, and the slower decrease in FRET ratio arises from dynamic subunit exchange within the protomer. On the other hand, large aggregate forming assemblies (i.e., assembly between GFP-17b and either Ceru+32o or Ceru+32b) show much slower change in normalized FRET ratio at all salt concentrations measured (**Figures 6e and 6f**). The difference in behavior between these two systems likely arises due to structural differences that affect the ability of proteins to exchange with nearby partners in the surrounding solution.

### Kinetic model for dynamic exchange

To quantitatively describe the kinetic stability experiments, we developed a simple kinetic model to understand the dynamic exchange of protein subunits for complexes between Ceru+32o / GFP-17 and Ceru+32b / GFP-17 (**Table S6**).^66–68^ In this model, monomers dissociate from the protomer sequentially according to a unimolecular rate constant *k*_*d*_, which is assumed to be constant for all protomers. The probability of a protomer dissociating to yield a free fluorescent monomer of GFP-17 (G) or free non-fluorescent analog (A) is determined by the composition of the assembled protein structure. In terms of association, monomeric GFP-17 (G) or non-fluorescent analog (A) proteins form protomers with a bimolecular rate constant *k*_*a*_. Because of the strong binding energy between oppositely charged proteins, association reactions are assumed to be fast, and dissociation reactions are rate limiting (**Figure S19**). Similar exchange kinetics were observed when different concentrations of non-fluorescent analogs are added to solution, which supports the assumption that dissociation is rate limiting (**Figure S20**). A detailed description of the kinetic model is provided in Supporting Information.

Results from the kinetic model are shown as solid lines in **Figures 6c-6d**. The incorporation of non-fluorescent analog (A) into the protomers is simulated by numerically solving the coupled differential rate equations while matching experimental conditions ([G_8_A_0_]_,initial_ = 0.05 μM, [G]_initial_ = 0 μM, [A]_initial_ = 4 μM).^69^ Here, *k*_*d*_ was varied to directly compare the simulated fraction of GFP-17 remaining in the protomer to the experimentally determined normalized FRET ratio. For protomer assembly between Ceru+32o and GFP-17 (**Figure 6c**), *k*_*d*_ increases as the salt concentration increases (**Figure S21**), ranging from 0.0004 s^-1^ to 0.003 s^-1^ at 25 mM and 150 mM NaCl, respectively. These values of *k*_*d*_ suggest kinetic lifetimes ranging from 7 to 40 minutes depending on the solution composition. For protomer assembly between Ceru+32b and GFP-17 (**Figure 6d**), exchange rates are less sensitive to salt concentration; *k*_*d*_ ranges from 0.0025 to 0.004 s^-1^ at 25 mM and 150 mM NaCl, respectively. Overall, the protomers formed from Ceru+32b / GFP-17 are observed to be less stable compared to protomers formed from Ceru+32o / GFP-17, which could arise from the heterogeneous structures of Ceru+32b / GFP-17 assemblies (**Figure 5d**) and lack of cooperative interactions between subunits. In addition, faster exchange kinetics could arise from protomers with fewer total subunits (**Figure S22**).

In general, the kinetic lifetimes for dynamic subunit exchange measurements of protomers are consistent with those observed in endogenous biological protein complexes,^70^ which range from seconds (e.g., RNA polymerase)^71^ to days (e.g., nuclear pore complex).^72^ From this view, assembly between oppositely supercharged proteins provides a mechanism to drive long-lasting association between two otherwise non-interacting proteins. However, the stability of the assembled structure strongly depends on the precise intermolecular interactions and solution conditions. In addition, assembly between oppositely supercharged proteins can be used to generate extremely stable biomaterials with transport-limited exchange leading to kinetic lifetimes >1 day (i.e., large aggregate forming assemblies).

## Conclusions

In this work, we study the assembly process of oppositely supercharged proteins using a combination of experiments and computational modeling. Our results highlight the role of surface charge distributions on the supramolecular assembly process for oppositely supercharged proteins. Using minimally mutated variants of supercharged proteins (GFP-min and Ceru-min) that retain only the mutations involved in stabilizing the assemblies, we systematically explored the role non-interacting charges play on assembly. Minimally charged variants were not observed to assemble into ordered hierarchical structures. Results from MD simulations show that the values of *ΔG*^*0*^_*bind*_ are relatively large for both the minimally charged and fully charged variants compared to naturally occurring protein-protein interfaces.^54^ However, *ΔG*^*0*^_*bind*_ for the minimally charged protein variants are significantly smaller (∼30%) than those of the fully charged variants even though the interfaces retain all specific H-bonding and salt bridge interactions. Such differences in binding energy are attributed to long-range electrostatic interactions between the fully charged variants. From this view, specific inter-protein interactions contribute to the bulk of the binding energy between oppositely supercharged proteins, but non-interacting excess charges are required to stabilize the assembled protein complexes.

Our results further highlight that identification of specific inter-protein interactions from high resolution structures alone is insufficient to fully describe the interactions and assembly process between oppositely supercharged proteins. From this view, our results will be useful for informing the rational design of new hierarchical structures based on predictions of complementary surfaces.^73–77^ In particular, rational design methods for supercharged proteins should take into consideration long-range electrostatic interactions away from the interface for stabilizing protein complexes, which are known to play a role in kinetic assembly and disassembly processes involving supramolecular protein complexes.

Our results elucidate how net charge and average Coulombic attraction can predict whether oppositely supercharged proteins will interact as a function of ionic strength conditions. We found that the supramolecular structure strongly depends on the precise distribution of charges on the proteins surface. Prior work has shown that different assembled structures can be obtained from a common parent protein building block by changing the supramolecular assembly strategy (e.g., ferritin cage proteins).^20,22,23,30,42^ Importantly, our results show that proteins with similar net charges but different surface charge distributions give rise to different assembled structures. In addition, the kinetic stabilities of these different supramolecular structures were found to depend on surface charge distribution. Overall, our results show that supercharging proteins provides an efficient method to create stable, long-lived protein assemblies while further highlighting the role of surface charge distribution and precise stabilizing inter-protein interactions on the structure of assembled supercharged proteins.

## Supporting Information

Description of materials, experimental methods, and computational methods including amino acid sequences, denaturing protein gels, and UV-vis and fluorescence spectra; Depiction of simulated protein-protein interfaces; PMFs of orientational, positional, and conformation constraints for bound and unbound states; Plots of FRET efficiency versus NaCl for protein assemblies at pH 6, 7.4, and 9; Plot of soluble fraction of GFP-17 / Ceru+32o assemblies versus NaCl concentration; FRET efficiency versus salt concentration for protein assemblies for different salt identities; TEM images of GFP-17 / Ceru+32o at different pH values and at different ratios; Confocal microscopy images of protein assembly; Plot of FRET ratio versus protein concentration; Comparison of assemblies from GFP-17 and non-fluorescent analog; Plots of transient FRET ratios after buffer addition, after non-fluorescent analog addition, and with/without large aggregates; Plot of final FRET ratios versus non-fluorescent analog concentrations; Simulated exchange kinetics with different association rate constants *k*_*a*_ and different protomer sizes; Plot of measured and simulated exchange kinetics for different concentrations of non-fluorescent analog; Plot of modeled dissociation rate constants *k*_*d*_; Tables containing contributing terms to *ΔG*^*0*^_*bind*_ for each interface; Table with critical salt concentrations and protein charges for different assemblies; Table with dynamic subunit exchange kinetic model; Table with relevant MD simulation parameters.

## Acknowledgments

This work was supported by the U.S. Department of Energy, Office of Science, Basic Energy Sciences under Award #DE-SC0022035. M.I.J. acknowledges a Beckman Institute Postdoctoral Fellowship and a Mistletoe Fellowship. D.S. and P.D.B. acknowledge support from NSF MCB-1845606. We thank Prof. Andrew Ellington for providing plasmids encoding for GFP-11, GFP-17, GFP-32, and Ceru+32o, and we thank Prof. Catherine Murphy for use of the Malvern Zetasizer Nano ZS. P.D.B. thanks Balaji Selvam, Soumajit Dutta, and Jiming Chen for useful discussions and The Blue Waters Petascale Computing Facility and National Center for Supercomputing Applications (NCSA), which are supported by the National Science Foundation (awards OCI-0725070 and ACI-1238993) and the state of Illinois.

---

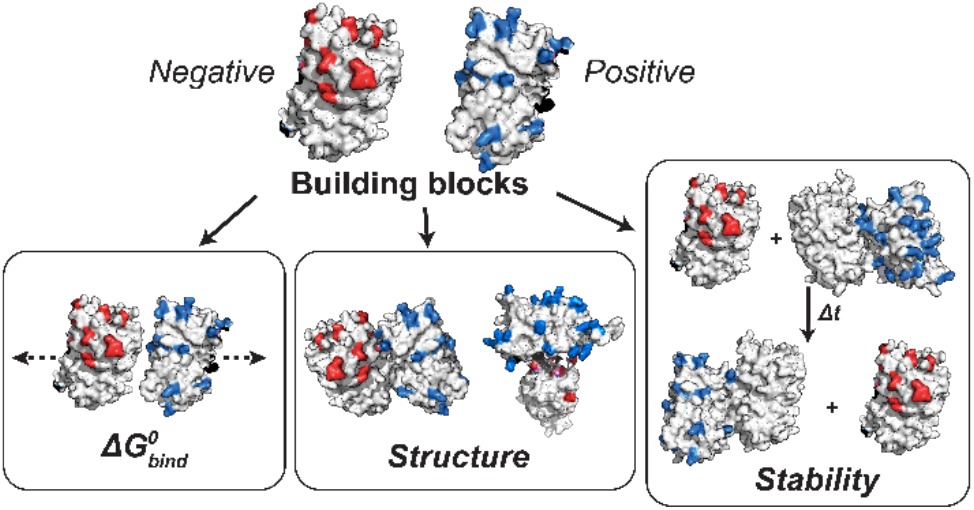
TOC Graphic.

**Synopsis:** The supramolecular assembly process is studied for a series of oppositely charged proteins using experiments and simulations to understand the fundamental mechanisms governing assembly of supercharged proteins.

