## Supporting Information for "Understanding Supramolecular Assembly of Supercharged Proteins"

**Table of Contents:**

| | |
|---|---|
| Page S-3 | Protein cloning |
| | Protein expression and purification |
| Page S-4 | Preparation of single-protein and mixed protein solutions |
| | Dynamic light scattering (DLS) measurements |
| | Fluorescence measurements |
| Page S-5 | Negative stain transmission electron microscopy (TEM) |
| | Subunit exchange measurements |
| Page S-6 | Subunit exchange kinetic model |
| | Outline of computational methods |
| Page S-7 | Replica exchange umbrella sampling simulations for separation PMF |
| Page S-8 | Overall methodology of computing the contribution of each term to $\Delta G^0_{bind}$ |
| Page S-12 | Adaptive Biasing Force (ABF) based simulations for calculating $W(\xi)$ |
| | Calculating separation PMF using REUS |
| Page S-13 | Error Analysis |
| Page S-14 | Figure S1. Four protein-protein interfaces in 16-mer. |
| Page S-15 | Figure S2. Intra-planar orientational and positional constraints in the bound state. |
| Page S-16 | Figure S3. Inter-planar orientational and positional constraints in the bound state. |
| Page S-17 | Figure S4. Intra-planar conformational constraints in the bound state. |
| Page S-18 | Figure S5. Inter-planar conformational constraints in the bound state. |
| Page S-19 | Figure S6. FRET efficiency vs. NaCl for supercharged GFP assemblies at pH 7.4. |
| | Figure S7. FRET efficiency vs. NaCl for supercharged GFP assemblies at pH 6. |
| Page S-20 | Figure S8. FRET efficiency vs. NaCl for supercharged GFP assemblies at pH 9. |
| | Figure S9. Soluble fraction of GFP-17 / Ceru+32o assemblies versus NaCl and pH. |
| Page S-21 | Figure S10. TEM images of GFP-17 / Ceru+32o at different pH values |
| Page S-22 | Figure S11. Critical salt concentrations for supercharged assembly in different types of salt. |
| Page S-23 | Figure S12. FRET efficiency vs. NaCl for GFP-GST-40 / Ceru variant assemblies. |
| | Figure S13. Confocal microscopy images of assembly at no salt concentration |

Page S-24 Figure S14. FRET ratio at different equimolar concentrations of oppositely supercharged proteins.
Figure S15. Comparison of assembly between Ceru+32o / GFP-17 and GFP-17nf
Page S-25 Figure S16. Transient FRET ratio after buffer addition.
Page S-26 Figure S17. Transient FRET ratio after addition of non-fluorescent analog.
Page S-27 Figure S18. Transient FRET ratio with and without large aggregates.
Figure S19. Simulated fraction of GFP remaining in protomer at different $k_a$.
Page S-28 Figure S20. Exchange kinetics versus ratio of non-fluorescent to fluorescent GFP.
Page S-29 Figure S21. Modeled dissociation rate constants $k_d$.
Figure S22. Simulated fraction of remaining GFP for differently sized protomers.
Page S-30 Figure S23. Amino acid sequences of positively charged Ceru variants.
Page S-31 Figure S24. Amino acid sequences of negatively charged GFP variants.
Page S-32 Figure S25. Amino acid sequences of minimally mutated variants.
Figure S26. Amino acid sequence of GFP-GST-40.
Page S-33 Figure S27. Denaturing gel of supercharged proteins.
Page S-34 Figure S28. UV-vis absorption and fluorescence spectra of GFP variants.
Page S-35 Figure S29. TEM images at different ratios of GFP-17 and Ceru+32o.
Figure S30. Final FRET ratio versus non-fluorescent analog concentration.
Page S-36 Figure S31. Constraints in dimer separation simulations.
Page S-37 Figure S32. Intra-planar conformational constraints in the unbound state.
Page S-38 Figure S33. Inter-planar conformational constraints in the unbound state.
Page S-39 Table S1. Contributing terms to $\Delta G^0_{bind}$ for 'Ceru+32 Clockwise' interface
Page S-40 Table S2. Contributing terms to $\Delta G^0_{bind}$ for 'GFP-17 Clockwise' interface
Page S-41 Table S3. Contributing terms to $\Delta G^0_{bind}$ for 'Inter-GFP-17' interface
Page S-42 Table S4. Contributing terms to $\Delta G^0_{bind}$ for 'Inter-Ring' interface
Page S-43 Table S5. Critical NaCl concentrations and charges for different assemblies.
Page S-44 Table S6. Dynamic subunit exchange kinetic model for 16-mer.
Page S-45 Table S7. List of Modelled Residues.
Table S8. Mutations to construct minimal mutants.
Page S-46 Table S9. General System information regarding each Simulation System.
Page S-47 References

### Experimental Methods

**Protein cloning:** Figure 1a shows the set of supercharged superfolder GFP variants that were used in this work, including previously reported and newly designed variants. GFP variants including GFP-11, GFP-17, and GFP-32 and positively charged Ceru+32o were based on previously reported supercharged GFP variants.[1] We created several new supercharged variants (as noted below) by introducing amino acid mutations into wild-type superfolder GFP. The amino acid sequences for all supercharged variants used in this work and their unmodified counterparts are provided in the Supporting Information (Figures S23-S26). The ROSIE online server[2] was used to generate new supercharged GFP variants using two routines: a deterministic supercharging routine that selects mutations based on fewest average neighboring atoms per side chain atom (AvNAPSA)[3] and a supercharging routine that uses Rosetta-based energy calculations to choose surface mutations.[4] These two routines generate nearly complementary sets of amino acids for a given target net charge. The net charge on each variant was determined by the computational tool PROPKA, which predicts the pKa values of ionizable groups in proteins on the basis of the folded structure (Figure 1b).[5,6]

To enable Förster resonance energy transfer (FRET) measurements between oppositely supercharged proteins, a Y66W mutation to blue shift the GFP chromophore and associated stabilizing mutations (F146G, N147I, H149D) were introduced to all positively charged variants (which were subsequently called 'Ceru').[1,7–9] Minimally mutated GFP (name GFP-min) and Ceru (named Ceru-min) variants were generated by only keeping the mutated charged residues from the 16-subunit protomer-forming variants that were previously identified as actively participating in the inter-protein interfaces.[1] Non-fluorescent supercharged GFP variants (named GFP-11nf, GFP-17nf, GFP-17bnf, and GFP-32nf) were generated with S65G and Y66G mutations to generate a Gly-Gly-Gly construct in place of the chromophore.[10] A fusion protein between previously reported negatively supercharged glutathione S-transferase (GST-40)[3] and neutrally-charged GFP was also generated (named GFP-GST-40). All genes were purchased as gBlocks (IDT) and inserted into pET21b expression vectors. DNA plasmids were transformed into NEB 5-alpha *E. coli* competent cells and their sequences verified by single pass Sanger DNA sequencing (ACGT, Inc.) before protein expression.

**Protein expression and purification:** GFP variants were expressed in *E. coli* T7 Express cells (NEB). Prior to induction, cells were grown (37 °C, shaking at 225 rpm) to an $OD_{600}$ between 0.6 and 0.8 in lysogeny broth (LB) for GFP variants and Terrific Broth (TB) for Ceru variants. Expression of protein variants was induced with 100 μM of β-D-1-thiogalactopyranoside (IPTG), and cells were grown at 25 °C for 16-20 hours. Cells were harvested by centrifugation (7000 × g for 10 min) and resuspended in lysis buffer containing 50 mM phosphate buffer (pH 7.4), 10 mM imidazole, and 2 M sodium chloride. All purification steps were performed either on ice or at 4 °C. The cell medium was sonicated with a 700 W sonicator (10 s on, 80 s off, for 30 cycles at 60% amplitude capacity). Following sonication, the soluble and insoluble fractions were separated by centrifugation (15,000 × g for 25 min). Supercharged proteins were purified from other soluble components using Ni-NTA agarose chromatography under native conditions. The wash buffer contained 50 mM phosphate buffer, 2 M NaCl, and 35 mM imidazole (pH 7.4), and the elution buffer contained 50 mM phosphate buffer (pH 7.4), 2 M NaCl, and 200 mM imidazole (pH 7.4). Purified protein samples were concentrated to ~10 mg/mL with centrifugal filters (molecular weight cutoff of 10 kDa), dialyzed in phosphate buffered saline (PBS, 137 mM NaCl, 2.7 mM

KCl, pH 7.4), and stored at 4 °C. Denaturing gels and absorption/emission spectra indicated each variant expressed monomerically and with the absorbance and fluorescence emission spectral properties expected for GFP parent proteins (Supporting Information, Figures S27 and S28). Dynamic light scattering (DLS) was used to check for aggregation, denoted as the presence of particles larger than the ~5-7 nm monomeric proteins, as discussed below. All proteins were stable for several months at the storage conditions described above.

**Preparation of single-protein and mixed protein solutions:** Proteins were slowly pipetted into the desired buffer and then mixed slowly by gentle swirling of the tube. For solutions containing oppositely supercharged proteins, both variants were diluted from stock solutions into buffer, and the negatively charged protein was mixed into the positively charged variant. Measurements were made in three different filter-sterilized buffer solutions: 50 mM Bis Tris buffer (pH 6.0), 50 mM Tris buffer (pH 7.4), and 50 mM Tris buffer (pH 9.0). Single component particle size was measured in each buffer to confirm consistency and lack of protein aggregation. DLS and fluorescence measurements were conducted on samples at room temperature (25 °C) approximately 15-30 min after mixing oppositely charged proteins. Repeated measurements were performed approximately 30-60 min after the initial measurements to ensure there was no drift in measurement and to monitor the progression of the assembly reactions.

**Dynamic light scattering (DLS) measurements:** DLS experiments were conducted using a Malvern Zetasizer Nano ZS. Measurements were made with default 175° backscatter optical configuration, automatic measurement duration, and in general-purpose analysis mode. Unless otherwise noted, experiments were performed at a monomer concentration of 0.1 mg/mL. For all conditions referenced (except where noted), at least three replicate measurements were obtained, and data from the averaged histograms are reported. The average particle sizes reported correspond to the average size of the dominant peak in the volume-averaged histogram, and the reported errors correspond to the standard deviation of this peak.

**Fluorescence measurements:** Fluorescence measurements were performed using a Cary Varian fluorometer. Positively charged Ceru variants serve as donors in FRET measurements and have excitation and emission maxima at $\lambda_{ex}$ = 433 nm and $\lambda_{em}$ = 475 nm, respectively. Negatively charged GFP variants serve as FRET acceptors and have excitation and emission maxima at $\lambda_{ex}$ = 485 and $\lambda_{em}$ = 510 nm, respectively. Equimolar concentrations of Ceru and GFP variants were mixed at a concentration of 0.1 mg/mL (unless otherwise noted). Aliquots of a stock solution of 3 M NaCl (prepared in the same buffer solution) were sequentially added to the mixture to change ionic strength, and fluorescence intensity was measured after the solution was mixed and allowed to equilibrate (typically after ~10 s).

FRET efficiency between different oppositely supercharged protein pairs was measured at different concentrations of NaCl using three-cube quantification, which requires three different measurements, as described below.[11] First, at the excitation wavelength that directly excites the donor molecule, the emission intensity is measured at the acceptor's and donor's emission maximum, $F^{ex_D,em_A}$ ($\lambda_{ex}$ = 433 nm, $\lambda_{em}$ = 510 nm) and $F^{ex_D,em_D}$ ($\lambda_{ex}$ = 433 nm, $\lambda_{em}$ = 475 nm), respectively. Next, the fluorescence intensity at the acceptor's emission maximum is measured at an excitation wavelength that selectively excites the acceptor, $F^{ex_A,em_A}$ ($\lambda_{ex}$ = 485 nm, $\lambda_{em}$ = 510 nm). The change in acceptor fluorescence intensity (*nF*) from FRET corrected for direct acceptor excitation and donor bleed through is:

$$nF = F^{ex_D,em_A} - \alpha F^{ex_A,em_A} - \beta F^{ex_D,em_D}, \quad \text{(S1)}$$

where $\alpha$ and $\beta$ are constants. In an acceptor-only measurement, the extent to which short (i.e., donor) wavelength excitation directly excites the acceptor is determined relative to the excitation at the longer (i.e., acceptor) wavelength, which results in the constant $\alpha = F_A^{ex_D,em_A} / F_A^{ex_A,em_A}$. Similarly, the amount of donor bleed-through into the acceptor emission maximum is determined by a donor-only measurement which provides the calibration constant $\beta = F_D^{ex_D,em_A} / F_D^{ex_D,em_D}$. The following equation was used to quantify apparent FRET efficiency ($Ef_A$) from the three emission measurements:

$$Ef_A = \gamma \frac{nF}{\alpha F_A^{ex_A,em_A}}. \quad \text{(S2)}$$

Here, $\gamma$ is the ratio of acceptor and donor extinction coefficients at donor excitation (i.e., $\gamma = \varepsilon_A^{ex_D} / \varepsilon_D^{ex_D}$), which were measured by UV-vis absorption spectroscopy. Unless otherwise noted, FRET efficiencies were measured for each supercharged protein combination at each NaCl concentration in triplicate.

**Negative stain transmission electron microscopy (TEM):** Mixed protein solutions were diluted to a concentration of 100 nM in a buffered solution with identical composition (ionic strength and pH) to that of the undiluted solution immediately prior to sample preparation. A 5-µL aliquot of sample was deposited onto a glow-discharged 400 mesh continuous carbon grid (Ted Pella, Inc.), and excess sample was removed after 3 min by wicking with a piece of filter paper. The bound particles were stained by washing the grid in 4 different 30-µL drops of 2% (w/v) uranyl acetate. The grid was floated on a fifth uranyl acetate droplet for 30 s before excess stain was removed by blotting with filter paper, and the grid was allowed to air-dry.

Images of stained protein assemblies were collected using a JEOL-2100 transmission electron microscope operated at 200 keV at a nominal magnification of 60,000× (pixel size of 1.9 Å) with a manually set defocus range of approximately -1 to 1.5 µm. All data were acquired under low dose conditions (≈35 $e^-/Å^2$). Approximately 50-70 images for each protein combination were manually recorded using a Gatan UltraScan 2k × 2k camera. TEM measurements were carried out in the Materials Research Laboratory Central Research Facilities, University of Illinois.

The initial image processing and classification steps were performed using the EMAN2 workflow.[12] Particles were first selected from micrographs using EMAN2 e2boxer and extracted in 128 pixel × 128 pixel boxes. CTF correction was performed using EMAN2 e2ctf, and particles were then subjected to 2D reference-free alignment and classification using EMAN2 e2refine2d. The initial 48 2D class averages were manually inspected to check for classes that appeared to correspond to particulate contaminants. Particles comprising such classes (typically less than 5% of total particles) were removed, and a second round of 2D reference-free alignment was performed to generate 24 2D class averages. No symmetry operator was applied at any point in this analysis.

**Subunit exchange measurements:** Dynamic subunit exchange measurements were performed by measuring changing FRET intensity between positively charged Ceru variants and negatively charged GFP variants after adding a non-fluorescent analog of the negatively charged GFP. Oppositely supercharged fluorescent proteins were mixed at equimolar concentrations (0.013 mg/mL) in 400 µL of buffer (50 mM Tris, pH 7.4) with variable concentrations of NaCl and allowed to incubate for 15-30 minutes. FRET intensity of the initial assembly was determined using a Cary Varian fluorometer by measuring the FRET ratio, which is defined as GFP emission intensity at Ceru excitation ($F^{ex_D,em_A}$) normalized by GFP emission intensity at GFP excitation ($F^{ex_A,em_A}$):[1]

$$FRET\ Ratio = \frac{F^{ex_D,em_A}}{F^{ex_A,em_A}}. \quad \text{(S3)}$$

The same excitation and emission wavelengths described above were used for FRET ratio measurements. After measuring the initial FRET ratio, a 100-μL aliquot containing the non-fluorescent analog of the GFP was added to solution and rapidly mixed by pipetting. The final concentration of non-fluorescent protein was 0.1 mg/mL (10-fold excess), unless otherwise noted. FRET intensity was measured every 12 seconds for at least 2 hours after adding the non-fluorescent protein. Dynamic exchange measurements were performed in buffered solution with different salt concentrations including 0, 25, 50, 75, 100, and 150 mM NaCl. All kinetic measurements were performed in triplicate at room temperature.

The final FRET ratios (defined as the FRET ratio values after all non-fluorescent protein had been added) were determined by pre-mixing the negatively charged GFP variant and non-fluorescent analog prior to adding the positively charged Ceru variant. The final FRET ratio was determined before and after a kinetic measurement to ensure that the value did not change over the timescale of measurement. All FRET ratio measurements were performed in triplicate.

**Subunit exchange kinetic model:** A simple kinetic model based on previous models of protein assembly was developed to help explain the dynamic exchange measurements.[72–74] The kinetic model requires some assumptions and initial conditions. First, all fluorescent proteins are assumed to be assembled into protomers containing 8 positively and 8 negatively charged proteins, consistent with prior reports for Ceru+32o / GFP-17.[38] We also assume that protomers from Ceru+32b / GFP-17 have the same number of subunits, consistent with DLS measurements (Figure 5b) and that excess non-fluorescent analogs of negatively charged GFP-17 do not affect the initial assembled protomers. Assembly between Ceru+32o / GFP-17 was observed by TEM to be invariant to GFP-17 concentration, which supports the assumption (Figure S29). Upon addition of non-fluorescent analog, fluorescent subunits dynamically exchange with the non-fluorescent analog, and FRET ratio decreases until the system approaches equilibrium (i.e. the final FRET ratio). Importantly, the final FRET ratio was found to change linearly with concentration of non-fluorescent analog (Figure S30), which suggests that the normalized FRET ratio is related to the fraction of non-fluorescent analog incorporated into protomers. In our model, monomers dissociate from the protomer sequentially according to some unimolecular rate constant $k_d$. While $k_d$ is constant for all protomers, the probability of a protomer dissociating into a free GFP-17 (G) or free non-fluorescent analog (A) is determined by its composition. Monomeric GFP-17 or non-fluorescent analog can re-form the protomer with some bimolecular rate constant $k_a$. The rate expressions for all possible dissociation and association processes are described in Supporting Information Table S6. The incorporation of non-fluorescent analog into the protomers is simulated at experimental conditions ($[G_8A_0]_{,initial}$ = 0.05 μM, $[G]_{initial}$ = 0 μM, $[A]_{initial}$ = 4 μM) using Kintecus.[75] To match experimental results, $k_d$ was varied to relate the simulated fraction of GFP-17 remaining in the protomer to the normalized FRET ratio.

**Outline of computational methods:** Atomistic molecular dynamics (MD) simulations were used to calculate the absolute binding free energy ($\Delta G^0_{bind}$) between oppositely supercharged and minimally mutated proteins. A detailed description of the computational methods is provided in the Supporting Information, and only a brief summary is provided here. The methodology used in this work was first established by Gumbart *et al.*, who computed the binding free energy of a protein-protein barnase-barstar dimeric complex.[13] Briefly, the $\Delta G^0_{bind}$ values were computed by implementing a separation potential of mean force (PMF)-based method where the proteins are pulled apart from a bound state (site) to an unbound state (bulk). To ensure smooth separation, the proteins were pulled along a vector $\vec{r}$ (defined along centers of masses of the two proteins) in the presence of configurational, positional, and conformational restraints (shown in Supporting Information, Figure S31). Replica Exchange Umbrella Sampling (REUS) was employed to sample the phase space along $\vec{r}$ on windows evenly spaced between the bound and unbound states. To compute the contribution of each of the positional and orientational restraints, Adaptive Biasing Force (ABF) simulations were used. To compute the contribution of the conformational restraints, the ABF simulations were used to generate the starting windows, where the windows were defined along that particular conformational restraint. Umbrella Sampling was then used to sample along the collective variable. Multistate-Bennett Acceptance Ratio (MBAR) was used to compute the PMF.[14]

**Replica exchange umbrella sampling simulations for separation PMF**

The high-resolution structure of the 16-subunit protomer between Ceru+32o / GFP-17 (PDB: 6MDR) was used as the starting structure for the simulations.[1] The four previously described interfaces within the protomer (Ceru+32 Clockwise, GFP-17 Clockwise, Inter-GFP-17, and Inter-ring) were prepared as separate systems by considering only the two proteins that preserved each interface. The missing loop regions in the proteins were modelled using MODELLER (Supporting Table S7).[15] The minimal mutant systems were made by mutating the residues not involved in the interfaces back to wild type superfolder GFP using open-source PyMOL,[16] resulting in four more systems (Ceru_min Clockwise, GFP_Min Clockwise, Inter-GFP_Min, Inter-ring_Min). The list of mutations performed to obtain the minimal mutants is listed in Supporting Table S8. The termini were capped using neutral termini caps Acetyl (ACE) and N-MethylAmide (NME) for the N- and C-termini respectively. The systems were solvated using TIP3P water[17] and 75 mM NaCl, to neutralize the system and mimic experimental conditions. TIP3P waters were added such that there was a 20 Å thick layer of water molecules on all sides of the proteins. Information specific to each system, such as box size, no. of atoms, and the various components are noted in Supporting Table S9. The solvation was done using the tleap module of Amber18[18,19] with the atomic interactions characterized using the Amber14 ff14SB forcefield.[20] The masses of hydrogen atoms (except of those that are part of the protein) were repartitioned to 3.024 Daltons, a technique that facilitates usage of a longer timestep (4 fs).[21] Parmed, a part of the AmberTools19 package, was used for this purpose.[22,23]

The eight systems were subjected to a two-stage minimization process where each stage was 20,000 steps. The first 15,000 steps in each stage used the gradient descent method, followed by 5,000 steps using the conjugate gradient method. In the second stage, the hydrogen bonds were constrained using the SHAKE algorithm.[24] The systems were then subjected to heating at NVT from 0 to 300 K, with a harmonic restraint of 10 kcal $mol^{-1}$ Å$^{-2}$ placed on the protein backbone to preserve the conformation. The systems were then equilibrated for 20 ns at NPT conditions, using a harmonic restraint of 10 kcal $mol^{-1}$ Å$^{-2}$ and periodic boundary conditions. The temperature was maintained at 300 K using Langevin dynamics with a 2 $ps^{-1}$ damping coefficient. Pressure was maintained at 1 bar using a Berendsen Barostat.[25] Electrostatic interactions were treated using the Particle Mesh Ewald method.[26] Nonbonded cutoff was set to 10 Å. All aforementioned simulations were performed using the AMBER simulation package[18,19,22,23,27–29] on NVIDIA Tesla K20X GPUs on the Blue Waters Supercomputer.

**Overall methodology of computing the contribution of each term to $\Delta G^0_{bind}$:**

To differentiate between the binding interfaces and interfacial strengths of each of the eight different systems (Figure 3a-d), we computed the Potential of Mean Force (PMF) for separating the two proteins in each system. The methodology followed in this study has been previously outlined in detail[13,30] and uses a combination of Umbrella Sampling and Adaptive Biased Force (ABF)-based simulations to compute the standard binding free energy ($\Delta G^0_{bind}$). The methodology can be divided into three main parts:

1. Calculating the PMF along the separation distance *r* using Replica Exchange Umbrella Sampling (REUS)
2. Calculating the contribution of each restraint applied to the systems using ABF (and Umbrella Sampling)
3. Using the equations described below to calculate the ($\Delta G^0_{bind}$).

The separation PMF was calculated by separating the two proteins using Steered MD and then performing Replica Exchange Umbrella Sampling (REUS) simulations to estimate the PMF with respect to separation distance *r*. Several additional restraints (four conformational, three orientational and two positional restraints) were applied during REUS simulations to ensure the configurational entropy of the systems was minimized and accelerate PMF convergence. The four configurational restraints included two applied to the two proteins' backbones $RMSD_{bb1}$ and $RMSD_{bb2}$ (shortened to $R_{1,bb}$ and $R_{2,bb}$) to restrict change of global conformation, and two applied to the two proteins' interfacial residues $RMSD_{int1}$ and $RMSD_{int2}$ (shortened to $R_{1,int}$ and $R_{2,bb}$) to maintain interfacial conformations. To define the separation and angular (orientational + positional) restraints, the two proteins in each system were each divided into three parts each, with the centers of masses of each part labelled P1, P2, P3 (protein 1) and P1', P2', P3' (protein 2) (Supporting Figure S32). The three orientational restraints were placed on angle **Θ** (P1-P1'-P2'), dihedral **Φ** (P1-P1'-P2'-P3'), and dihedral **Ψ** (P2-P1-P1'-P2'). The two positional restraints were placed on angle **θ** (P1'-P1-P2) and dihedral **φ** (P1'-P1-P2-P3). Additionally, a restraint was applied on the separation *r* (P1-P1') to maintain sampling within each window. In total, these restraints limit the configuration space the proteins could explore while separated.

To generate the starting structures for each window for REUS simulations, Steered MD Simulations (SMD) were performed. A harmonic pulling force of 20 kcal $mol^{-1}$ $Å^{-1}$ was applied to each protein, increasing the separation P1-P1' (*r*) by 25 Å over 40 ns while maintaining all other configurational restraints. From each SMD trajectory, 83 evenly spaced windows along *r* (each 0.3 Å apart) were chosen as starting structures for REUS. All windows were subjected to 10,000 steps of energy minimization while constraining the backbone, and 6 ns/window were obtained using REUS (498 ns/system) to sample along separation *r*. The trajectories were sorted to ensure the windows were consistent. All biased simulations (SMD + REUS + ABF) were performed using the CUDA-accelerated NAMD 2.14,[31,32] and the Colvars module was used to define the constraints.[33]

For every restrained collective variable (CV) $\xi_i$ using a harmonic restraint, a harmonic potential $u_{\xi_i}$ given by Eq (S4) is added to the system Hamiltonian:

$$u_{\xi_i} = \frac{1}{2} k_f \left(\xi_i - \xi_{i,ref}\right)^2 \quad \text{(S4)}$$

where $\xi_{i,ref}$ is the reference value of the CV, and $k_f$ is the force constant. The quantity of interest, $G^0_{bind}$ is given by

$$\Delta G_{bind}^{0} = -\beta^{-1} ln\left(K_{eq} C^{0}\right) \tag{S5}$$

where,

$$\beta = \frac{1}{k_B T} \qquad C^0 = \frac{1}{1661} \text{Å}^3$$

$C^0$ is the standard concentration, and $K_{eq}$ is the equilibrium constant for dissociation, where

$$K_{eq} = S^* I^* * e^{-\beta\left[\left(G_{R_{1,bb}}^{bulk} - G_{R_{1,bb}}^{site}\right) + \left(G_{R_{2,bb}}^{bulk} - G_{R_{2,bb}}^{site}\right)\right]} * e^{-\beta\left[\left(G_{R_{1,int}}^{bulk} - G_{R_{1,int}}^{site}\right) + \left(G_{R_{2,int}}^{bulk} - G_{R_{2,int}}^{site}\right)\right]} * e^{-\beta\left[\left(G_{ori}^{bulk} - G_{ori}^{site}\right) - G_{pos}^{site}\right]} \tag{S6}$$

Here, the term $S^*$ depends on the positional restraints $(\boldsymbol{\theta}, \boldsymbol{\phi})$:

$$S^* = r^{*2} \int_0^{\pi} sin\theta d\theta \int_0^{2\pi} e^{-\beta u_{pos}(\theta,\phi)}\, d\phi \tag{S7}$$

The term $I^*$ computes the separation PMF along the separation $r$:

$$I^* = \int_{site} e^{-\beta[W(r) - W(r^*)]}\, dr \tag{S8}$$

The exponential terms in Eq. (S6) are described by Eq. (S9) – (S17).[13] The next four equations outline the contribution of the conformational restraints $(R_{1,bb}, R_{2,bb}, R_{1,int}, R_{2,int})$ at the site (bound pose):

$$e^{\beta G_{R_{1,bb}}^{site}} = \frac{\int_{site} d1 \int d\mathbf{X} e^{-\beta U}}{\int_{site} d1 \int d\mathbf{X} e^{-\beta U + u_{R_{1,bb}}}} = < e^{\beta u_{R_{1,bb}}} >_{(site,U)} \tag{S9}$$

$$e^{\beta G_{R_{2,bb}}^{site}} = \frac{\int_{site} d1 \int d\mathbf{X} e^{-\beta\left(U + u_{R_{1,bb}}\right)}}{\int_{site} d1 \int d\mathbf{X} e^{-\beta\left(U + u_{R_{1,bb}} + u_{R_{2,bb}}\right)}} = < e^{\beta u_{R_{2,bb}}} >_{\left(site,U,u_{R_{1,bb}}\right)} \tag{S10}$$

$$e^{\beta G_{R_{1,res}}^{site}} = \frac{\int_{site} d1 \int d\mathbf{X} e^{-\beta\left(U + u_{R_{1,bb}} + u_{R_{2,bb}}\right)}}{\int_{site} d1 \int d\mathbf{X} e^{-\beta\left(U + u_{R_{1,bb}} + u_{R_{2,bb}} + u_{R_{1,res}}\right)}} = < e^{\beta u_{R_{1,res}}} >_{\left(site,U,u_{R_{1,bb}},u_{R_{2,bb}}\right)} \tag{S11}$$

$$e^{\beta G_{R_{2,res}}^{site}} = \frac{\int_{site} d1 \int d\mathbf{X} e^{-\beta\left(U+u_{R_{1,bb}}+u_{R_{2,bb}}+u_{R_{1,res}}\right)}}{\int_{site} d1 \int d\mathbf{X} e^{-\beta\left(U+u_{R_{1,bb}}+u_{R_{2,bb}}+u_{R_{1,res}}+u_{R_{2,res}}\right)}} = \langle e^{\beta u_{R_{2,res}}} \rangle_{\left(site,U,u_{R_{1,bb}},u_{R_{2,bb}},u_{R_{1,res}}\right)} \tag{S12}$$

Define $u_{c,all}$ :

$$u_{c,all} = u_{R_{1,bb}} + u_{R_{2,bb}} + u_{R_{1,res}} + u_{R_{2,res}}$$

For the contribution of the orientational restraints $(\mathbf{\Theta}, \mathbf{\Psi}, \mathbf{\Phi})$ at the site (bound pose):

$$e^{\beta G_{\Theta}^{site}} = \frac{\int d\mathbf{X} \boldsymbol{e}^{-\beta(\boldsymbol{U}+\boldsymbol{u}_{\boldsymbol{c},\boldsymbol{all}})}}{\int_{site} d\mathbf{1} \int d\mathbf{X} \boldsymbol{e}^{-\beta(\boldsymbol{U}+\boldsymbol{u}_{\boldsymbol{c},\boldsymbol{all}}+\boldsymbol{u}_{\Theta})}} = \langle e^{\beta u_{\Theta}} \rangle_{(site,U,u_{c,all})} \tag{S13}$$

$$e^{\beta G_{\Phi}^{site}} = \frac{\int_{site} d1 \int d\mathbf{X} e^{-\beta(U+u_{c,all}+u_{\Theta})}}{\int_{site} d1 \int d\mathbf{X} e^{-\beta(U+u_{c,all}+u_{\Theta}+u_{\Phi})}} = \langle e^{\beta u_{\Phi}} \rangle_{(site,U,u_{c,all},u_{\Theta})} \tag{S14}$$

$$e^{\beta G_{\Psi}^{site}} = \frac{\int_{site} d1 \int d\mathbf{X} e^{-\beta(U+u_{c,all}+u_{\Theta}+u_{\Phi})}}{\int_{site} d1 \int d\mathbf{X} e^{-\beta(U+u_{c,all}+u_{\Theta}+u_{\Phi}+u_{\Psi})}} = \langle e^{\beta u_{\Psi}} \rangle_{(site,U,u_{c,all},u_{\Theta},u_{\Phi})} \tag{S15}$$

We define $u_o$ as:

$$u_o = u_{\Theta} + u_{\Phi} + u_{\Psi}$$

For the contribution of the positional restraints $(\boldsymbol{\theta}\,, \boldsymbol{\phi})$ at the site (bound pose):

$$e^{\beta G_{\theta}^{site}} = \frac{\int_{site} d1 \int d\mathbf{X} e^{-\beta(U+u_{c,all}+u_o)}}{\int_{site} d1 \int d\mathbf{X} e^{-\beta(U+u_{c,all}+u_o+u_{\theta})}} = \langle e^{\beta u_{\theta}} \rangle_{(site,U,u_{c,all},u_{\Theta},u_{\Phi},u_{\Psi})} \tag{S16}$$

$$e^{\beta G_{\phi}^{site}} = \frac{\int_{site} d1 \int d\mathbf{X} e^{-\beta(U+u_{c,all}+u_o+u_{\theta})}}{\int_{site} d1 \int d\mathbf{X} e^{-\beta(U+u_{c,all}+u_o+u_{\theta}+u_{\phi})}} = \langle e^{\beta u_{\phi}} \rangle_{(site,U,u_{c,all},u_o,u_{\theta})} \tag{S17}$$

Constraints in the bulk (unbound pose):

Contribution of the conformational restraints $(R_{1,bb}, R_{2,bb}, R_{1,int}, R_{2,int})$ in the bulk (unbound pose):

$$e^{-\beta G_{R_{1,bb}}^{bulk}} = \frac{\int_{bulk} d1\, \delta(r_1 - r_1^*) \int d\mathbf{X} e^{-\beta\left(U+u_{R_{1,bb}}\right)}}{\int_{bulk} d1\, \delta(r_1 - r_1^*) \int d\mathbf{X} e^{-\beta U}} = \langle e^{-\beta u_{R_{1,bb}}} \rangle_{(bulk,U)} \tag{S18}$$

$$e^{-\beta G_{R_{2,bb}}^{bulk}} = \frac{\int_{bulk} d1\, \delta(r_1 - r_1^*) \int d\mathbf{X} e^{-\beta\left(U+u_{R_{1,bb}}+u_{R_{2,bb}}\right)}}{\int_{bulk} d1\, \delta(r_1 - r_1^*) \int d\mathbf{X} e^{-\beta\left(U+u_{R_{1,bb}}\right)}} = \langle e^{-\beta u_{R_{2,bb}}} \rangle_{\left(bulk,U,u_{R_{1,bb}}\right)} \tag{S19}$$

$$e^{-\beta G_{R_{1,res}}^{bulk}} = \frac{\int_{bulk} d1\, \delta(r_1 - r_1^*) \int d\mathbf{X} e^{-\beta\left(U+u_{R_{1,bb}}+u_{R_{2,bb}}+u_{R_{1,res}}\right)}}{\int_{bulk} d1\, \delta(r_1 - r_1^*) \int d\mathbf{X} e^{-\beta\left(U+u_{R_{1,bb}}+u_{R_{2,bb}}\right)}} = \langle e^{-\beta u_{R_{1,res}}} \rangle_{\left(bulk,U,u_{R_{1,bb}},u_{R_{2,bb}}\right)} \tag{S20}$$

$$e^{-\beta G_{R_{2,res}}^{bulk}} = \frac{\int_{bulk} d1\, \delta(r_1 - r_1^*) \int d\mathbf{X} e^{-\beta\left(U+u_{R_{1,bb}}+u_{R_{2,bb}}+u_{R_{1,res}}+u_{R_{2,res}}\right)}}{\int_{bulk} d1\, \delta(r_1 - r_1^*) \int d\mathbf{X} e^{-\beta\left(U+u_{R_{1,bb}}+u_{R_{2,bb}}+u_{R_{1,res}}\right)}} = \langle e^{-\beta u_{R_{2,res}}} \rangle_{\left(bulk,U,u_{R_{1,bb}},u_{R_{2,bb}},u_{R_{1,res}}\right)} \tag{S21}$$

The contribution of the orientational restraints $(\boldsymbol{\Theta}, \boldsymbol{\Psi}, \boldsymbol{\Phi})$ in the bulk solvent is given by –

$$e^{-\beta G_{\Theta}^{bulk}} = \frac{\int_{bulk} d1\, \delta(r_1 - r_1^*) \int d\mathbf{X} e^{-\beta(U+u_{c,all}+u_\Theta)}}{\int_{bulk} d1\, \delta(r_1 - r_1^*) \int d\mathbf{X} e^{-\beta(U+u_{c,all})}} = \langle e^{-\beta u_\Theta} \rangle_{(bulk,U,u_{c,all})} \tag{S22}$$

$$e^{-\beta G_{\Phi}^{bulk}} = \frac{\int_{bulk} d1\, \delta(r_1 - r_1^*) \int d\mathbf{X} e^{-\beta(U+u_{c,all}+u_\Theta+u_\Phi)}}{\int_{bulk} d1\, \delta(r_1 - r_1^*) \int d\mathbf{X} e^{-\beta(U+u_{c,all}+u_\Theta)}} = \langle e^{-\beta u_\Phi} \rangle_{(bulk,U,u_{c,all},u_\Theta)} \tag{S23}$$

$$e^{-\beta G_{\Psi}^{bulk}} = \frac{\int_{bulk} d1\, \delta(r_1 - r_1^*) \int d\mathbf{X} e^{-\beta(U+u_{c,all}+u_\Theta+u_\Phi+u_\Psi)}}{\int_{bulk} d1\, \delta(r_1 - r_1^*) \int d\mathbf{X} e^{-\beta(U+u_{c,all}+u_\Theta+u_\Phi)}} = \langle e^{-\beta u_\Psi} \rangle_{(bulk,U,u_{c,all},u_\Theta,u_\Phi)} \tag{S24}$$

$$e^{-\beta G_{ori}^{bulk}} = \frac{1}{8\pi^2}\int_0^{\pi} d\Theta \int_0^{2\pi} d\Phi \int_0^{2\pi} d\Psi sin\Theta e^{\beta u_{ori}(\Theta,\Phi,\Psi)} \quad \text{(S25)}$$

The expectation values defined in Equations (S9-S21) can be calculated using a relevant PMF $W(\xi)$ for that CV. For example, for the contribution due to the bulk conformational restraint ($R_{1,bb}$) (defined by equation (S18)) is given by

$$e^{-\beta G_{R_{1,bb}}^{bulk}} = < e^{-\beta u_{R_{1,bb}}} >_{bulk,U} = \frac{\int_{bulk} d\xi e^{-\beta W(\xi)} e^{-\beta u_{R_{1,bb}}(\xi)}}{\int_{bulk} d\xi e^{-\beta W(\xi)}} \quad \text{(S26)}$$

**Adaptive Biasing Force (ABF) based simulations for calculating $W(\xi)$ for the bound orientational and positional restraints** (defined by equations S13-S17):

For calculating the PMFs for the angular constraints, ABF simulations[34,35] were set up. These simulations were set to explore the phase space of the angle for which they were defined. For example, to compute $e^{\beta G_{\Psi}^{site}}$ term in equation (S12), the phase space of the dihedral Ψ at the site (bound state) is explored. As described in the equation, this simulation restrains the following variables $\xi_i - R_{1,bb}, R_{1,bb}, R_{1,bb}, R_{1,bb}, \Theta, \Phi$ at their reference positions, $\xi_{i,ref}$. The CV Ψ is then varied from $-180°$ to $+180°$ using an adaptive biasing force, to explore the phase space of the dihedral Ψ. These simulations were set for five angles each in the bound pose, for eight systems, forty simulations in total. Each simulation was run for 10 ns, using a force constant of 0.5 kcal $mol^{-1}$ $deg^{-1}$ for restraining the rest of the angles. RMSDs were constrained using a force constant of 5 kcal.$mol^{-1}$Å$^{-1}$. Each bin was sampled at least 2000 times to ensure overlap among adjacent windows and to ensure convergence. The PMFs computed by using the ABF method are all shown in Supporting Figures S9 and S10 and in Supporting Tables S1-S4.

**Adaptive Biasing Force (ABF) based simulations for calculating $W(\xi)$ for the bound/unbound conformational restraints** (defined by equations S10-S13, S19-S22):

For calculating the PMFs for the conformational constraints, ABF simulations were set up to generate starting structures. These simulations were set to explore the phase space of the RMSD for which they were defined. Each RMSD was varied from 0-4 Å using an Adaptive Biased Force, to explore the phase space of the RMSD. The trajectory generated was then used to generate windows for Umbrella Sampling, and a series of 21 windows 0.2 Å apart were used. Each window was sampled for 5ns each, with a total of 105 ns simulated for all windows in each run. These simulations for eight RMSDs per system (four bound + four unbound), with a total of 64 runs. The force constant used to constrain each window was 5 kcal.$mol^{-1}$ Å$^{-1}$. The PMFs computed by using the ABF method are all shown in Supporting Figures S11 and S12 for the bound state (site), Figures S33 and S34 for the unbound state (bulk), and in Supporting Tables S1-S4.

**Calculating separation PMF using REUS:**

The separation PMFs were calculated using the multistate Bennett acceptance ratio (MBAR) method, using its implementation in the python package pymbar.[36] The trajectories obtained from REUS were sorted to ensure the samples were uncorrelated (Supporting Tables S1-S4). CPPTRAJ[37] was used for extracting frames and calculating RMSDs. VMD[38] and open-source Pymol[16] were used for visualization. Various python programming libraries were used for handling data – mdtraj[39] was used for trajectory post-processing and computing observables. numpy[40] and scipy[41] were used for numerical computations. Data were organized using pandas.[42] Plots were made using matplotlib[43] and seaborn.[44] Standard python libraries random, math, os, subprocess were used in in-house scripts for facilitating computations.

**Error analysis:**

The errors were computed for every observable calculated (Supporting Tables S1-S4). All PMFs were calculated using pymbar, a module that also outputs the errors in energies. Using these errors in PMF as basis, errors in the free energy contribution of each term were computed, using bootstrapping.

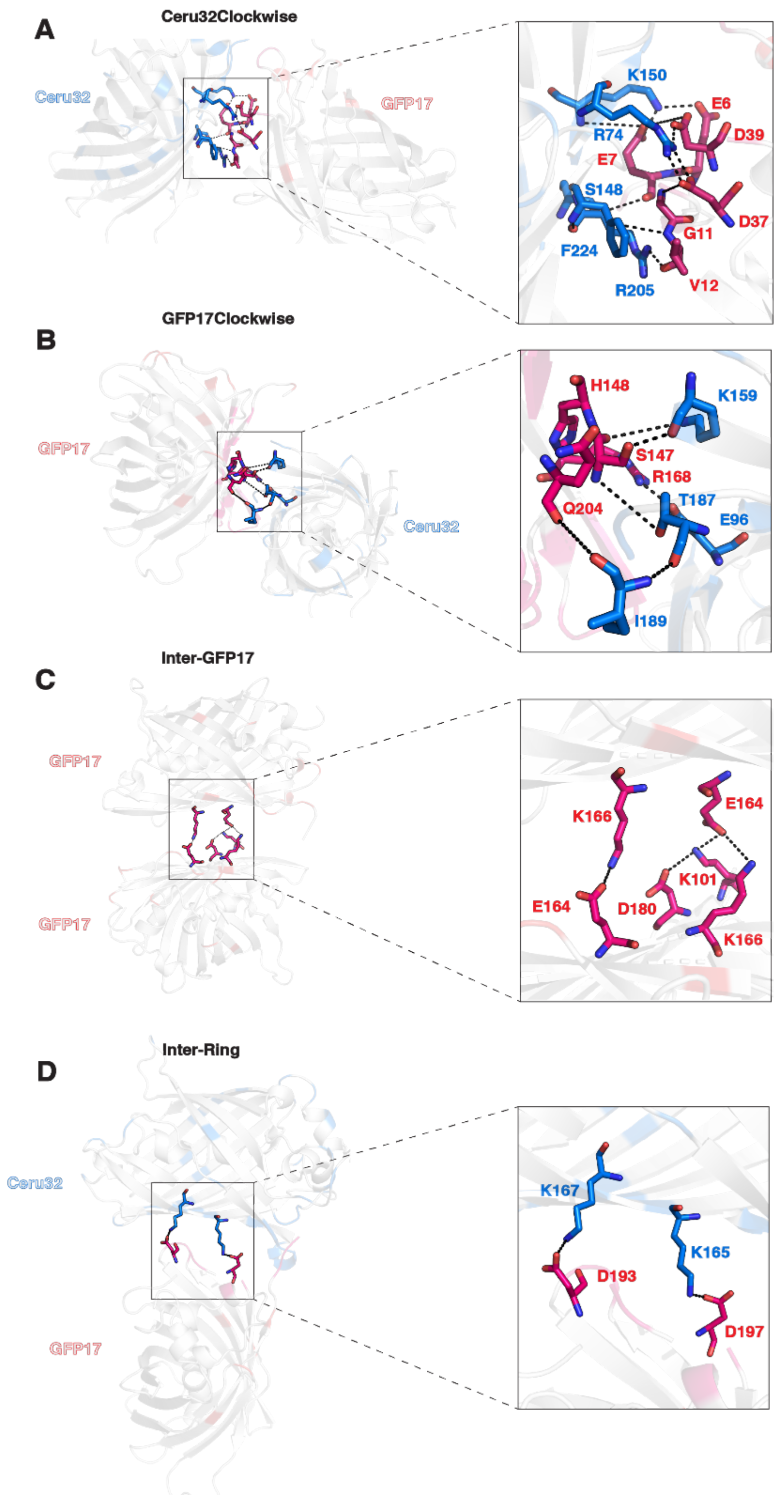


**Figure S1.** The four primary protein-protein interfaces in the assembled 16-mer structure described in Simon et al. for the Ceru+32o / GFP-17 hexadecamer. (A) Ceru32Clockwise, (B) GFP17Clockwise, (C) InterGFP17, (D) Inter-Ring. The images on the right show the interfacial residues. (Ceru+32o – Blue, GFP-17 – Red).

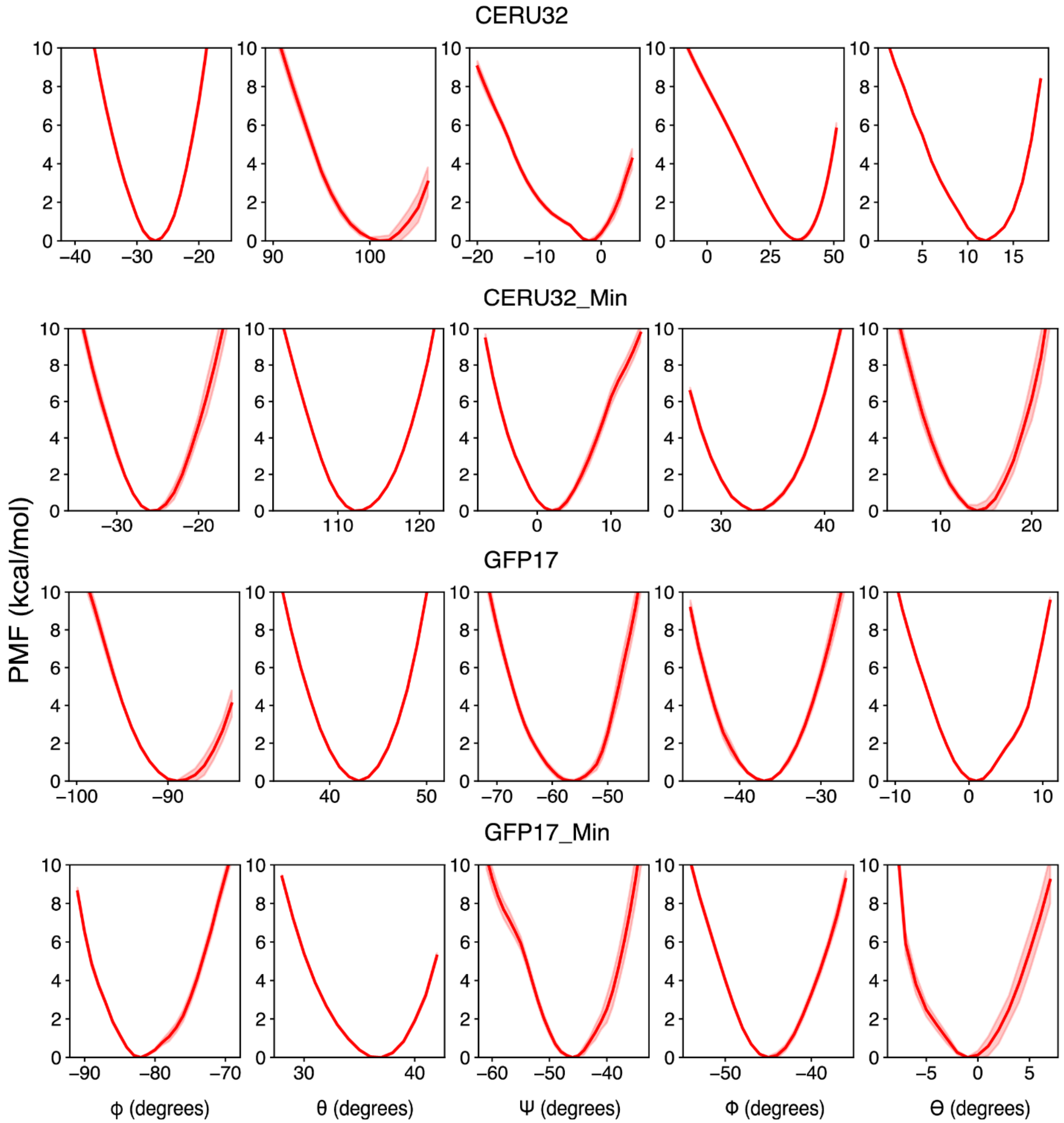


**Figure S2.** PMFs of the intra-planar systems' contributions of the orientational and positional constraints in the bound state (site).

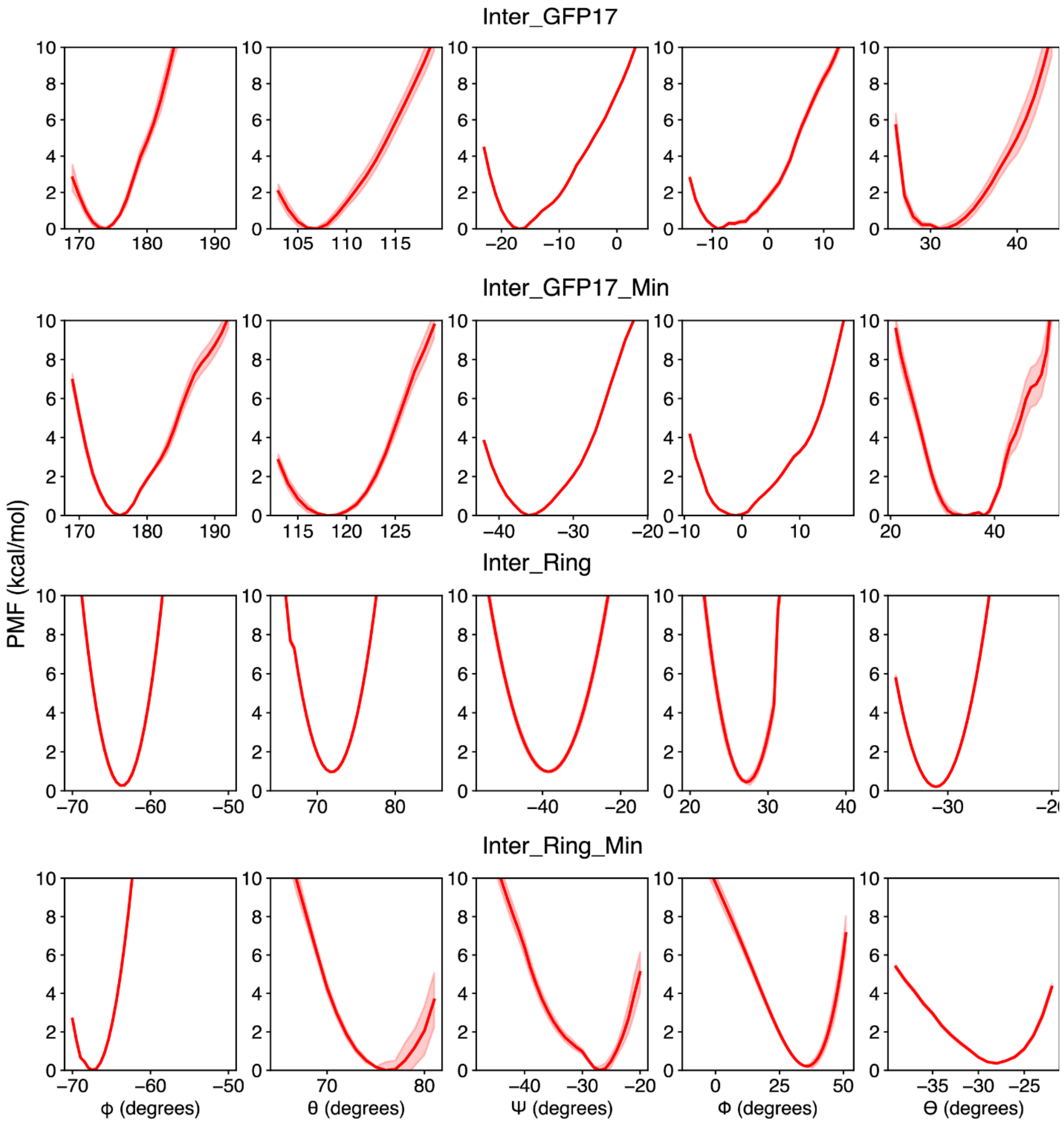


**Figure S3.** PMFs of the inter-planar systems' contributions of the orientational and positional constraints in the bound state (site).

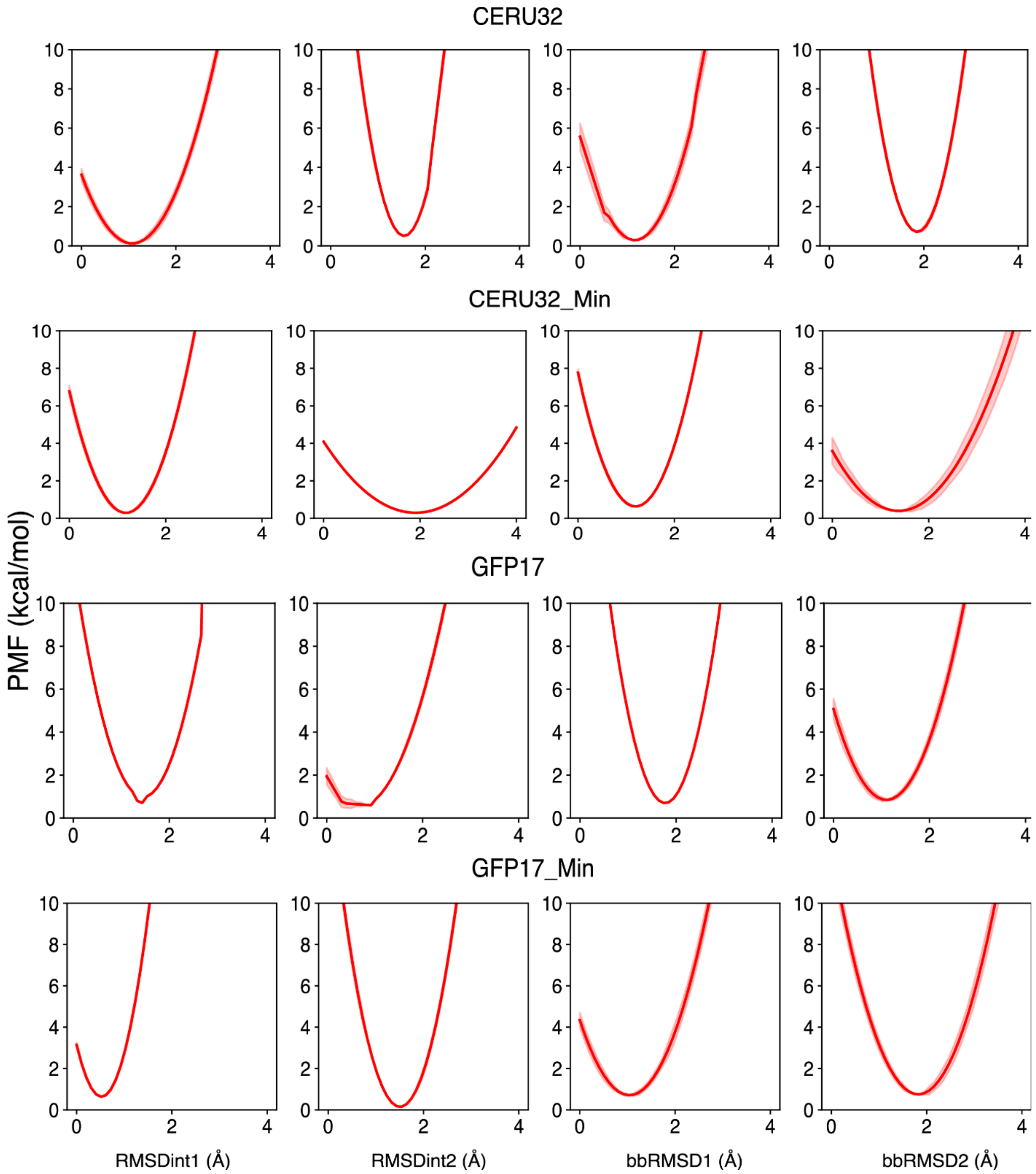


**Figure S4.** PMFs of the intra-planar systems' contributions of the conformational constraints in the bound state (site).

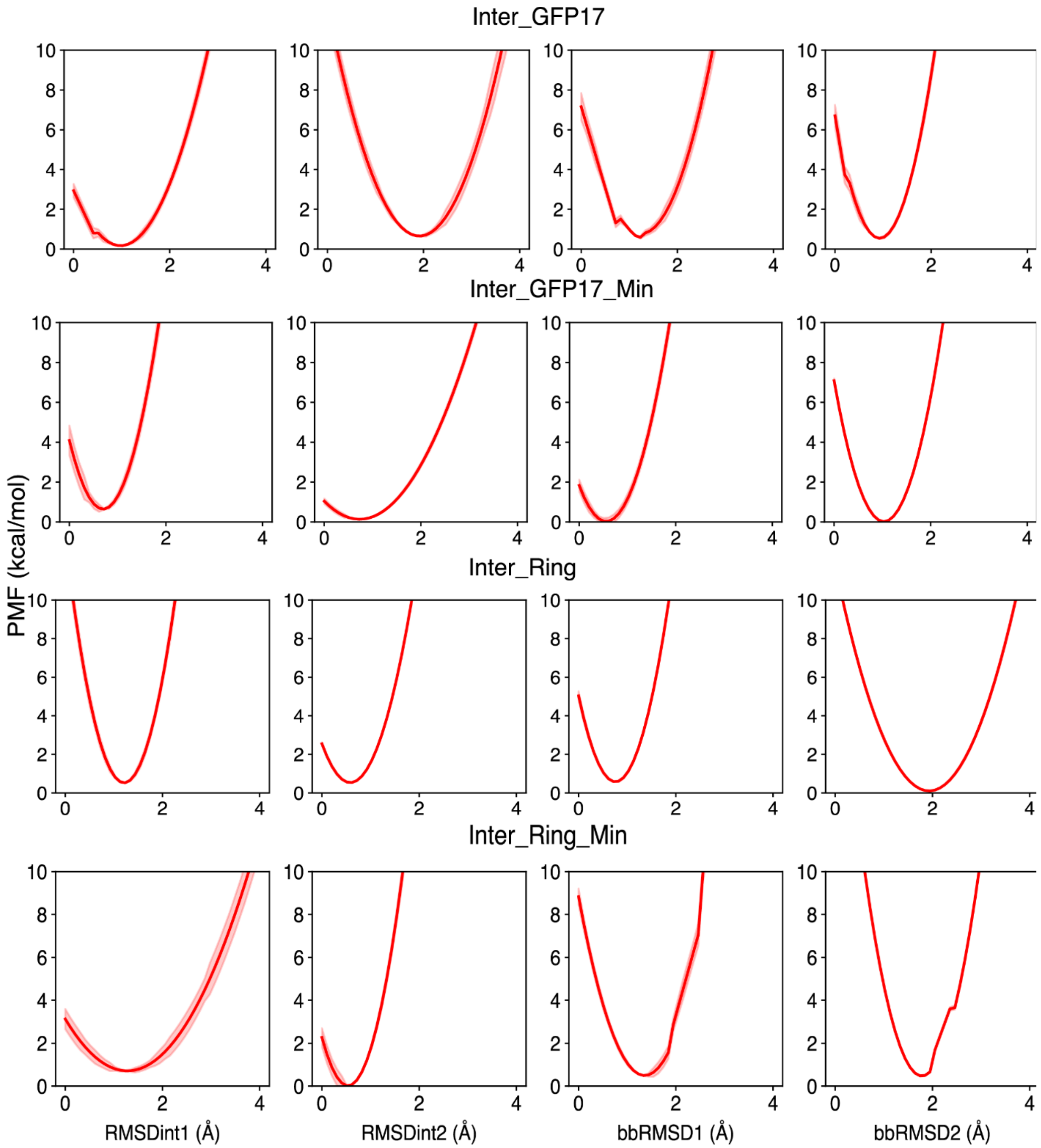


**Figure S5.** PMFs of the inter-planar systems' contributions of the conformational constraints in the bound state (site).

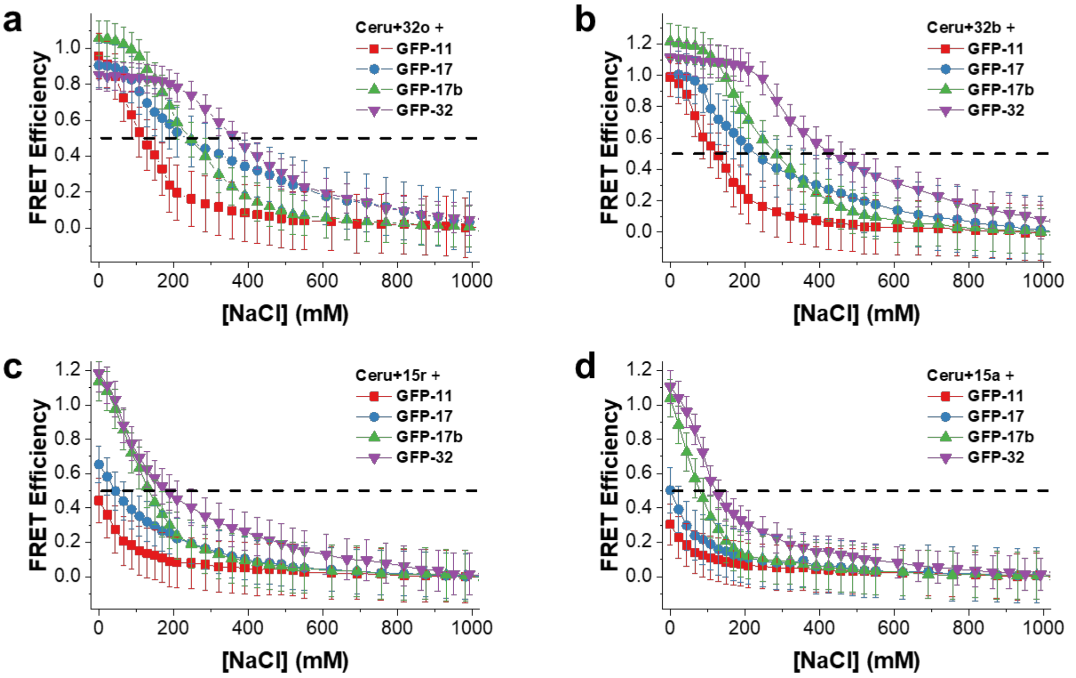


**Figure S6.** FRET efficiency versus NaCl concentration for each of the negatively charged proteins and Ceru+32o (a), Ceru+32b (b), Ceru+15r (c), and Ceru+15a (d) at pH 7.4. The concentration of each protein is 0.1 mg/mL.

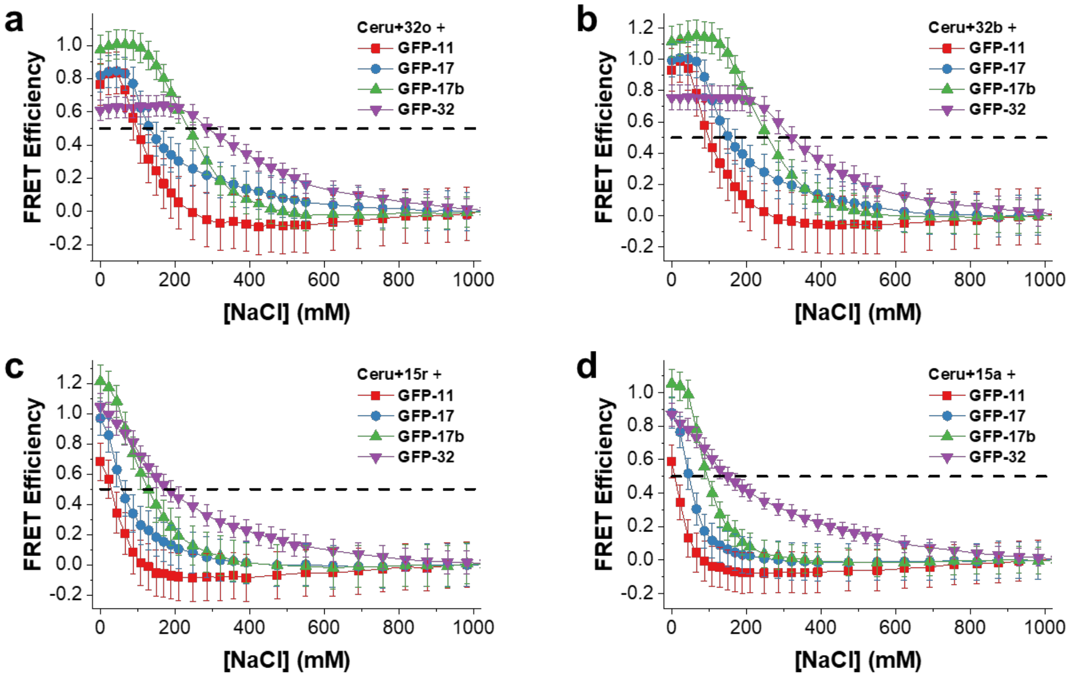


**Figure S7.** FRET efficiency versus NaCl concentration for each of the negatively charged proteins and Ceru+32o (a), Ceru+32b (b), Ceru+15r (c), and Ceru+15a (d) at pH 6. The concentration of each protein is 0.1 mg/mL.

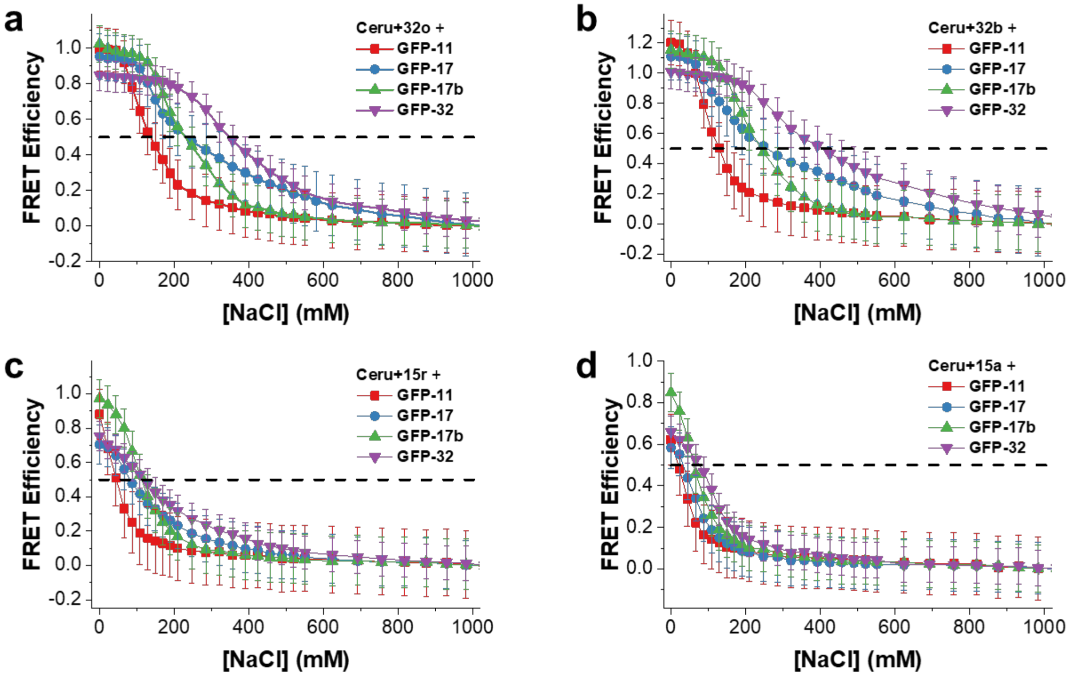


**Figure S8.** FRET efficiency versus NaCl concentration for each of the negatively charged proteins and Ceru+32o (a), Ceru+32b (b), Ceru+15r (c), and Ceru+15a (d) at pH 9. The concentration of each protein is 0.1 mg/mL.

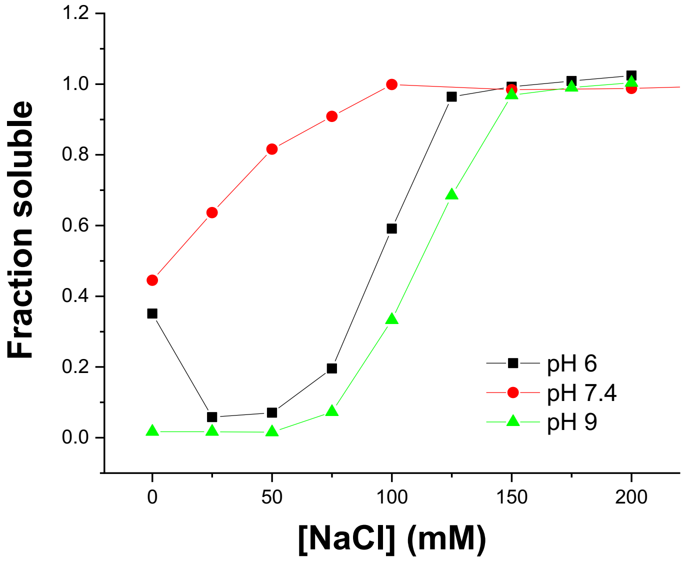


**Figure S9.** Fraction of GFP-17 / Ceru+32o assemblies that remained in solution after centrifugation at 18,000g for 2 min. Fluorescence intensity from GFP-17 was measured before and after centrifugation to determine the fraction soluble.

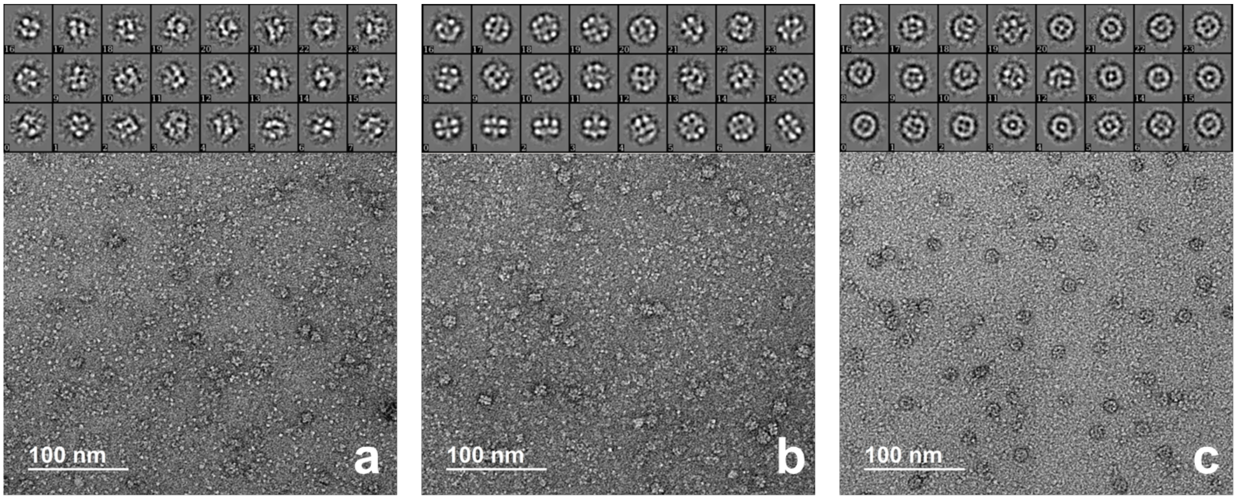


**Figure S10.** Representative negative stain TEM images and particle class averages of Ceru+32o / GFP-17 assemblies in 150 mM NaCl at: (a) pH 6 (n = 1,178 particles); (b) pH 7.4 (n = 1,558 particles); and (c) pH 9 (n = 1,212 particles). Protomers at pH 6 are much more heterogeneous than those at pH 7.4 and pH 9.

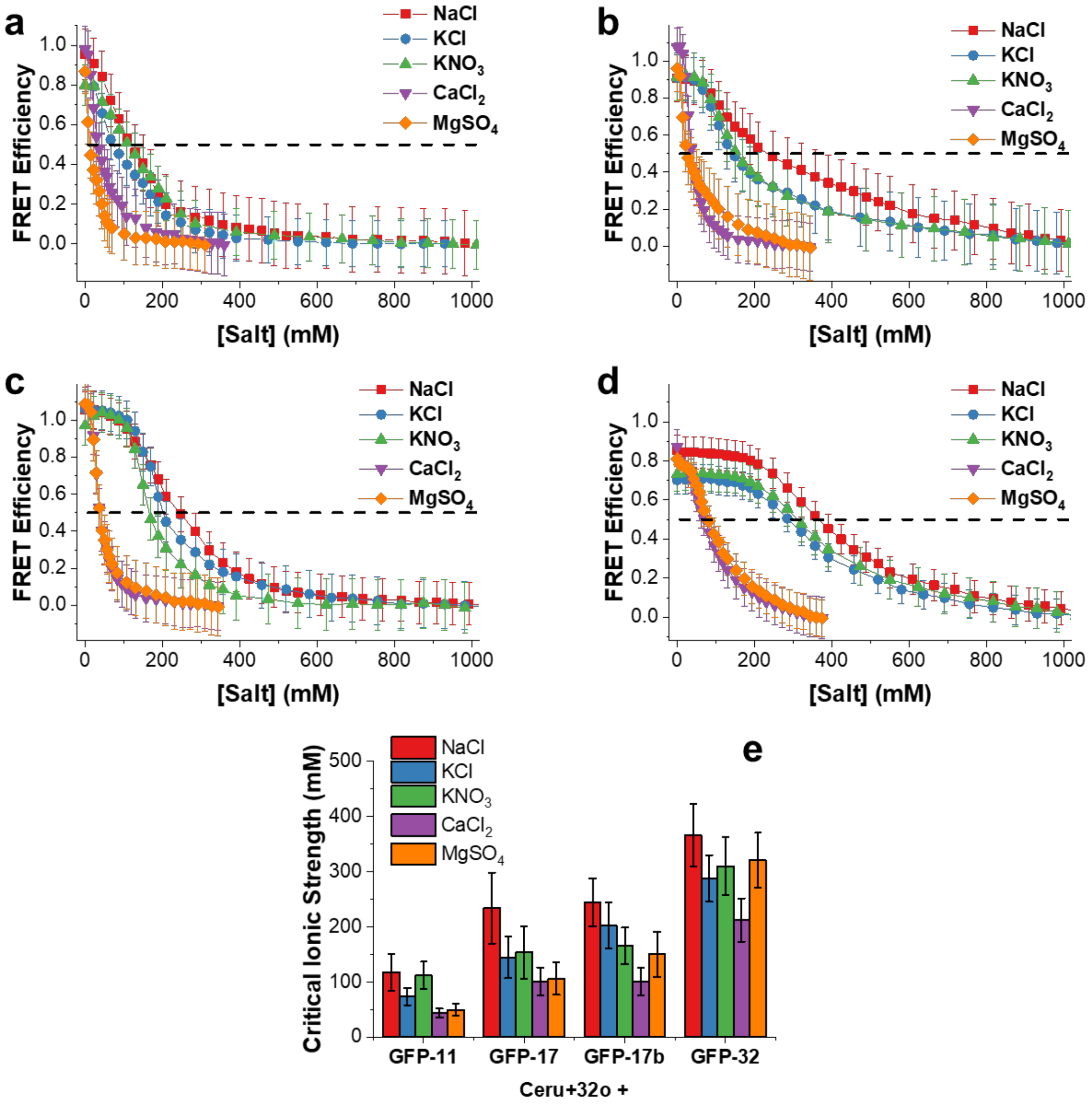


**Figure S11.** FRET efficiency versus salt concentration (NaCl, KCl, $KNO_3$, $CaCl_2$, and $MgSO_4$) between Ceru+32o and (a) GFP-11, (b) GFP-17, (c) GFP-17b, and (d) GFP-32 at pH 7.4. The concentration of each protein is 0.1 mg/mL. Dashed lines represent the FRET efficiency = 0.5. (e) The critical ionic strength for each protein combination in each salt solution. Error bars represent ±1 standard deviation.

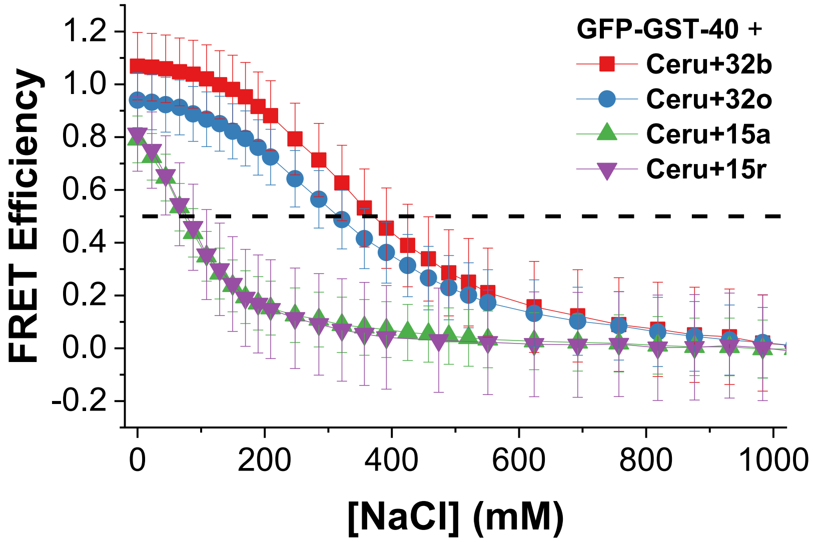


**Figure S12.** FRET efficiency versus NaCl concentration between negatively supercharged GFP-GST-40 fusion protein and each of the positively supercharged Ceru variants at pH 7.4. The concentration of each protein is 4 µM.

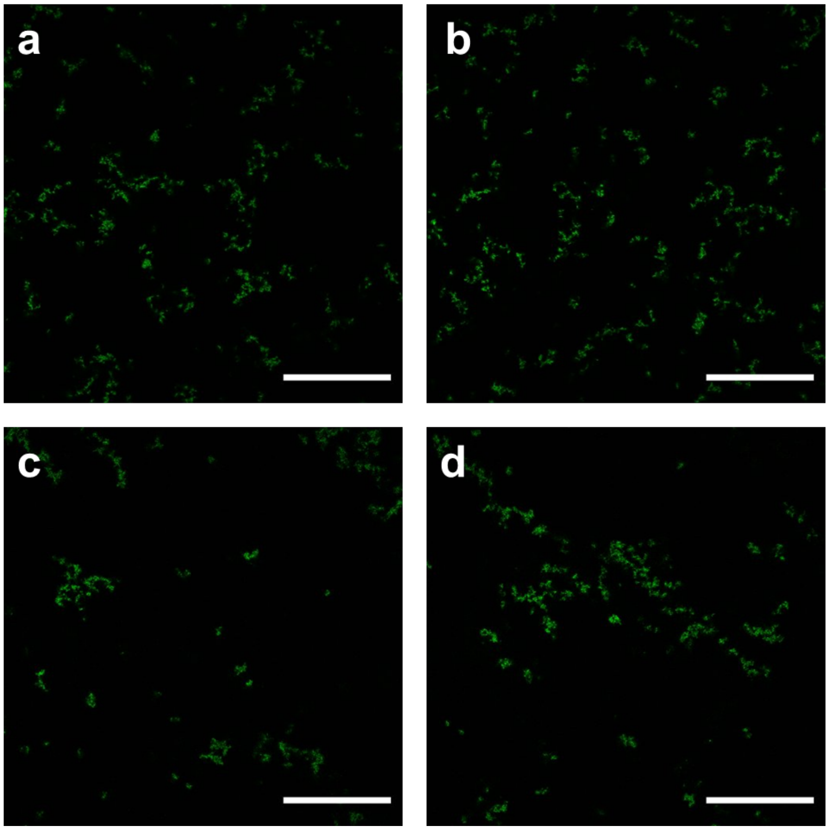


**Figure S13.** Confocal microscopy images of assemblies at 0 mM NaCl between: (a) Ceru+32o / GFP-17, (b) Ceru+32b / GFP-17, (c) Ceru+32o / GFP-17b, (d) and Ceru+32b / GFP-17b. GFP was excited using the 488 nm laser line, and fluorescence intensity from 500-600 nm was imaged. Scale bars represent 50 µm.

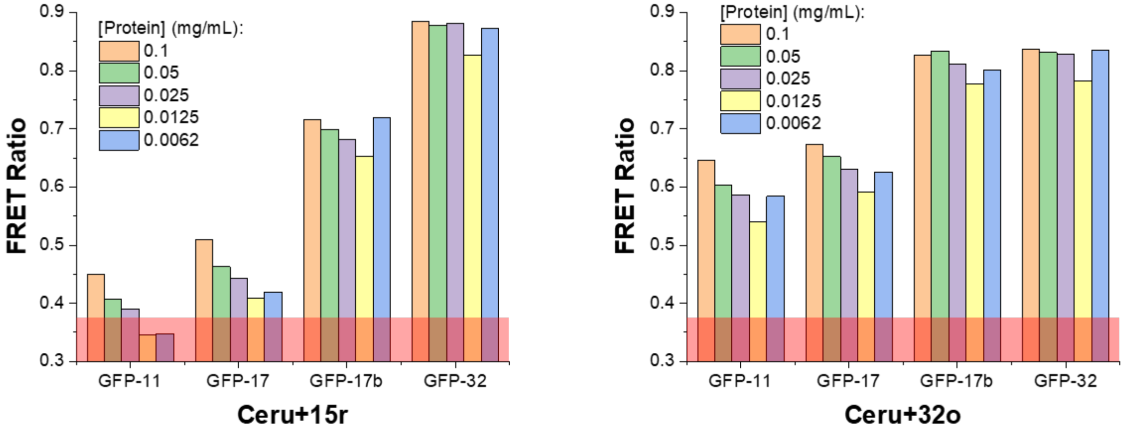


**Figure S14.** FRET ratio between each of the negatively supercharged variants and (a) Ceru+15r or (b) Ceru+32o at different equimolar concentrations in 0 M NaCl solution. Shaded red regions represent the FRET ratio for non-interacting proteins. FRET ratio decreases at lower concentrations for weakly associating proteins and remains constant at all concentrations for strongly associating proteins.

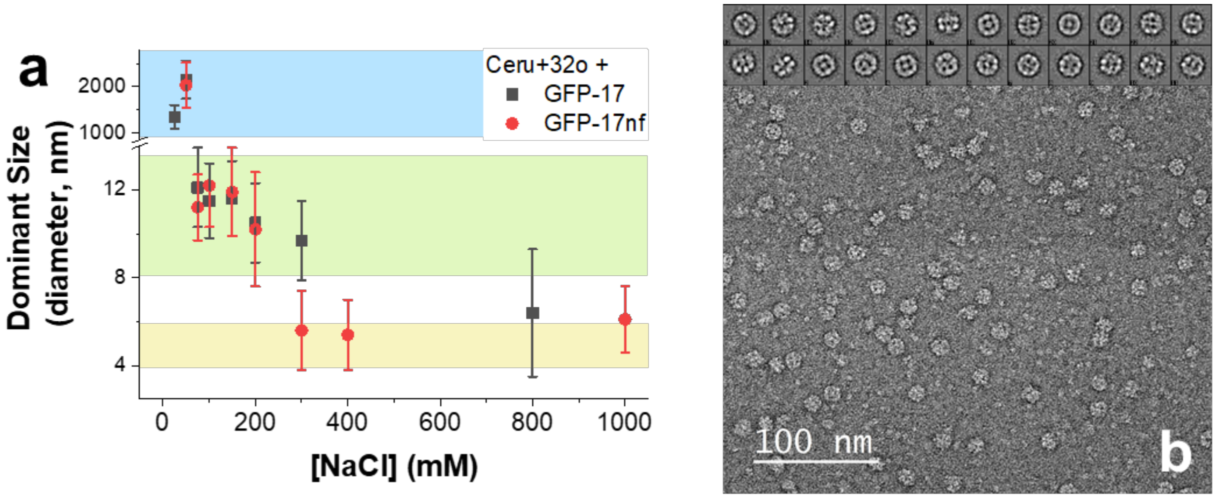


**Figure S15.** Comparison of assembly between Ceru+32o / GFP-17 and Ceru+32o / GFP-17nf. (a) Dominant sizes (measured by DLS) between the two protein combinations are similar at all NaCl concentrations and show evidence of protomer formation at 50-200 mM NaCl. (b) TEM images and class averages ($n$ = 2,743 particles) show evidence of protomer formation for Ceru+32o / GFP-17nf at 75 mM NaCl (pH 7.4), which is consistent with results from assemblies between Ceru+32o / GFP-17 (Figure 5c).

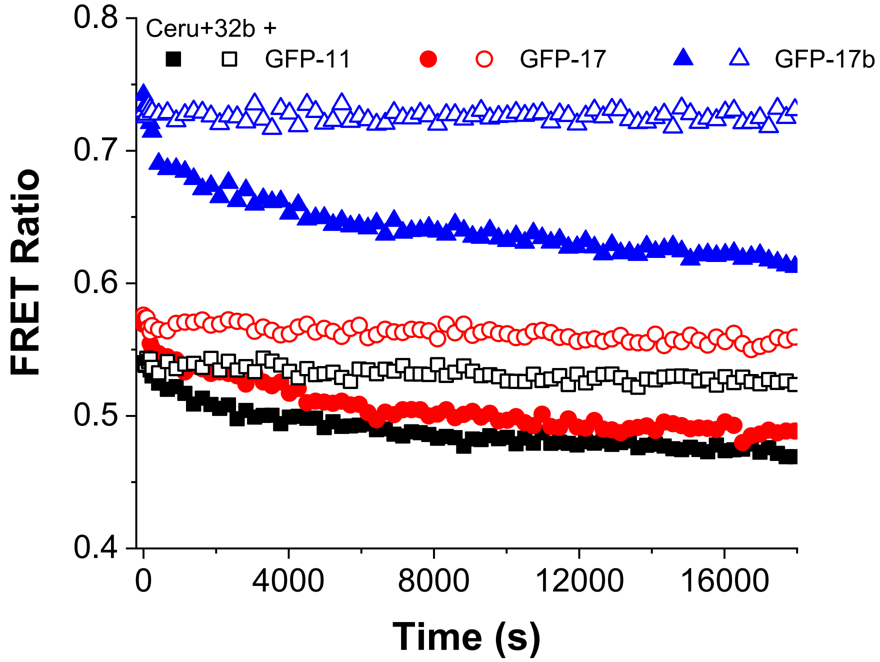


**Figure S16.** Transient FRET ratio when the negatively charged non-fluorescent analog or buffer is added to solution for different protein combinations at 0 mM NaCl (pH 7.4). The initial concentration of each fluorescent protein is 0.01 mg/mL and the non-fluorescent analog was added to 0.1 mg/mL. Adding buffer did not cause FRET ratio to change.

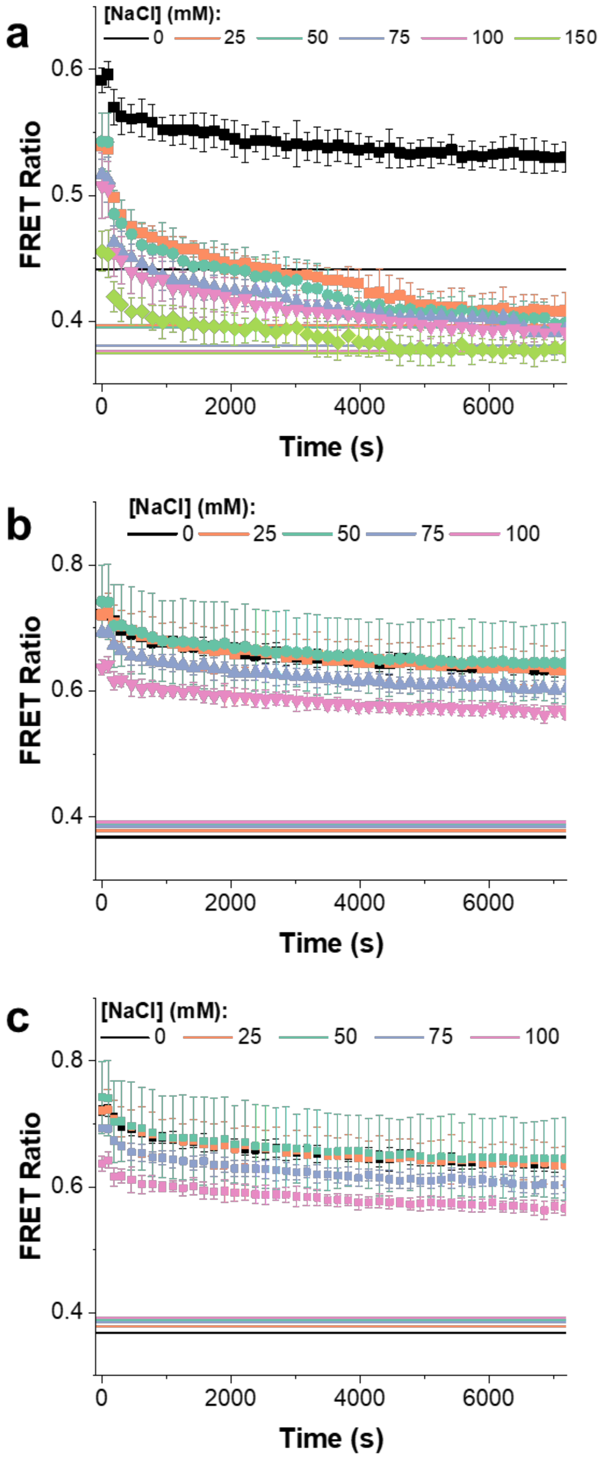


**Figure S17.** Transient FRET ratio between (a) Ceru+32b / GFP-17, (b) Ceru+32o / GFP-17b, and (c) Ceru+32b / GFP-17b after the corresponding non-fluorescent negatively charged analog (GFP-17nf or GFP-17bnf) is added to solution at different concentrations of NaCl. The solid lines represent the final FRET ratio with all GFP-17nf or GFP-17bnf incorporated at each salt concentration.

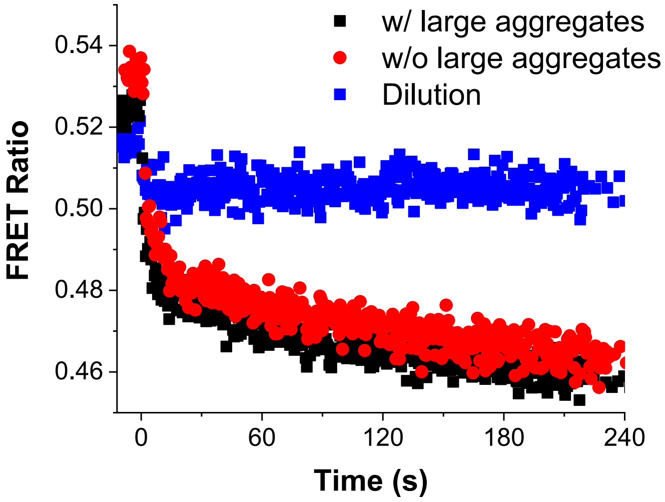


**Figure S18.** Transient FRET ratio between Ceru+32o / GFP-17 at 75 mM NaCl when 10x GFP-17nf (or buffer solution) are added. Any large aggregates were removed by centrifugation (18,000 g for 2 min). The initial fast drop in FRET ratio was present both with and without large aggregates and was slower than the dilution timescale. This suggests the long timescale changes in FRET ratio arise from dynamic subunit exchange of protomers, and the fast change in FRET ratio arises from displacement of weakly associating proteins from the protomers.

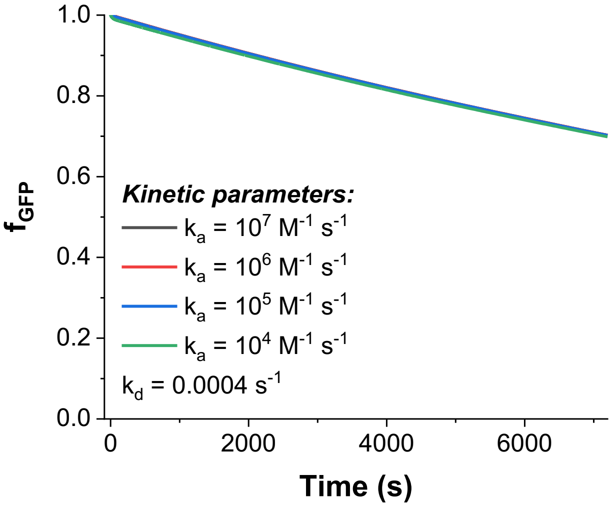


**Figure S19.** Simulated fraction of fluorescent GFP remaining in protomers with different reassembly rate constants $k_a$. Dissociation was rate limiting, and the simulation was not sensitive to the precise value of the reassembly rate constant $k_a$.

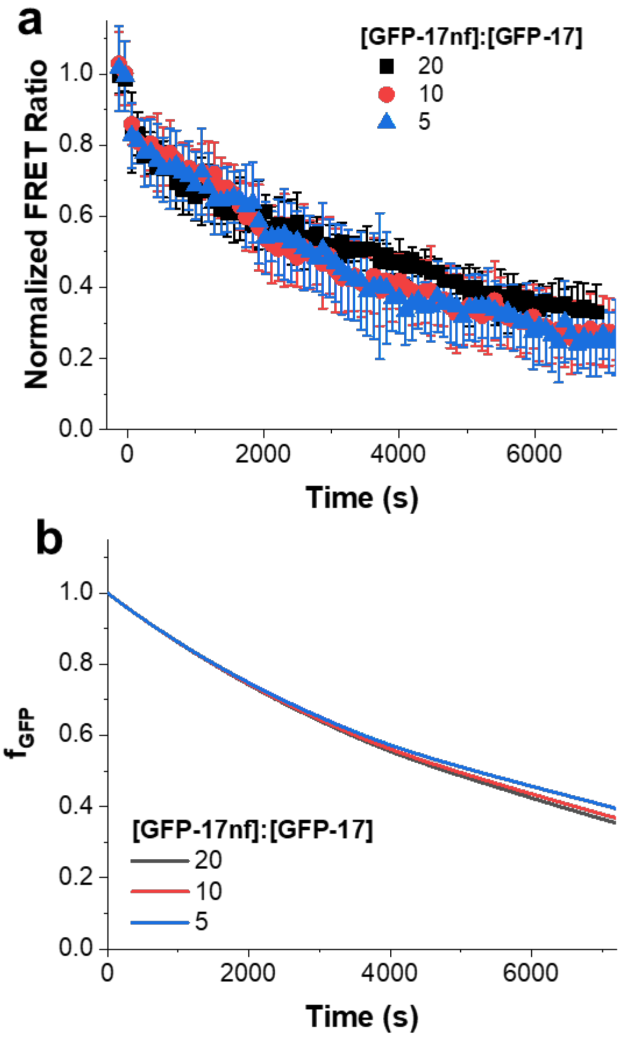


**Figure S20.** Similar exchange kinetics are observed both experimentally and computationally when different concentrations of non-fluorescent analog are added to solution. (a) Changing normalized FRET ratio when different concentrations of non-fluorescent analog are added Ceru+32o / GFP-17 assembly at 75 mM NaCl (pH 7.4). The concentration of GFP-17 / Ceru+32o is constant (0.01 mg/mL). (b) Simulated fraction of fluorescent GFP remaining in the protomers when different concentrations of non-fluorescent analog are incorporated into protomers ($k_d$ = 0.0012 $s^{-1}$, $k_a$ = $10^6$ $M^{-1}$ $s^{-1}$).

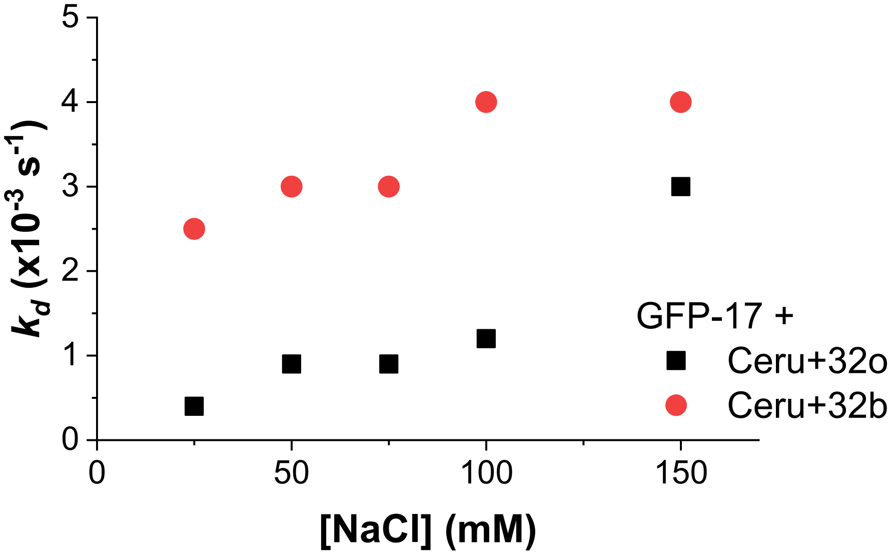


**Figure S21.** Modeled dissociation rate constants $k_d$ that best fit the experimental data for protomer assembly between Ceru+32o / GFP-17 (black squares) and Ceru+32b / GFP-17 (red circles).

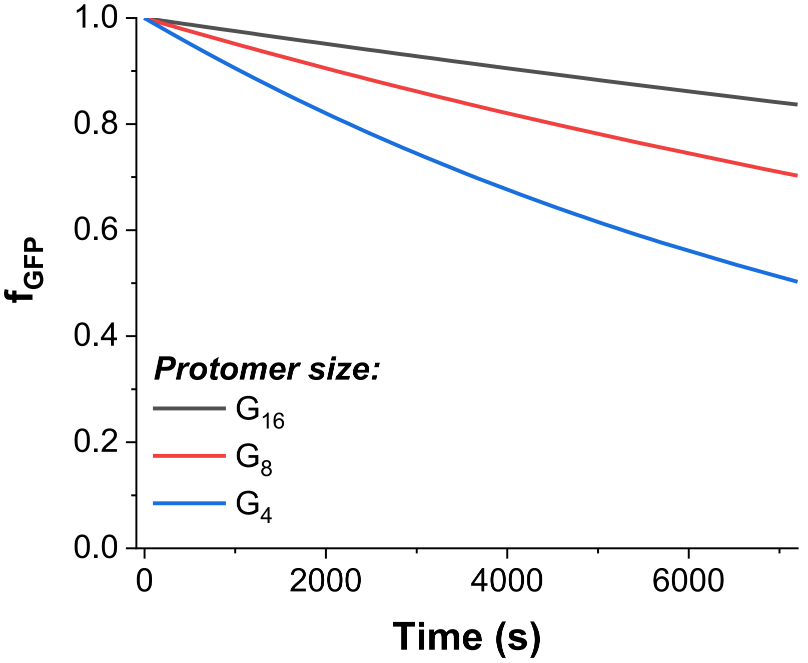


**Figure S22.** Simulated fraction of fluorescent GFP remaining in protomers with different sized protomers. The total assembled fluorescent protein (0.4 μM), non-fluorescent protein concentration (4 μM), and rate constants ($k_d$ = 0.0004 s$^{-1}$ and $k_a$ = 10$^6$ M$^{-1}$ s$^{-1}$) are the same for each simulation. Smaller sized protomers exchange faster.

```
         1          11         21         31         41         51         61
GFP(wt)   MSKGEELFT GVVPILVELD GDVNGHKFSV RGEGEGDATN GKLTLKFICT TGKLPVPWPT LVTTLTWGVQ
Ceru+15r MASKGEELFK GVVPILVELK GDVNGHKFKV RGEGEGDAKN GKLRLKFICT TGKLPVPWPT LVTTLTWGVQ
Ceru+15a MASKGEKLFT GVVPILVELD GDVNGHKFSV RGEGEGDATK GKLTLKFICT TGKLPVPWPT LVTTLTWGVQ
Ceru+32o MASKGERLFR GKVPILVELK GDVNGHKFSV RGKGKGDATN GKLTLKFICT TGKLPVPWPT LVTTLTWGVQ
Ceru+32b MASKGEKLFK GVVPILVELK GKVNGHKFKV RGEGEGDAKN GKLRLKFICT TGKLPVPWPT LVTTLTWGVQ

         71         81         91         101        111        121        131
GFP(wt)  CFSRYPDHMK RHDFFKSAMP EGYVQERTIS FKDDGTYKTR AEVKFEGDTL VNRIELKGID FKEDGNILGH
Ceru+15r CFSRYPDHMK RHDFFKSAMP EGYVQERTIS FKDDGTYKTR AEVKFEGDTL VNRIELKGIK FKKDGNILGH
Ceru+15a CFSRYPKHMK RHDFFKSAMP EGYVQERTIS FKKDGTYKTR AEVKFEGKTL VNRIELKGIK FKKKGNILGH
Ceru+32o CFARYPKHMK RHDFFKSAMP KGYVQERTIS FKKDGTYKTR AEVKFEGRTL VNRIKLKGRD FKEKGNILGH
Ceru+32b CFSRYPDHMK RHDFFKSAMP EGYVQKRKIS FKKDGTYKTR AEVKFEGKTL VNRIKLKGIK FKKDGNILGH

         141        151        161        171        181        191        201
GFP(wt)  KLEYNFNSHN VYITADKQKN GIKANFKIRH NVEDGSVQLA DHYQQNTPIG DGPVLLPDNH YLSTQSVLSK
Ceru+15r KLEYNGISDK VYITADKKKN GIKAKFKIRH NVKDGSVQLA DHYQQNTPIG DGPKLLPRKH YLSTQSVLSK
Ceru+15a KLEYNGISDN VYITADKKKN GIKANFKIRH NVEKGSVQLA DHYQQNTPIG KGPVLLPDNH YLSTQSVLSK
Ceru+32o KLRYNGISDK VYITADKRKN GIKAKFKIRH NVKDGSVQLA DHYQQNTPIG RGPVLLPRNH YLSTRSVLSK
Ceru+32b KLRYNGISDK VYIKADKKKN GIKAKFKIRH NVKDGSVQLA DHYQQNTPIG DGPKLLPRKH YLSTKSVLRK

         211        221        231        241
GFP(wt)  DPNEKRDHMV LLEFVTAAGI THGMDELYKG SHHHHHH
Ceru+15r DPNEKRDHMV LLEFVTAAGI TKGMDELYKL EHHHHHH
Ceru+15a DPKEKRDHMV LLEFVTAAGI THGMDELYKL EHHHHHH
Ceru+32o DPKEKRDHMV LLEFVTAAGI KHGRDERYKL EHHHHHH
Ceru+32b DPNEKRDHMV LLEFVTAAGI TKGMDELYKL EHHHHHH
```

**Figure S23.** Amino acid sequences of positively supercharged proteins used in this work. Blue highlighted amino acids were mutated (compared to wild-type superfolder GFP) to incorporate a positive charge. Other bold amino acids were mutated for stability purposes. To create non-fluorescent variants, yellow highlighted amino acids were mutated to GGG.

| | 1 | 10 | 20 | 30 | 40 | 50 | 60 |
|---|---|---|---|---|---|---|---|
| GFP(wt) | MSKGEELFT | GVVPILVELD | GDVNGHKFSV | RGEGEGDATN | GKLTLKFICT | TGKLPVPWPT | LVTTLTYGVQ |
| GFP-11 | MSKGEELFT | GVVPILVELD | GDVNGHKFSV | RGEGEGDA**D**N | GKL**D**LKFICT | TGKLPVPWPT | LVTTLTYGVQ |
| GFP-17 | MSKGEELFT | GVVPILVELD | GDVNGHKFSV | RGEGEGDA**D**N | GKL**D**LKFICT | TGKLPVPWPT | LVTTLTYGVQ |
| GFP-17b | MSKGEELF**D** | GVVPILVELD | GDVNGH**E**FSV | RGEGEGDATN | G**E**LTLKFICT | TG**E**LPVPWPT | LVTTLTYGVQ |
| GFP-32 | MSKGEELF**E** | GVVPILVELD | GDVNGHKF**E**V | RGEGEGDA**D**N | GKL**D**LKFICT | TG**E**LPVPWPT | LVTTLTYGVQ |

| | 70 | 80 | 90 | 100 | 110 | 120 | 130 |
|---|---|---|---|---|---|---|---|
| GFP(wt) | CFSRYPDHMK | RHDFFKSAMP | EGYVQERTIS | FKDDGTYKTR | AEVKFEGDTL | VNRIELKGID | FKEDGNILGH |
| GFP-11 | CFSRYPDHMK | RHDFFKSAMP | EGYVQERTIS | FKDDGTYKTR | AEVKFEGDTL | VNRIELKGID | FKEDGNILGH |
| GFP-17 | CFSRYPDHMK | **E**HDFFKSAMP | EGYVQERTIS | FKDDGTYKTR | AEVKFEGDTL | VNRIELKGID | FKEDGNILGH |
| GFP-17b | CFSRYPDHMK | RHDFFKSAMP | EGYVQERTIS | F**E**DDGTYKTR | AEVKFEGDTL | VNRIELKGID | FKEDGNILGH |
| GFP-32 | CFSRYPDHMK | **E**HDFFKSAMP | EGYVQERTIS | F**D**DDGTY**E**TR | AEVKFEGDTL | VNRIELKGID | FKEDGNILGH |

| | 140 | 150 | 160 | 170 | 180 | 190 | 200 |
|---|---|---|---|---|---|---|---|
| GFP(wt) | KLEYNFNSHN | VYITADKQKN | GIKANFKIRH | NVEDGSVQLA | DHYQQNTPIG | DGPVLLPDNH | YLSTQSVLSK |
| GFP-11 | KLEYNFNSHN | VYITADK**E**KN | GIKANFKIRH | NVEDGSVQLA | DHYQQNTPIG | DGP**D**LLPD**E**H | YLSTQSVLSK |
| GFP-17 | KLEYNFNSH**E** | VYITAD**DE**KN | GIKA**E**FKIRH | NVEDGSVQLA | DHYQQNTPIG | DGP**D**LLPD**E**H | YLSTQSVLSK |
| GFP-17b | KLEYNFNSHN | VYITADKQKN | GIKANFKIRH | NVEDGSVQ**E**A | DHYQQNTPIG | DGPVLLPDNH | YLSTQSVLSK |
| GFP-32 | KLEY**D**FNSH**E** | VYI**E**AD**DE**KN | GIKA**E**FKI**E**H | NVEDGSVQLA | DHYQQNTPIG | DGP**D**LLPD**E**H | YLSTQSVLSK |

| | 210 | 220 | 230 | 240 |
|---|---|---|---|---|
| GFP(wt) | DPNEKRDHMV | LLEFVTAAGI | THGMDELYKG | SHHHHHH |
| GFP-11 | DPNEKRDHMV | LLEFVTAAGI | T**E**GHHHHHHHH | |
| GFP-17 | DPNEKRDHMV | LLEFVTA**D**GI | T**E**GHHHHHHHH | |
| GFP-17b | DP**D**E**E**RDHMV | LLEFVTAAGI | THGHHHHHH | |
| GFP-32 | DPNE**E**RDHMV | LLEFVTA**D**GI | T**E**GHHHHHHHH | |

**Figure S24.** Amino acid sequences of negatively supercharged proteins used in this work. Bold red amino acids were mutated (compared to wild-type superfolder GFP) to incorporate a negative charges. To create non-fluorescent variants, the yellow highlighted amino acids were mutated to GGG.

```
             1          11         21         31         41         51         61
GFP(wt)       MSKGEELFT GVVPILVELD GDVNGHKFSV RGEGEGDATN GKLTLKFICT TGKLPVPWPT LVTTLTYGVQ
GFP_min       MSKGEELFT GVVPILVELD GDVNGHKFSV RGEGEGDADN GKLTLKFICT TGKLPVPWPT LVTTLTYGVQ
Ceru_min     MASKGEELFT GVVPILVELD GDVNGHKFSV RGEGEGDATN GKLTLKFICT TGKLPVPWPT LVTTLTWGVQ

             71         81         91         101        111        121        131
GFP(wt)      CFSRYPDHMK RHDFFKSAMP EGYVQERTIS FKDDGTYKTR AEVKFEGDTL VNRIELKGID FKEDGNILGH
GFP_min      CFSRYPDHMK RHDFFKSAMP EGYVQERTIS FKDDGTYKTR AEVKFEGDTL VNRIELKGID FKEDGNILGH
Ceru_min     CFARYPDHMK RHDFFKSAMP EGYVQERTIS FKDDGTYKTR AEVKFEGDTL VNRIKLKGID FKEDGNILGH

             141        151        161        171        181        191        201
GFP(wt)      KLEYNFNSHN VYITADKQKN GIKANFKIRH NVEDGSVQLA DHYQQNTPIG DGPVLLPDNH YLSTQSVLSK
GFP_min      KLEYNFNSHN VYITADDEKN GIKAEFKIRH NVEDGSVQLA DHYQQNTPIG DGPDLLPDNH YLSTQSVLSK
Ceru_min     KLEYNGISDK VYITADKQKN GIKAKFKIRH NVEDGSVQLA DHYQQNTPIG RGPVLLPDNH YLSTRSVLSK

             211        221        231        241
GFP(wt)      DPNEKRDHMV LLEFVTAAGI THGMDELYKG SHHHHHH
GFP_min      DPNEKRDHMV LLEFVTAAGI THGHHHHHHH
Ceru_min     DPKEKRDHMV LLEFVTAAGI KHGMDELYKG SHHHHHH
```

**Figure S25.** Amino acid sequences of the minimally mutated variants used in this work. Bold red amino acids were mutated (compared to wild-type superfolder GFP) to incorporate negative charges, and bold blue amino acids were mutated to incorporate positive charges. The green highlighted and italicized amino acids were previously identified to be involved in the protomer interface.[1]

```
1        10        20        30        40        50        60        70        80
MGHHHHHHSKGEELFTGVVPILVELDGDVNGHKFSVSGEGEGDATYGKLTLKFICTTGKLPVPWPTLVTTLTYGVQCFSRYPDHMKRHD
90       100       110       120       130       140       150       160       170
FFKSAMPEGYVQERTISFKDDGNYKTRAEVKFEGDTLVNRIELKGIDFKEDGNILGHKLEYNYNSHNVYITADKQKNGIKANFKIRHNIE
180      190       200       210       220       230       240       250       260
DGSVQLADHYQQNTPIGDGPVLLPDNHYLSTQSALSKDPNEKRDHMVLLEFVTAAGITHGMDELYKKLAAALGGPPYTITYFPVRGRCEA
270      280       290       300       310       320       330       340       350
MRMLLADQDQSWEEEVVTMETWPPLKPSCLFRQLPKFQDGDLTLYQSNAILRHLGRSFGLYGEDEEEAALVDMVNDGVEDLRCKYATLIY
360      370       380       390       400       410       420       430       440
TDYEAGKEEYVEELPEHLKPFETLLSENEGGEAFVVGSEISFADYNLLDLLRIHQVLNPSCLDAFPLLSAYVARLSARPEIEAFLASPEH
450      460
VDRPINGNGKQ*
```

**Figure S26.** Amino acid sequence of GFP-GST-40 fusion protein. GFP and GST-40 amino acids are highlighted in yellow and blue, respectively.

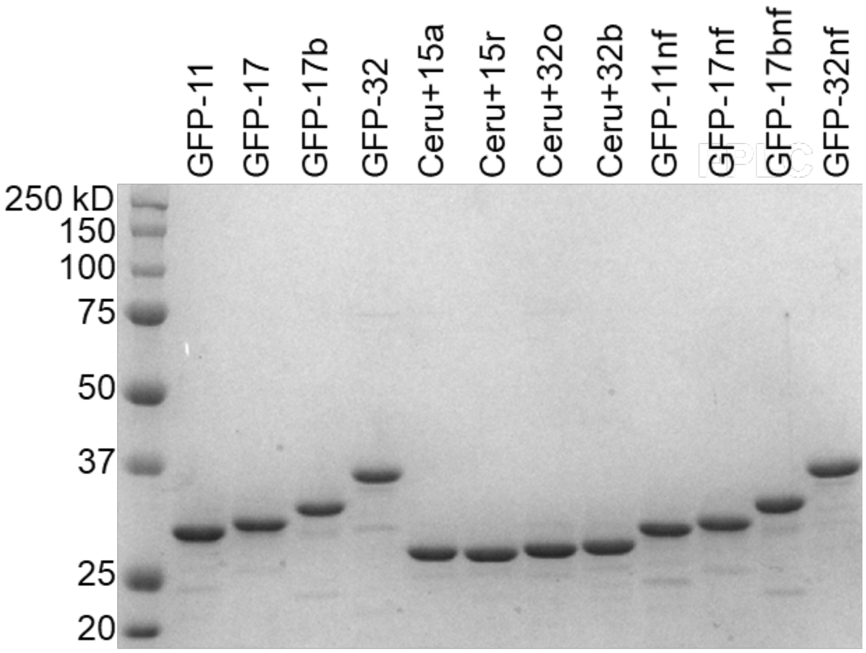


**Figure S27.** SDS-PAGE denaturing gel of the supercharged proteins used here. Approximately 3 μg of each protein in PBS were run on each lane, and proteins were stained with Coomassie blue dye.

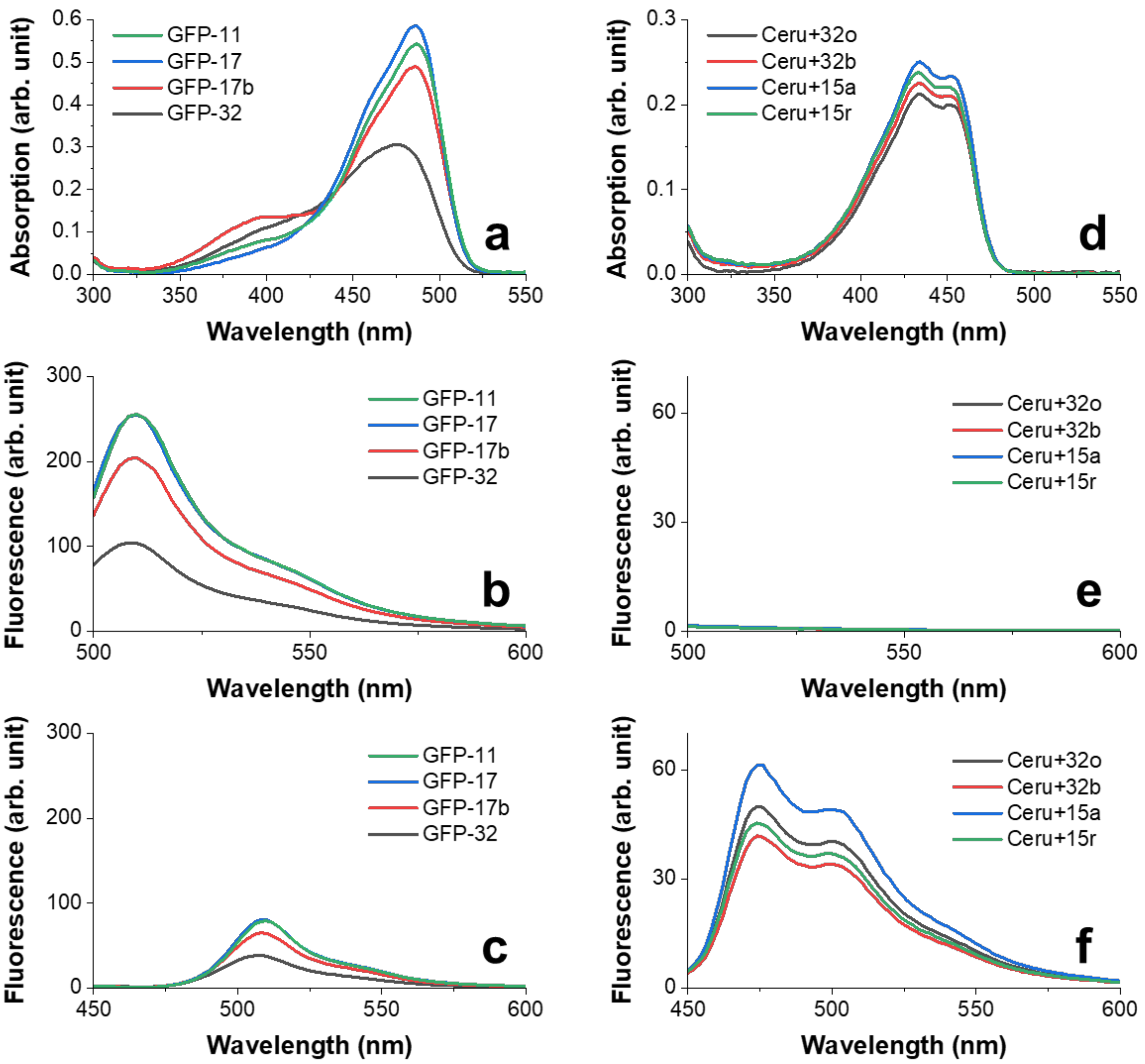


**Figure S28.** (a) UV-vis absorption spectra of each negatively charged GFP variant. Fluorescence emission spectra at 485 nm (b) and 433 nm (c) excitation for each negatively charged GFP variant. d) UV-vis absorption spectra of each positively charged Ceru variant. Fluorescence emission spectra at 485 nm (e) and 433 nm (f) excitation for each positively charged Ceru variant. The concentration of each protein is approximately 0.2 mg/mL in 50 mM Tris buffer, pH 7.4.

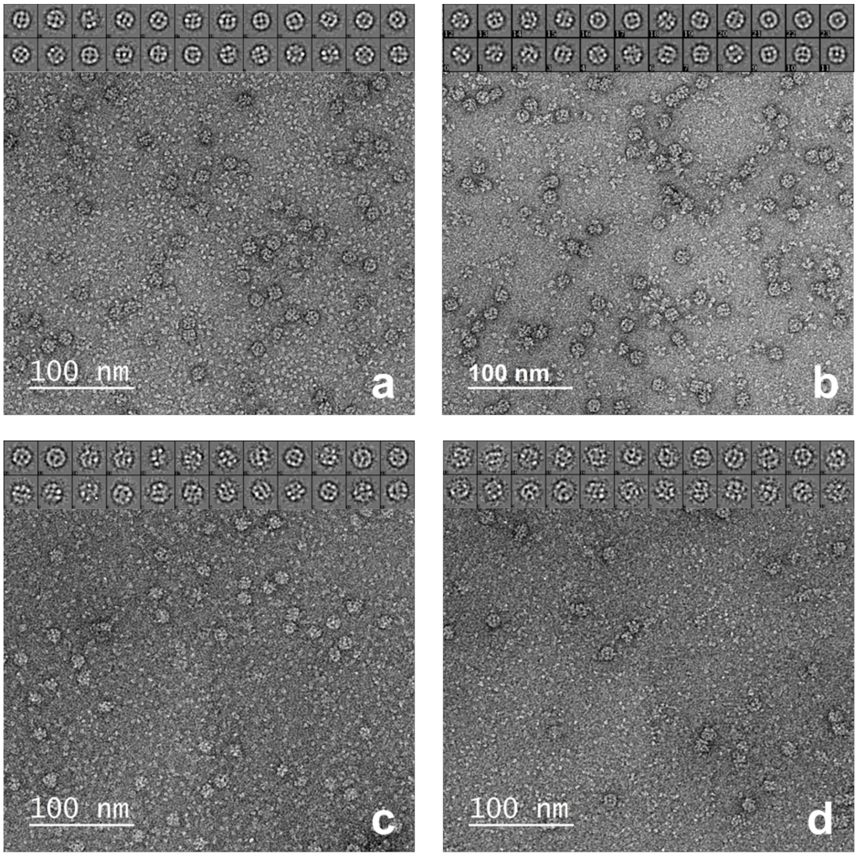


**Figure S29.** Representative negative stain TEM images and particle class averages for assembly between Ceru+32o / GFP-17 at 75 mM NaCl with the following ratios of positively to negatively charged subunits: (a) 2:1 (*n* = 1,203 particle); (b) 1:1 (*n* = 1,222 particles); (c) 1:2 (*n* = 836 particles); and (d) 1:4 (*n* = 505 particles). The protomer structure remains constant at all ratios.

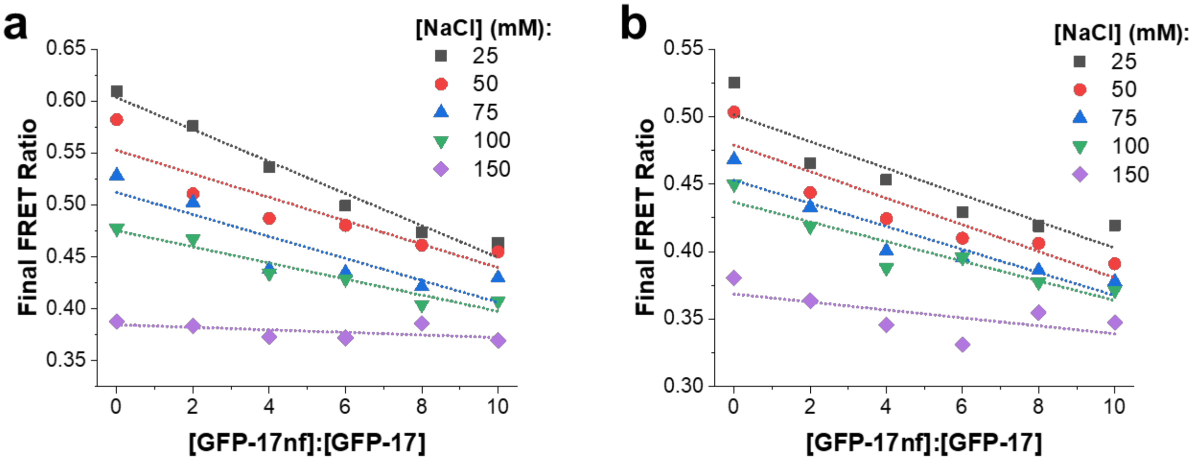


**Figure S30.** Experimentally measured final FRET ratios with different concentrations of non-fluorescent analog at different solution conditions for (a) Ceru+32o / GFP-17 and (b) Ceru+32b / GFP-17. The linear relationship between final FRET ratio and non-fluorescent analog concentration suggests FRET ratio is proportional to fraction of non-fluorescent analog in the assembled structure.

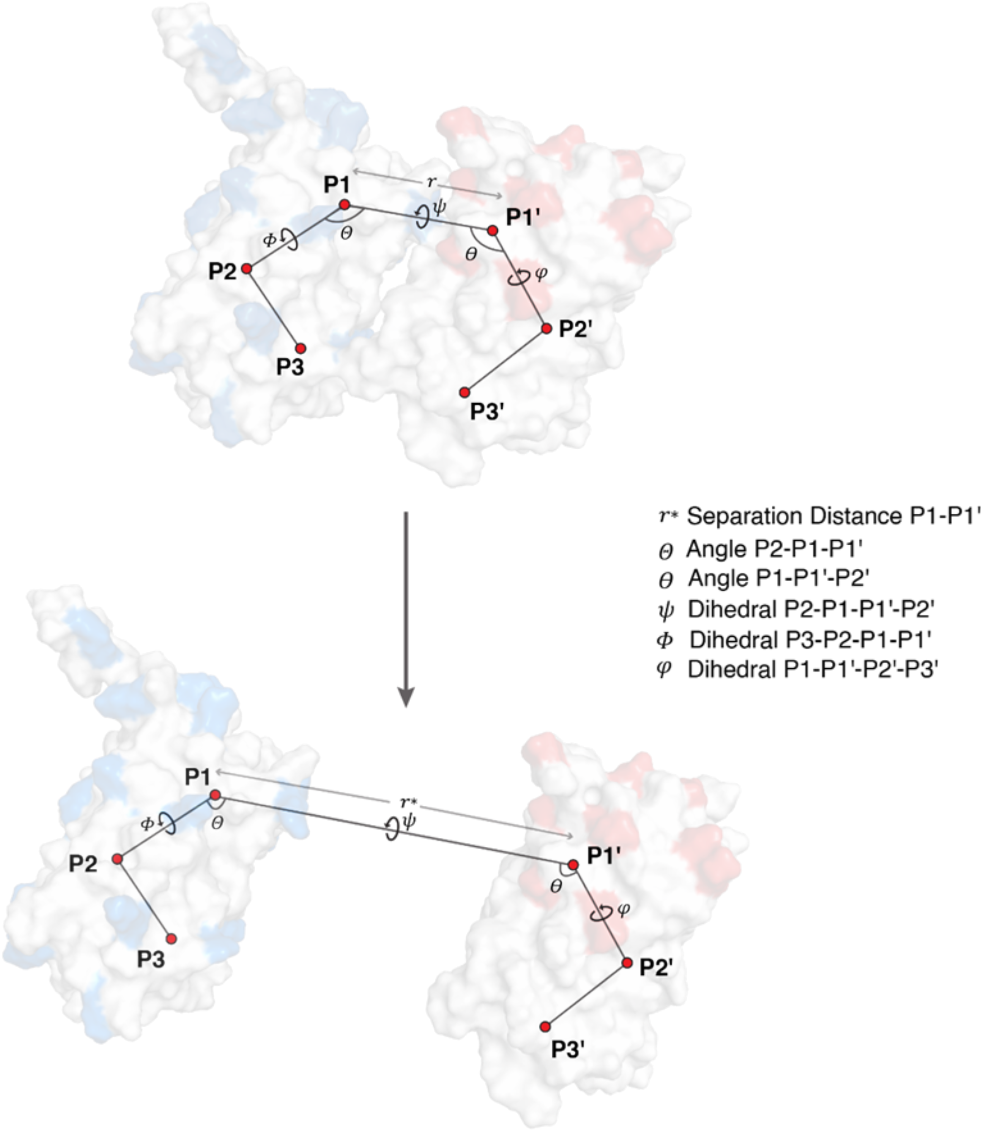


**Figure S31.** Constraints used in separating the two proteins. (top) Proteins in the bound state (site). (bottom) Proteins in the unbound state (bulk). Orientational (**Θ**, **Φ**,**Ψ**) and positional (**θ**,**φ**) constraints are applied, as separation distance ***r*** is increased till ***r**.**

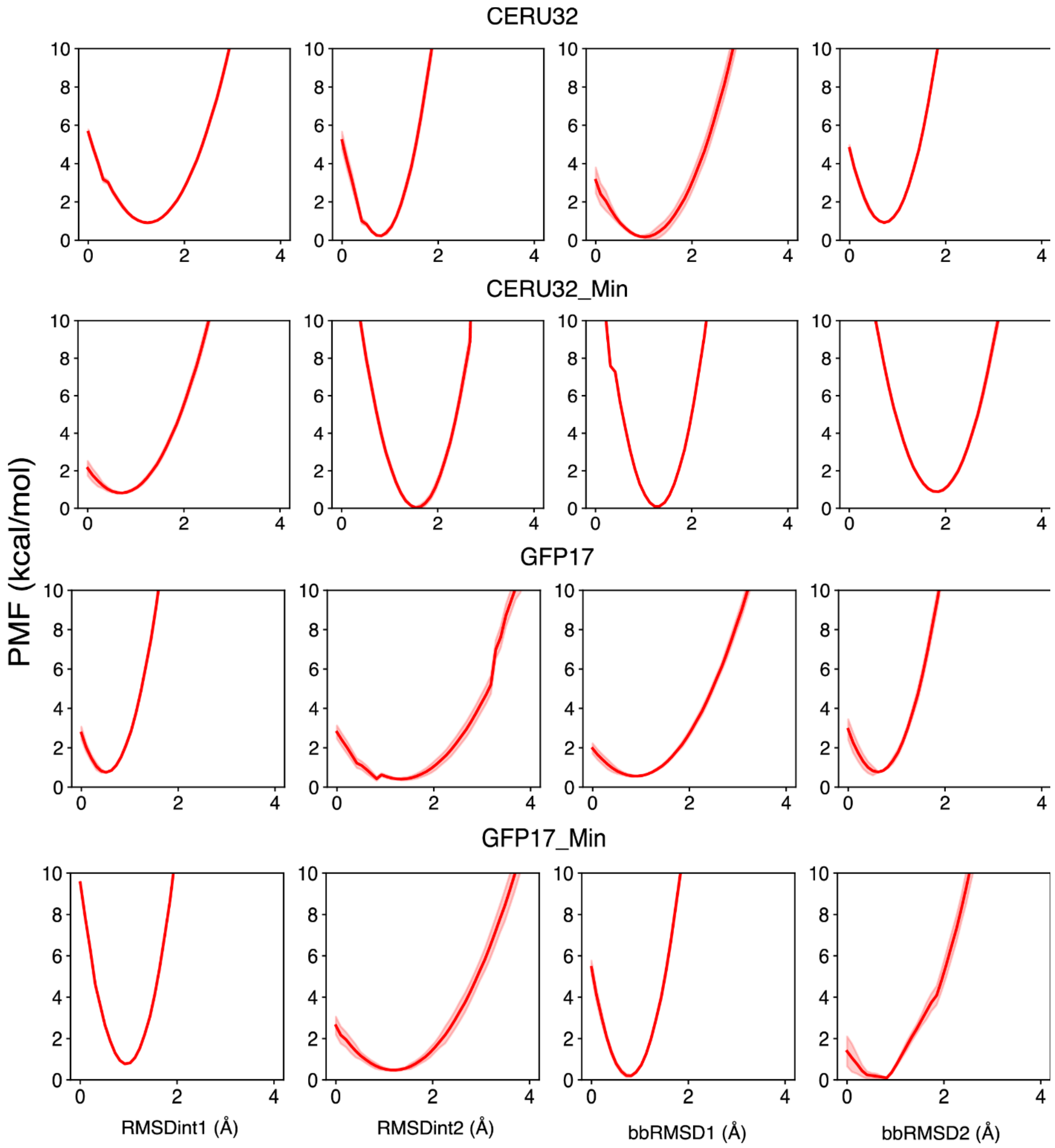


**Figure S32:** PMFs of the intra-planar systems' contributions of the conformational constraints in the unbound state (bulk).

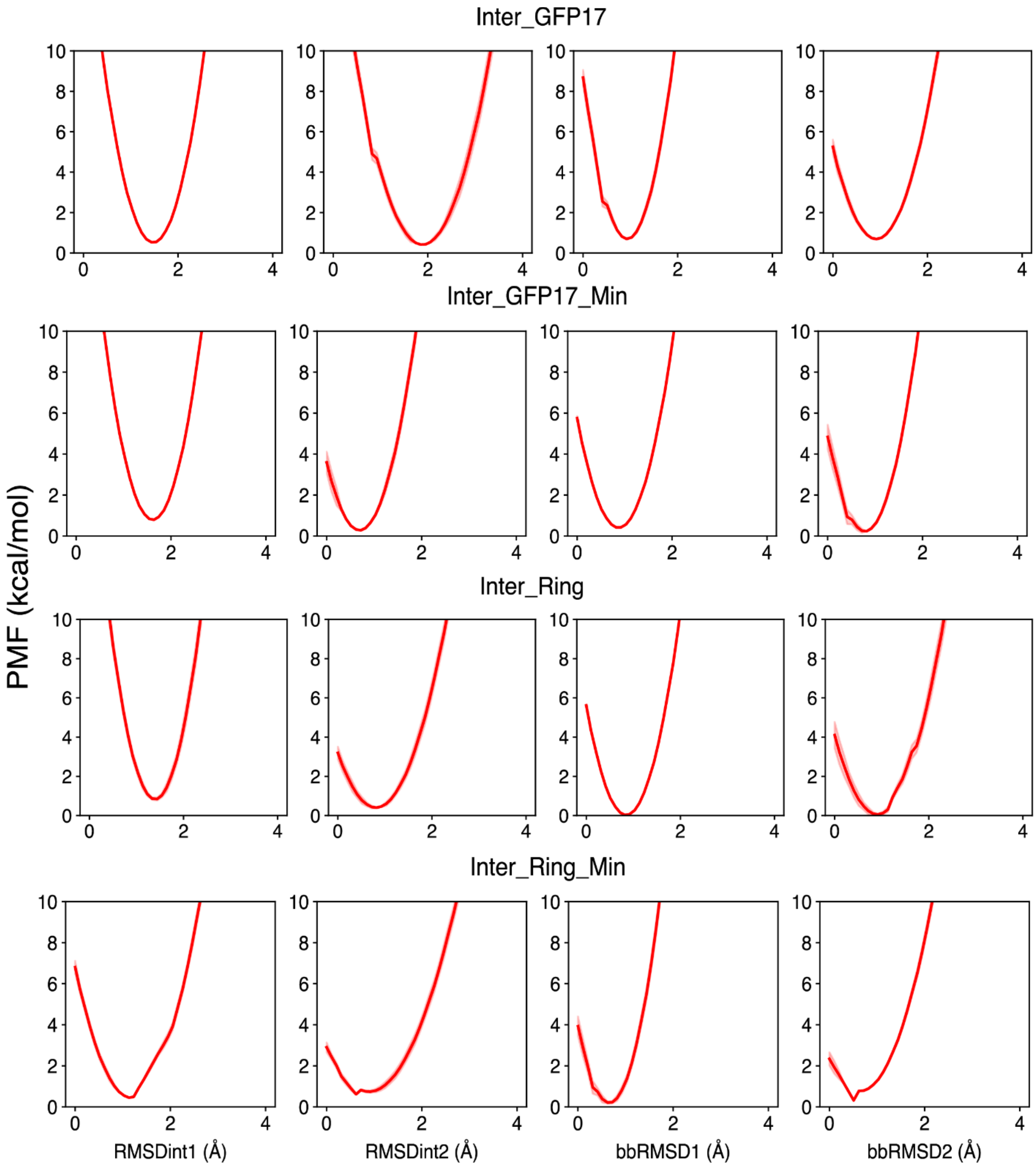


**Figure S33.** PMFs of the inter-planar systems' contributions of the conformational constraints in the unbound state (bulk).

**Table S1:** Contributing terms to $\Delta G^0_{bind}$ with errors for 'Ceru+32 Clockwise' for fully charged and minimally mutated proteins.

| Contributions | Fully charged | | Minimally mutated | | Unit | State |
|---|---|---|---|---|---|---|
| | | ± | | ± | | |
| $G_{RMSD_{int1}}$ | 3.15 | 0.1171 | 4.754 | 0.1015 | kcal/mol | Bound |
| $G_{RMSD_{int2}}$ | 12.12 | 0.3004 | 4.227 | 0.1055 | kcal/mol | Bound |
| $G_{RMSD_{bb1}}$ | 4.188 | 0.1364 | 5.089 | 0.0512 | kcal/mol | Bound |
| $G_{RMSD_{bb2}}$ | 16.57 | 0.3218 | 3.379 | 0.2039 | kcal/mol | Bound |
| $G_{\phi}$ | 1.778 | 0.0363 | 1.125 | 0.0322 | kcal/mol | Bound |
| $G_{\theta}$ | 2.417 | 0.0962 | 1.574 | 0.0509 | kcal/mol | Bound |
| $G_{\psi}$ | 2.862 | 0.064 | 3.201 | 0.1051 | kcal/mol | Bound |
| $G_{\Phi}$ | 1.744 | 0.0289 | 5.205 | 0.1829 | kcal/mol | Bound |
| $G_{\Theta}$ | 3.143 | 0.0509 | 3.273 | 0.0411 | kcal/mol | Bound |
| $G_{RMSD_{int1}}$ | 3.097 | 0.0626 | 1.214 | 0.0375 | kcal/mol | Unbound |
| $G_{RMSD_{int2}}$ | 1.989 | 0.0241 | 7.37 | 0.1566 | kcal/mol | Unbound |
| $G_{RMSD_{bb1}}$ | 2.174 | 0.0994 | 5.495 | 0.0982 | kcal/mol | Unbound |
| $G_{RMSD_{bb2}}$ | 1.718 | 0.0378 | 8.932 | 0.2357 | kcal/mol | Unbound |
| $G_0$ | 8.053 | 0.3479 | 8.053 | 0.279 | kcal/mol | Unbound |
| $S^*$ | 1.868 | 0.09391 | 4.779 | 0.2841 | Å$^2$ | |
| $I^*$ | 1.393E+44 | 4.310E+42 | 1.81E+37 | 1.140E+36 | Å | |
| $G_{bind}$ | 83.44 | 3.5503 | 56 | 2.6823 | kcal/mol | |

**Table S2:** Contributing terms to $\Delta G^0_{bind}$ with errors for 'GFP-17 Clockwise' for fully charged and minimally mutated proteins.

| Contributions | Fully charged | | Minimally mutated | | Unit | State |
|---|---|---|---|---|---|---|
| | | ± | | ± | | |
| $G_{RMSD_{int1}}$ | 7.608 | 0.1558 | 1.538 | 0.0295 | kcal/mol | Bound |
| $G_{RMSD_{int2}}$ | 1.547 | 0.0651 | 9.627 | 0.1729 | kcal/mol | Bound |
| $G_{RMSD_{bb1}}$ | 13.22 | 0.1677 | 3.159 | 0.0939 | kcal/mol | Bound |
| $G_{RMSD_{bb2}}$ | 3.558 | 0.0756 | 9.116 | 0.2734 | kcal/mol | Bound |
| $G_{\phi}$ | 1.686 | 0.0638 | 1.399 | 0.0367 | kcal/mol | Bound |
| $G_{\theta}$ | 1.84 | 0.0403 | 2.542 | 0.2364 | kcal/mol | Bound |
| $G_{\psi}$ | 1.493 | 0.0485 | 2.779 | 0.166 | kcal/mol | Bound |
| $G_{\Phi}$ | 1.921 | 0.0236 | 2.131 | 0.104 | kcal/mol | Bound |
| $G_{\Theta}$ | 3.23 | 0.0572 | 0.7506 | 0.0134 | kcal/mol | Bound |
| $G_{RMSD_{int1}}$ | 0.9392 | 0.0399 | 3.117 | 0.1656 | kcal/mol | Unbound |
| $G_{RMSD_{int2}}$ | 2.262 | 0.082 | 2.127 | 0.1086 | kcal/mol | Unbound |
| $G_{RMSD_{bb1}}$ | 1.516 | 0.0395 | 2.049 | 0.0282 | kcal/mol | Unbound |
| $G_{RMSD_{bb2}}$ | 1.203 | 0.0421 | 1.107 | 0.0693 | kcal/mol | Unbound |
| $G_0$ | 8.05 | 0.4155 | 8.049 | 0.5295 | kcal/mol | Unbound |
| $S^*$ | 4.779 | 0.1085 | 4.779 | 0.2536 | Å$^2$ | |
| $I^*$ | 2.02E+28 | 1.08E+27 | 1.46E+19 | 5.94E+17 | Å | |
| $G_{bind}$ | 52.42 | 1.489 | 33.88 | 1.741 | kcal/mol | |

**Table S3:** Contributing terms to $\Delta G^0_{bind}$ with errors for 'Inter-GFP-17' for fully charged and minimally mutated proteins.

| Contributions | Fully charged | | Minimally mutated | | Unit | State |
|---|---|---|---|---|---|---|
| | | ± | | ± | | |
| $G_{RMSD_{int1}}$ | 2.557 | 0.0683 | 2.291 | 0.1081 | kcal/mol | Bound |
| $G_{RMSD_{int2}}$ | 9.282 | 0.4979 | 1.503 | 0.0366 | kcal/mol | Bound |
| $G_{RMSD_{bb1}}$ | 5.088 | 0.1705 | 1.48 | 0.0383 | kcal/mol | Bound |
| $G_{RMSD_{bb2}}$ | 3.756 | 0.082 | 4.517 | 0.0862 | kcal/mol | Bound |
| $G_{\phi}$ | 1.873 | 0.0529 | 1.294 | 0.0478 | kcal/mol | Bound |
| $G_{\theta}$ | 2.737 | 0.056 | 1.834 | 0.0495 | kcal/mol | Bound |
| $G_{\psi}$ | 1.449 | 0.0286 | 2.855 | 0.0449 | kcal/mol | Bound |
| $G_{\Phi}$ | 2.279 | 0.1416 | 1.102 | 0.0821 | kcal/mol | Bound |
| $G_{\Theta}$ | 2.308 | 0.037 | 2.022 | 0.0354 | kcal/mol | Bound |
| $G_{RMSD_{int1}}$ | 6.881 | 0.1037 | 8.407 | 0.2019 | kcal/mol | Unbound |
| $G_{RMSD_{int2}}$ | 8.521 | 0.257 | 1.562 | 0.0492 | kcal/mol | Unbound |
| $G_{RMSD_{bb1}}$ | 2.874 | 0.0827 | 2.385 | 0.0612 | kcal/mol | Unbound |
| $G_{RMSD_{bb2}}$ | 2.382 | 0.0343 | 1.948 | 0.0664 | kcal/mol | Unbound |
| $G_0$ | 8.451 | 0.2326 | 8.496 | 0.5561 | kcal/mol | Unbound |
| $S^*$ | 4.779 | 0.1277 | 1.531 | 0.07416 | Å$^2$ | |
| $I^*$ | 3.1E+12 | 7.953E+10 | 1.275E+08 | 3.538E+06 | Å | |
| $G_{bind}$ | 13.36 | 0.3943 | 7.116 | 0.6567 | kcal/mol | |

**Table S4:** Contributing terms to $\Delta G^0_{bind}$ with errors for 'Inter-Ring' for fully charged and minimally mutated proteins.

| Contributions | Fully charged | | Minimally mutated | | Unit | State |
|---|---|---|---|---|---|---|
| | | ± | | ± | | |
| $G_{RMSD_{int1}}$ | 7.01 | 0.1916 | 2.814 | 0.2114 | kcal/mol | Bound |
| $G_{RMSD_{int2}}$ | 1.592 | 0.0171 | 1.524 | 0.0504 | kcal/mol | Bound |
| $G_{RMSD_{bb1}}$ | 2.757 | 0.0586 | 6.082 | 0.2373 | kcal/mol | Bound |
| $G_{RMSD_{bb2}}$ | 9.316 | 0.1582 | 13.17 | 0.147 | kcal/mol | Bound |
| $G_{\phi}$ | 1.1 | 0.0175 | 0.9923 | 0.0193 | kcal/mol | Bound |
| $G_{\theta}$ | 2.519 | 0.099 | 0.8688 | 0.0167 | kcal/mol | Bound |
| $G_{\psi}$ | 2.884 | 0.0607 | 2.194 | 0.076 | kcal/mol | Bound |
| $G_{\Phi}$ | 1.044 | 0.0235 | 2.336 | 0.0462 | kcal/mol | Bound |
| $G_{\Theta}$ | 2.338 | 0.0441 | 1.673 | 0.033 | kcal/mol | Bound |
| $G_{RMSD_{int1}}$ | 6.584 | 0.1627 | 3.401 | 0.0554 | kcal/mol | Unbound |
| $G_{RMSD_{int2}}$ | 1.751 | 0.0307 | 1.665 | 0.0528 | kcal/mol | Unbound |
| $G_{RMSD_{bb1}}$ | 2.333 | 0.0439 | 1.433 | 0.0507 | kcal/mol | Unbound |
| $G_{RMSD_{bb2}}$ | 2.212 | 0.0839 | 1.183 | 0.0251 | kcal/mol | Unbound |
| $G_0$ | 8.496 | 0.3708 | 8.496 | 0.5214 | kcal/mol | Unbound |
| $S^*$ | 5.056 | 0.1999 | 2.351 | 0.09781 | Å$^2$ | |
| $I^*$ | 9.27E+09 | 5.97E+08 | 1.554E+07 | 8.401E+05 | Å | |
| $G_{bind}$ | 7.465 | 0.556 | 5.61 | 0.5743 | kcal/mol | |

**Table S5.** Measured critical NaCl concentrations and calculated net charge for each oppositely supercharged protein combination.

| Combination | pH | Pos. Charge | Neg. Charge | $[NaCl]_{crit}$ (mM) |
|---|---|---|---|---|
| Ceru+15a / GFP-11 | 7.4 | 16.7 | -10 | --- |
| Ceru+15a / GFP-17b | 7.4 | 16.7 | -16.8 | 63±11 |
| Ceru+15a / GFP-17 | 7.4 | 16.7 | -14.8 | --- |
| Ceru+15a / GFP-32 | 7.4 | 16.7 | -28.7 | 97±13 |
| Ceru+15r / GFP-11 | 7.4 | 15.8 | -10 | --- |
| Ceru+15r / GFP-17b | 7.4 | 15.8 | -16.8 | 127±27 |
| Ceru+15r / GFP-17 | 7.4 | 15.8 | -14.8 | 47±11 |
| Ceru+15r / GFP-32 | 7.4 | 15.8 | -28.7 | 147±25 |
| Ceru+32b / GFP-11 | 7.4 | 31.5 | -10 | 143±45 |
| Ceru+32b / GFP-17b | 7.4 | 31.5 | -16.8 | 277±61 |
| Ceru+32b / GFP-17 | 7.4 | 31.5 | -14.8 | 243±85 |
| Ceru+32b / GFP-32 | 7.4 | 31.5 | -28.7 | 398±77 |
| Ceru+32o / GFP-11 | 7.4 | 30.7 | -10 | 118±40 |
| Ceru+32o / GFP-17b | 7.4 | 30.7 | -16.8 | 245±43 |
| Ceru+32o / GFP-17 | 7.4 | 30.7 | -14.8 | 234±64 |
| Ceru+32o / GFP-32 | 7.4 | 30.7 | -28.7 | 366±57 |
| Ceru+15a / GFP-11 | 6.0 | 20.9 | -6.4 | 8±2 |
| Ceru+15a / GFP-17b | 6.0 | 20.9 | -12.5 | 95±14 |
| Ceru+15a / GFP-17 | 6.0 | 20.9 | -10.8 | 46±11 |
| Ceru+15a / GFP-32 | 6.0 | 20.9 | -23.8 | 150±16 |
| Ceru+15r / GFP-11 | 6.0 | 18.8 | -6.4 | 29±8 |
| Ceru+15r / GFP-17b | 6.0 | 18.8 | -12.5 | 128±25 |
| Ceru+15r / GFP-17 | 6.0 | 18.8 | -10.8 | 59±13 |
| Ceru+15r / GFP-32 | 6.0 | 18.8 | -23.8 | 183±25 |
| Ceru+32b / GFP-11 | 6.0 | 34.6 | -6.4 | 99±30 |
| Ceru+32b / GFP-17b | 6.0 | 34.6 | -12.5 | 258±52 |
| Ceru+32b / GFP-17 | 6.0 | 34.6 | -10.8 | 153±35 |
| Ceru+32b / GFP-32 | 6.0 | 34.6 | -23.8 | 320±44 |
| Ceru+32o / GFP-11 | 6.0 | 34.4 | -6.4 | 98±26 |
| Ceru+32o / GFP-17b | 6.0 | 34.4 | -12.5 | 236±35 |
| Ceru+32o / GFP-17 | 6.0 | 34.4 | -10.8 | 133±28 |
| Ceru+32o / GFP-32 | 6.0 | 34.4 | -23.8 | 288±36 |
| Ceru+15a / GFP-11 | 9.0 | 13.4 | -13.9 | 19±4 |
| Ceru+15a / GFP-17b | 9.0 | 13.4 | -20.6 | 60±10 |
| Ceru+15a / GFP-17 | 9.0 | 13.4 | -18.6 | 32±7 |
| Ceru+15a / GFP-32 | 9.0 | 13.4 | -32.4 | 76±11 |
| Ceru+15r / GFP-11 | 9.0 | 11.1 | -13.9 | 45±18 |
| Ceru+15r / GFP-17b | 9.0 | 11.1 | -20.6 | 113±30 |
| Ceru+15r / GFP-17 | 9.0 | 11.1 | -18.6 | 82±19 |
| Ceru+15r / GFP-32 | 9.0 | 11.1 | -32.4 | 121±20 |
| Ceru+32b / GFP-11 | 9.0 | 25.7 | -13.9 | 129±52 |
| Ceru+32b / GFP-17b | 9.0 | 25.7 | -20.6 | 241±64 |
| Ceru+32b / GFP-17 | 9.0 | 25.7 | -18.6 | 257±99 |
| Ceru+32b / GFP-32 | 9.0 | 25.7 | -32.4 | 403±100 |
| Ceru+32o / GFP-11 | 9.0 | 25.3 | -13.9 | 137±37 |
| Ceru+32o / GFP-17b | 9.0 | 25.3 | -20.6 | 230±49 |
| Ceru+32o / GFP-17 | 9.0 | 25.3 | -18.6 | 234±67 |
| Ceru+32o / GFP-32 | 9.0 | 25.3 | -32.4 | 348±58 |
| Ceru+15a / GFP-GST-40 | 7.4 | 16.7 | -39.0 | 74±14 |
| Ceru+15r / GFP-GST-40 | 7.4 | 15.8 | -39.0 | 77±24 |
| Ceru+32b / GFP-GST-40 | 7.4 | 31.5 | -39.0 | 371±112 |
| Ceru+32o / GFP-GST-40 | 7.4 | 30.7 | -39.0 | 315±65 |

**Table S6.** Dynamic subunit exchange reaction model for a protomer containing 8 exchangeable subunits. The protomers initially contain fluorescent subunits (G) and exchange with non-fluorescent subunits (A).

| **Reactions:** | **Rate Constants:** |
|---|---|
| G8A0==>G7A0+G | $k_d$ |
| G7A0+G==>G8 | $k_a$ |
| G7A0+A==>G7A | $k_a$ |
| G7A1==>G6A1+G | (7/8)*$k_d$ |
| G7A1==>G7A0+A | (1/8)*$k_d$ |
| G6A1+G==>G7A1 | $k_a$ |
| G6A1+A==>G6A2 | $k_a$ |
| G6A2==>G5A2+G | (6/8)*$k_d$ |
| G6A2==>G6A1+A | (2/8)*$k_d$ |
| G5A2+G==>G6A2 | $k_a$ |
| G5A2+A==>G5A3 | $k_a$ |
| G5A3==>G4A3+G | (5/8)*$k_d$ |
| G5A3==>G5A2+A | (3/8)*$k_d$ |
| G4A3+A==>G4A4 | $k_a$ |
| G4A3+G==>G5A3 | $k_a$ |
| G4A4==>G3A4+G | (4/8)*$k_d$ |
| G4A4==>G4A3+A | (4/8)*$k_d$ |
| G3A4+A==>G3A5 | $k_a$ |
| G3A4+G==>G4A4 | $k_a$ |
| G3A5==>G2A5+G | (3/8)*$k_d$ |
| G3A5==>G3A4+A | (5/8)*$k_d$ |
| G2A5+A==>G2A6 | $k_a$ |
| G2A5+G==>G3A5 | $k_a$ |
| G2A6==>G1A6+G | (2/8)*$k_d$ |
| G2A6==>G2A5+A | (6/8)*$k_d$ |
| G1A6+G==>G2A6 | $k_a$ |
| G1A6+A==>G1A7 | $k_a$ |
| G1A7==>G10A7+G | (1/8)*$k_d$ |
| G1A7==>G1A6+A | (7/8)*$k_d$ |
| G0A7+G==>G1A7 | $k_a$ |
| G0A7+A==>G0A8 | $k_a$ |
| G0A8==>G0A7+A | $k_d$ |

**Table S7.** List of Modelled Residues:

| Ceru32Clockwise: |
| --- |
| R234 |
| D235 |
| E236 |
| R237 |
| Y238 |
| K239 |
| L240 |
| E241 |

**Table S8.** Mutations performed to construct the minimal mutants.

| Ceru32Clockwise → Ceru_minClockwise | GFP17Clockwise → GFP_minClockwise |
| --- | --- |
| R7E | D43T |
| R10T | E80R |
| K12V | E149N |
| K20D | E198N |
| K33E | D227A |
| K35E | E231H |
| K77D | |
| K91E | |
| K103D | |
| R118D | |
| R129I | |
| K134D | |
| R143E | |
| R158Q | |
| K173E | |
| R198D | |
| R234M | |
| R237L | |
| L240G | |
| E241S | |

**Table S9.** General System information regarding each Simulation System.

| System | n(Atoms) | Box Dimensions (ÅxÅxÅ) | n($Na^+$/$Cl^-$) | n(Waters) |
|---|---|---|---|---|
| Ceru32Clockwise | 119,570 | 124x103x94 | 84/97 | 37,264 |
| Ceru_minClockwise | 122,052 | 124x103x96 | 93/85 | 38,144 |
| GFP17Clockwise | 115,424 | 124x102x92 | 81/94 | 35,884 |
| GFP_minClockwise | 114,156 | 124x101x91 | 88/81 | 35,515 |
| InterGFP17 | 102,383 | 123x92x91 | 109/73 | 31,659 |
| InterGFP_min | 102,382 | 123x92x91 | 91/73 | 31,652 |
| Inter_Ring | 138,166 | 138x123x94 | 97/110 | 43,454 |
| Inter_Ring_min | 132,475 | 130x114x89 | 101/93 | 41,613 |
